# Building virtual cell dynamics that evolve with biological measurements

**DOI:** 10.64898/2026.10.23.761350

**Authors:** Zhenyi Zhang, Zihan Wang, Qinxue Tian, Baichuan Zhou, Tianyu Zhou, Qing Nie, Peijie Zhou

## Abstract

Virtual cell models built from single-cell measurements need to predict how cells change over time, but new biological measurements often reveal dynamical relationships that existing models cannot express. Translating such measurements into dynamics therefore requires deciding what biological relation should be modeled, where it should enter the formulation and how its recovery should be validated. Here we present CellCompass, an agentic AI system that couples a scientific agent with an extensible dynamical engine for modeling state transitions and population change, along with fate and molecular readout. CellCompass reuses existing formulations when sufficient and designs, implements and refines new ones when experimental measurements introduce additional constraints. To assess this capability, we developed DynBench100, a benchmark of 100 simulated single-cell time-course scenarios with hidden dynamical ground truth. CellCompass achieved the highest aggregate performance among the tested general-purpose and biomedical agents across six dynamical tasks. Across four real single-cell studies, it retained an existing velocity-growth model to reconstruct context-dependent epithelial-mesenchymal transition and developed three new dynamical formulations for distinct experimental measurements. In drug-response data, incorporating metabolic RNA labeling reduced pulse-control identifiability error by 31.1%, revealing staged response programs and generating testable predictions for perturbing a candidate adaptive program. In hematopoietic lineage tracing, incorporating clonal barcodes improved clone-fate agreement by 54% over the strongest dynamical baseline, identifying early lineage priming among transcriptionally similar progenitors. Finally, in time-resolved CITE-seq data, CellCompass reduced future surface-protein distribution error by 13.2% for RNA-initialized trajectories, uncovering lineage-specific temporal offsets between transcriptional and protein programs. Together, CellCompass provides a framework in which virtual-cell dynamics can expand with the biological measurements used to constrain them.

## Introduction

Predicting how cells behave over time is a central goal of biology. A useful virtual cell should predict how cells respond to genetic and pharmacological perturbations, progress through development and disease, and adapt to changing environments^1–5^. Recent advances in single-cell measurement and artificial intelligence have made this goal increasingly plausible by capturing cellular state, history, response and molecular context at large scale. While cellular foundation models organize millions of profiles into unified state representations^6–13^, building a predictive virtual cell requires moving beyond static representation toward dynamical models that simulate how cellular states, population abundances, and molecular phenotypes evolve over time and under intervention^2,14–16^.

What these models can predict depends on the biological relationships revealed by experiments. For instance, time-resolved measurements constrain how cellular populations move through molecular state space, whereas cell counts and population composition reveal which states expand, persist or disappear^17,18^. Perturbation experiments show how genetic, chemical or environmental interventions redirect cellular responses^19–21^, and lineage measurements connect early cells to their descendants, separating present molecular similarity from shared developmental potential^22–25^. Metabolic labeling reveals molecular production and persistence, while paired and multimodal measurements show how regulatory changes propagate across molecular layers and become phenotype^26–28^. These measurements are particularly important when different biological processes produce similar observed states. For example, transcriptionally similar progenitors may have different lineage potentials, and changes in cancer-cell populations may arise from state transitions, selective expansion or cell loss. Incorporating the relevant measurements into dynamical models is therefore essential for distinguishing these processes and predicting their subsequent behavior.

A range of mathematical approaches is available for modeling cellular dynamics, including pseudotime and RNA velocity to infer progression from molecular state^28–37^, optimal transport and Schrödinger bridges to couple populations across destructive time points, with extensions that incorporate spatial information to reconstruct cellular organization and dynamics across time and space^38–47^, unbalanced formulations to represent changes in effective abundance^48–51^, and flow-based methods to generate continuous trajectories^52–57^. These approaches differ in how they represent state transitions, stochasticity, population growth and fate branching. Their assumptions give dynamical models mechanistic structure and determine the biological questions each model can answer^58^. Recent models have advanced the prediction of cellular responses to perturbations across biological contexts and over time^59–62^. These advances raise the question of how a model’s formulation should change when new measurements reveal biological relationships that it does not yet represent. As new measurement technologies and experimental designs reveal additional aspects of cellular behavior^63–65^, existing formulations may need to be extended to incorporate the information they provide. This requires identifying what the measurements constrain, determining how that information enters the governing dynamics, and validating whether the resulting model recovers the intended biological relationship. Model design remains an expert-intensive process, often repeated independently for each study. Scientific agents now help generate hypotheses, analyze data, develop scientific software and integrate evidence across biomedical and therapeutic discovery workflows^66–74^. Extending these capabilities to virtual-cell modeling requires designing and validating dynamical formulations that match the measurements at hand.

Here we introduce CellCompass, an agentic AI system that selects, constructs and validates dynamical models according to the biological relationships revealed by each experiment (Fig. 1). CellCompass combines (1) a scientific agent with (2) an extensible dynamical engine built on modular transport, bridge and flow-matching formulations. Within this architecture, the scientific agent maps measurement-specific constraints onto the engine’s modular coupling, path and readout components and validates candidates against measurement-derived biological criteria, reusing existing formulations when sufficient and retaining validated new formulations in the engine as new experimental modalities are encountered.

**Figure 1:**
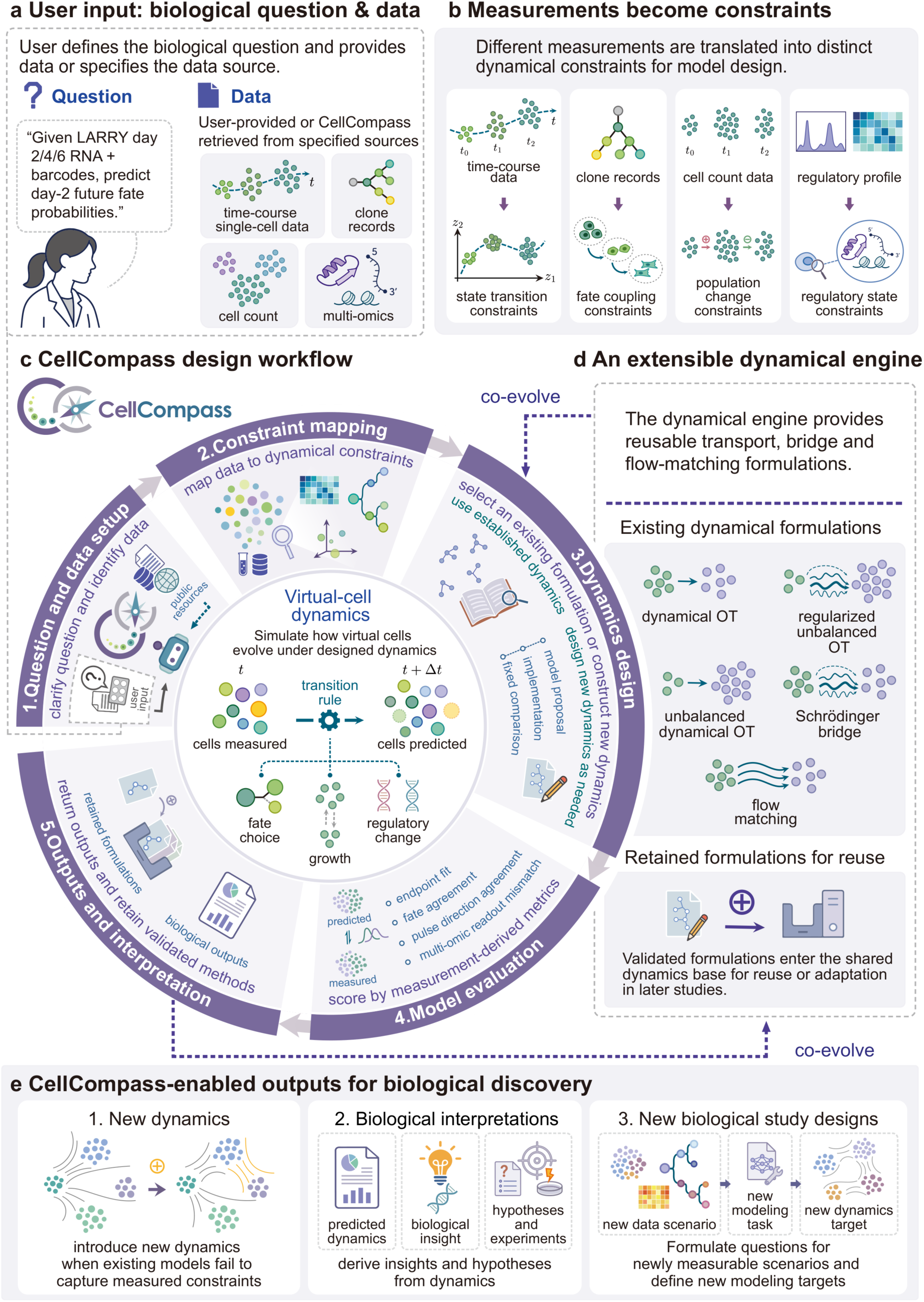
CellCompass maps biological questions and single-cell measurements to co-evolving virtual-cell dynamics. **a,** A study begins with a biological question and the available data. The example asks CellCompass to predict future fate probabilities for day-2 progenitors using LARRY day-2, day-4 and day-6 RNA profiles and lineage barcodes. Inputs may include time-course single-cell data, clone records, cell-count measurements and multi-omic profiles supplied by the user or retrieved from specified sources. b, Different measurements are translated into distinct dynamical constraints: time-course data constrain state transitions, clone records connect early cells to later fates, cell-count data inform population change and regulatory or multi-omic profiles constrain changes in molecular state. **c,** CellCompass carries the study through five linked stages that define the question and data, map measurements to dynamical constraints, select or construct the dynamics, evaluate generated predictions and interpret the outputs. Candidate dynamics are compared using measurement-derived criteria, including endpoint fit, fate agreement, pulse-direction recovery and multi-omic readout agreement. At the center, cells observed at time t are propagated by a learned transition rule to predicted cells at t+Δt, with movement, fate choice, effective growth and regulatory change represented according to the biological question. **d,** The extensible dynamical engine supports dynamical optimal transport, regularized unbalanced optimal transport, unbalanced dynamical optimal transport, Schrödinger bridges and flow matching. Validated formulations are retained within the engine for reuse or adaptation, allowing the scientific agent and dynamical engine to co-evolve as new measurements and questions arise. **e,** CellCompass can produce new dynamics, biological interpretations and hypotheses, and study designs that translate newly measurable biological scenarios into new modeling questions and dynamical targets.

We evaluated CellCompass using DynBench100, our newly contributed benchmark of 100 simulated single-cell scenarios with hidden dynamical ground truth, and four real single-cell applications. On DynBench100, CellCompass achieved a higher aggregate score across six dynamical recovery tasks than general-purpose coding and biomedical agents. In the four experimental studies, it applied an existing velocity–growth model to context-dependent epithelial–mesenchymal transition (EMT) and constructed three new formulations (PulseBridge, LineageBridge and ProteinBridge) to incorporate metabolic RNA labeling, clonal barcodes and paired RNA–surface-protein measurements. These formulations improved predictions of drug-response direction, clonal fate and future surface-protein states.

## Results

### CellCompass designs virtual-cell dynamics that represent the biological relations revealed by new experiments

Single-cell snapshots alone often leave alternative dynamical explanations indistinguishable, making model formulation dependent on what an experiment actually measures (Fig. 1a,b). CellCompass therefore asks *which relation between present and future cells an experiment makes observable and what a dynamical model must preserve to recover it.* For instance, clone barcodes, metabolic labeling, cell counts and paired multi-omic readouts each expose different temporal relationships and call for distinct dynamical structures. CellCompass uses a *scientific agent* to identify which of these biological relations is relevant to the question and whether the available dynamics can represent it.

The scientific agent drives each study through five stages (Fig. 1c): (1) question and data setup to define the prediction target and inputs; (2) constraint mapping from measurements to dynamical constraints; (3) dynamics design to select an existing formulation or construct a new one; (4) model evaluation against measurement-derived criteria; and (5) outputs and interpretation using the validated trajectories. If a candidate fails the prespecified comparison, CellCompass returns to dynamics design, revises how the measurement constrains the model or modifies the formulation itself, and evaluates the revised model again (Supplementary Note S1).

The second component of CellCompass is an extensible *dynamical engine* that provides the transport, bridge and flow-matching families used for single-cell dynamics^48,49,53–56^ (Fig. 1d; Supplementary Notes S2 and S3). Within this modular engine, each model specifies how cells are coupled across time, how coupled cells are connected, and what additional quantities are carried along the path, such as mass, fate or molecular readouts. This structure allows experimental constraints to map directly onto model components. When an existing formulation already represents the required biology, CellCompass reuses it; otherwise, the scientific agent modifies or combines only the components needed to incorporate the new measurement. Validated formulations are retained in the engine for reuse or adaptation in later studies.

Because matching marginal snapshots alone does not guarantee faithful inter-timepoint trajectories, candidate models undergo staged validation against measurement-derived biological criteria before acceptance (Fig. 1c; Supplementary Fig. 1 and Methods). During this process, context-isolated reviewer agents examine whether the proposed relation is supported by the data, whether the implementation matches the formulation, and whether the planned comparison tests that relation. Each reviewer works in a separate conversation and receives a task brief, the relevant proposal or code, and numerical results. Candidates are compared with fixed reference methods and tuned to improve population agreement while retaining the proposed biological improvement. Models that pass these checks support three types of downstream output: new dynamical formulations, biological interpretations and experimentally testable study designs (Fig. 1e).

### DynBench100 evaluates CellCompass across six dynamical tasks

To evaluate whether CellCompass can recover cellular dynamics under controlled ground truth, we developed DynBench100, a benchmark of simulated single-cell time courses with hidden dynamical structure. Each scenario was simulated from a regulatory system with a defined fate structure, a growth or death process and a defined response to perturbation, and agents received only the resulting time course, a task specification and a common terminal-state classifier (Fig. 2a; Methods). They were then asked to recover six quantities that are known to the simulator but not directly observed in the data.

**Figure 2:**
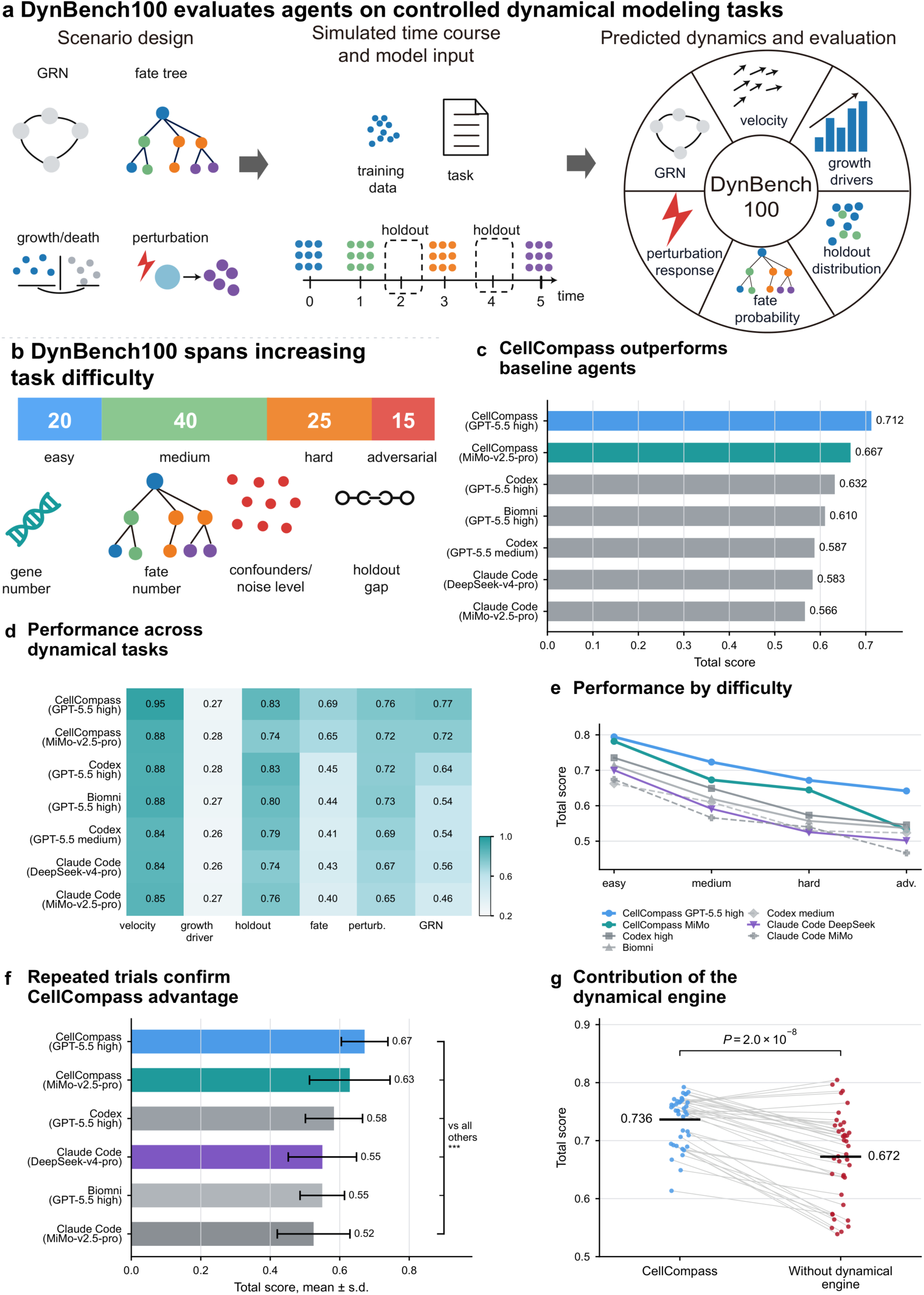
CellCompass performance across six DynBench100 tasks. **a,** DynBench100 converts simulated single-cell dynamical scenarios with hidden ground truth into benchmark tasks. Agents receive time-course data, task specifications and a common terminal-state classifier, then recover velocity fields, growth-driver genes, holdout distributions, per-cell fates, perturbation responses and gene-regulatory-network (GRN) structure. **b,** DynBench100 contains 100 scenarios across easy, medium, hard, and adversarial settings, with difficulty varied by gene number, fate number, confounders, noise level, and temporal holdout gap. **c,** CellCompass achieves the highest total scores among tested AI-agent configurations. **d,** Task-level scores across six dynamical objectives show broad CellCompass performance, with growth-driver identification remaining the most difficult objective across agents. **e**, CellCompass retains the highest total scores across increasing task difficulty. **f**, Repeated evaluation on 20 selected scenarios confirms the CellCompass advantage. Bars show the mean and s.d. across 60 runs from 20 scenarios with three runs each. Two-sided paired Wilcoxon signed-rank tests used the scenario means (n = 20), with Holm correction across five comparisons. ***adjusted P < 0.001. g, Total scores with and without the dynamical engine across all 40 medium-complexity scenarios. Each point represents one scenario, averaged over two runs per condition. Lines connect the same scenario, and black horizontal bars show the group means. Both conditions used GPT-5.5 with high reasoning and the same per-run budget. P is from a two-sided paired Wilcoxon test with Holm correction across the total score and six component scores.

DynBench100 comprises 100 scenarios spanning easy, medium, hard, and adversarial settings. Task difficulty was increased by raising the number of genes and terminal fates, and by adding noise, time-correlated confounders and larger temporal holdout gaps (Fig. 2b). Predictions were evaluated using six equally weighted dynamical objectives: velocity-field accuracy, growth-driver recovery, holdout distribution matching, per-cell fate prediction, perturbation-response recovery, and gene regulatory network (GRN) recovery. The six objectives are defined in the Methods and Supplementary Note S4.

Across DynBench100, CellCompass achieved the highest overall performance among the tested agent configurations: CellCompass with GPT-5.5 high reached a total score of 0.712, followed by CellCompass with MiMo-v2.5-pro at 0.667, exceeding both the general coding agents and the biomedical agent (Fig. 2c). The GPT-5.5 high configuration led across five of the six individual objectives (velocity, holdout distribution, fate, perturbation response and regulatory recovery; Fig. 2d), whereas growth-driver recovery remained the most challenging objective (mean agent scores 0.26–0.28) because effective growth in DynBench100 depends on a hidden subset of reporter genes while confounders correlate more strongly with time, causing simple correlation-based strategies to prioritize the wrong genes (Methods; Supplementary Note S4).

The top CellCompass configuration retained the highest total score at every difficulty level (Fig. 2e). To test run-to-run stability, we evaluated 20 selected scenarios three times each. CellCompass with GPT-5.5 high again achieved the highest mean total score (0.672) and remained significantly better than the other agents after paired two-sided Wilcoxon signed-rank testing with Holm correction (Fig. 2f; Methods).

To examine the contribution of the dynamical engine, we compared CellCompass with a version that retained scientific reasoning, review and general-purpose Python but lacked the dynamical engine and its training and simulation tools. Across all 40 medium-complexity scenarios, with two runs per condition, the mean six-task score was 0.736 with the engine and 0.672 without it. CellCompass scored higher in 35 of 40 scenarios, with the largest gains in fate and regulatory recovery (Fig. 2g and Supplementary Fig. 2).

### CellCompass reveals distinct EMT routes across cancer cell lines and stimuli

The same EMT-inducing stimulus can produce distinct cellular futures depending on cancer-cell context. We focused on the EMT time-course scRNA-seq dataset, which profiles EMT induction across A549 lung adenocarcinoma, DU145 prostate carcinoma, MCF7 breast carcinoma, and OVCA420 ovarian carcinoma cells treated with EGF, TGF-β1, or TNF over 0 d, 8 h, 1 d, 3 d, and 7 d (Fig. 3a; Methods)^75^. To capture both transcriptional progression and effective abundance changes across the 12 cell-line × stimulus time courses (34,994 cells, 11,058 genes), CellCompass selected the existing velocity–growth flow-matching (VGFM) formulation and fitted condition-specific models under a shared preprocessing and gene-program framework (Fig. 3b; Methods; Supplementary Note S5).

**Figure 3:**
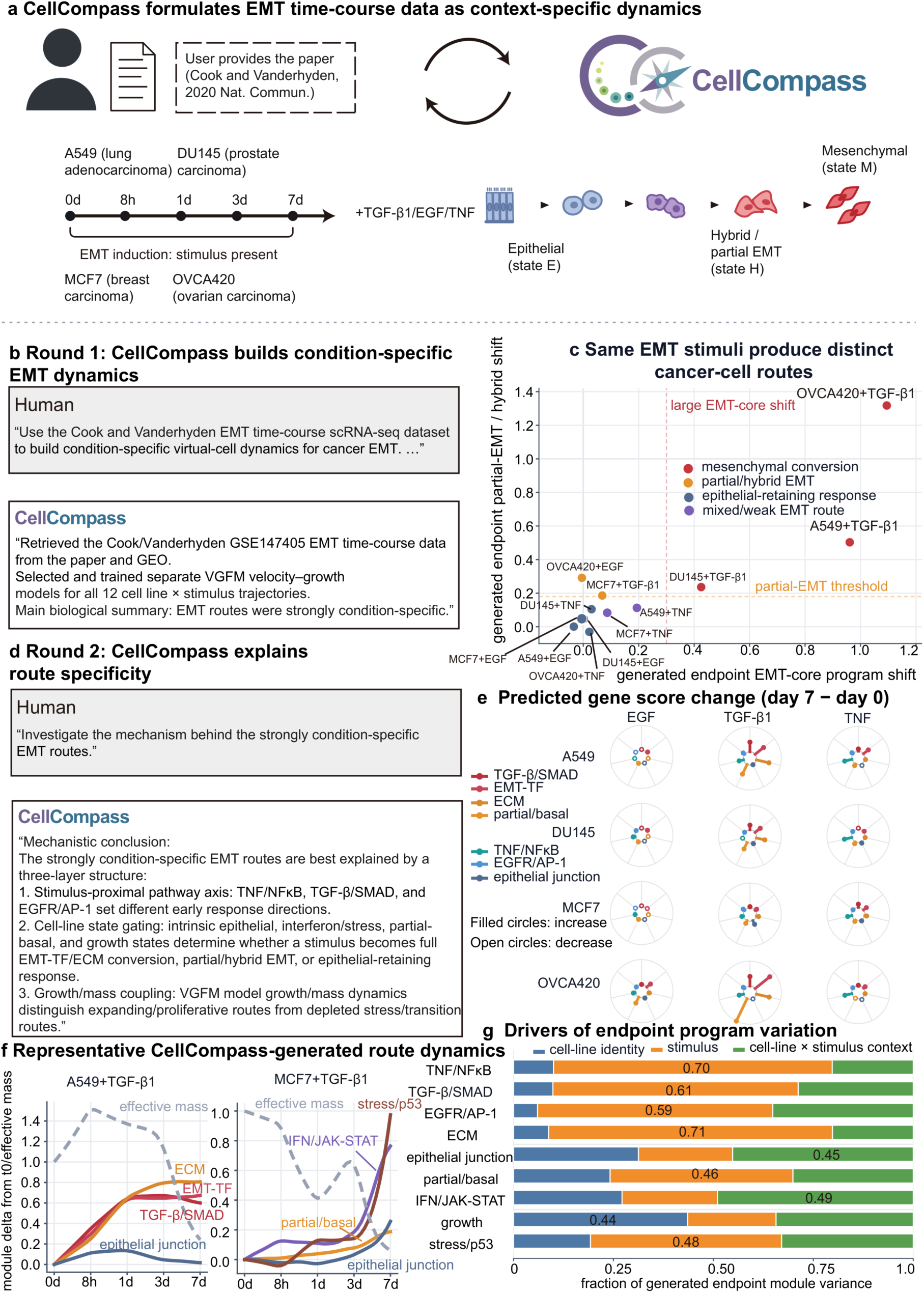
CellCompass reconstructs EMT dynamics that differ across cancer-cell states and stimuli. **a,** Overview of the EMT time-course scRNA-seq dataset and its analysis with CellCompass. A549 lung adenocarcinoma, DU145 prostate carcinoma, MCF7 breast carcinoma and OVCA420 ovarian carcinoma cells were treated with EGF, TGF-β1 or TNF and sampled at day 0, 8 h, day 1, day 3 and day 7. **b,** First interaction round. CellCompass obtained and prepared the dataset and fitted a separate velocity–growth flow-matching (VGFM) model to each of the 12 cell-line–stimulus combinations. **c,** Generated EMT routes. Each point represents one combination, positioned by the predicted day-0-to-day-7 changes in EMT-core score (x axis) and partial-EMT/hybrid score (y axis). Colors indicate route classes. Dashed lines mark the EMT-core (0.30) and partial-EMT/hybrid (0.18) thresholds. **d,** Second interaction round. CellCompass examined route specificity in terms of stimulus-responsive pathways, cancer-cell state and population growth. **e**, Predicted gene-program changes from day 0 to day 7. Rows indicate cell lines and columns indicate stimuli. Within each circular plot, colored lines represent the indicated gene programs, with the same color and direction used across plots. Line length represents the magnitude of the score change on a shared scale. Filled circles indicate increases and open circles indicate decreases. **f,** Generated trajectories for A549 and MCF7 cells treated with TGF-β1. Solid lines show program-score changes relative to day 0. Gray dashed lines show effective mass, the total simulated particle weight, initialized at 1. **g,** Fractions of endpoint program variation attributed to cell-line identity (blue), stimulus (orange) and the remaining cell-line–stimulus context (green). Numbers indicate the largest fraction in each bar.

The generated trajectories showed that EMT progression was strongly context-dependent (Supplementary Figs. 3–4). Each stimulus produced different generated routes across cancer-cell backgrounds, ranging from mesenchymal conversion through partial or hybrid EMT to epithelial-retaining and weakly responsive routes (Fig. 3c). TGF-β1 produced the strongest EMT-associated changes, with large EMT-core and partial/hybrid endpoint shifts in A549 and OVCA420 but a more moderate EMT-core shift in DU145. EGF and TNF responses were generally weaker or more context-dependent, with several conditions retaining epithelial character despite stimulus exposure. These context-dependent responses were consistent with those reported in the original study^75^. The fitted dynamics describe how each endpoint is approached and how effective population abundance changes along the route.

CellCompass then analyzed why the generated routes separated across conditions. Using the dynamics fitted separately for each cell-line–stimulus combination, it annotated trajectory changes with three kinds of gene-program module: stimulus-proximal signaling (TGF-β/SMAD, TNF/NFκB, EGFR/AP-1), cell-state identity (epithelial junction, partial or basal hybrid, interferon/JAK–STAT and stress) and downstream remodeling (EMT transcription factors, extracellular matrix, growth-associated programs) (Fig. 3d,e; Methods; Supplementary Note S5). This analysis suggested a three-layer organization of route specificity. First, stimulus-proximal pathways set early response directions, with TGF-β/SMAD, TNF/NFκB, and EGFR/AP-1 modules separating the major stimulus axes. Second, the initial cancer-cell state gated the route taken by each stimulus, allowing similar stimuli to yield mesenchymal conversion, partial/hybrid EMT, or epithelial-retaining responses in different cell lines. Third, the VGFM effective-growth component showed that several conditions with strong EMT-associated transcriptional shifts had declining effective abundance, whereas others had weaker shifts with stable or increasing effective abundance (Fig. 3d–g and Supplementary Fig. 5).

Representative trajectories illustrated these context-dependent programs. In A549 cells treated with TGF-β1, the fitted VGFM dynamics generated a route with coordinated TGF-β/SMAD, EMT transcription-factor, and extracellular-matrix activation, together with a decline in effective abundance. In MCF7 cells treated with TGF-β1, the route was more partial/basal and stress-associated, with stronger interferon/JAK– STAT and stress/p53 dynamics and weaker mesenchymal conversion. These examples show how the same stimulus can be filtered through different initial cancer-cell states to produce distinct EMT-associated futures (Fig. 3f).

To quantify the relative contributions of stimulus and cell-line context, CellCompass decomposed generated endpoint module variation across the twelve conditions. Stimulus identity explained a large fraction of the variation in canonical pathway and remodeling modules, including the TNF/NFκB, TGF-β/SMAD and EGFR/AP-1 signaling modules and extracellular-matrix remodeling. In contrast, epithelial junction, interferon/JAK–STAT, and growth-associated modules retained larger cell-line or cell-line × stimulus components (Fig. 3g; Methods). Together, these results show that EMT progression across conditions is jointly shaped by stimulus identity and cancer-cell context, with route-specific changes in both transcriptional state and effective abundance.

### CellCompass designs PulseBridge to incorporate metabolic RNA labeling into drug-response dynamics

Metabolic RNA labeling separates newly synthesized from pre-existing RNA and so constrains the direction of molecular change during a drug response. Starting from the Well-TEMP-seq HCT116 colorectal cancer dataset of cells treated with 5-aza-2′-deoxycytidine (5-AZA-CdR, decitabine), CellCompass organized the control and labeled day-0 baseline states together with treatment days 1, 2 and 3 and their total, newly synthesized and pre-existing RNA measurements (Fig. 4a; Methods)^26^. Because existing dynamics did not explicitly use this directional information, CellCompass designed PulseBridge, a flow-matching formulation in which metabolic RNA measurements constrain the local direction of the conditional path while accounting for baseline control-to-day-0 labeling variation (Fig. 4a; Methods; Supplementary Note S6).

**Figure 4:**
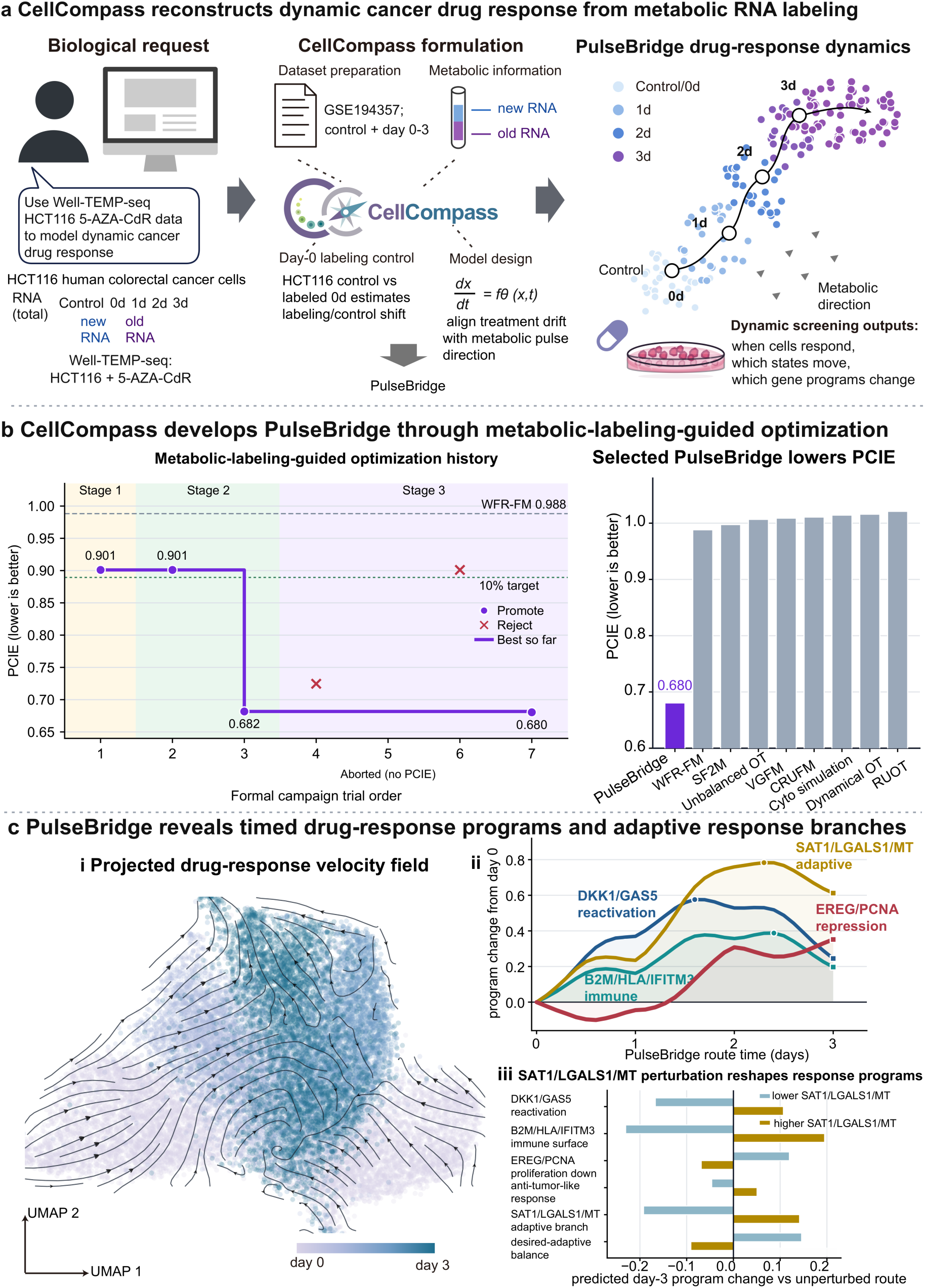
CellCompass designs PulseBridge to reconstruct cancer drug response dynamics from metabolic RNA labeling. **a,** CellCompass designed PulseBridge using Well-TEMP-seq data from HCT116 colorectal carcinoma cells treated with 5-aza-2′-deoxycytidine (5-AZA-CdR, decitabine). Measurements of total, newly synthesized and pre-existing RNA at days 0–3 constrain response dynamics, while the unlabeled and labeled day-0 cells identify labeling-associated variation. **b,** PulseBridge development and evaluation. Left, pulse-control identifiability error (PCIE) across seven trials. Circles indicate retained candidates and crosses rejected candidates. The aborted trial has no PCIE value and is labeled below the horizontal axis. The purple line tracks the lowest evaluated PCIE. Shading marks development stages, and dashed lines mark the Wasserstein–Fisher–Rao flow-matching (WFR-FM) reference and the 10% improvement target. Right, comparison with reference methods. PulseBridge achieved PCIE = 0.680, a 31.1% reduction relative to WFR-FM. **c,** PulseBridge drug-response dynamics and in silico perturbations. **i**, Observed cells colored by sampling day, with streamlines showing projected model velocities. **ii**, Program-score changes from day 0 along generated trajectories, showing reactivation (blue), immune-surface remodeling (teal), adaptation (gold) and EREG/PCNA repression (red). Positive red values indicate repression. Circles mark maxima and squares day-3 values. **iii**, Changes in predicted day-3 program scores after decreasing (blue) or increasing (gold) the SAT1/LGALS1/mitochondrial-gene program in initial cells, relative to unperturbed trajectories. The anti-tumor-like score combines DKK1/GAS5 reactivation, B2M/HLA/IFITM3 immune-surface remodeling and EREG/PCNA repression. The desired–adaptive balance is the difference between this combined score and the SAT1/LGALS1/mitochondrial-gene program score.

To evaluate directional fidelity, CellCompass defined pulse-control identifiability error (PCIE), which measures the discrepancy between simulated trajectory directions and held-out metabolic RNA-labeling directions after removing baseline labeling-associated motion (Methods). CellCompass then developed PulseBridge through staged optimization that progressively increased agreement between generated trajectories and metabolic RNA-labeling directions. Early candidate formulations improved PCIE relative to Wasserstein–Fisher–Rao flow matching (WFR-FM)^56^ but did not reach the 10% improvement target. Subsequent revisions strengthened the directional-disagreement penalty and the endpoint-preserving path adjustment, reducing PCIE from 0.901 to 0.682, and the final selected model reached PCIE = 0.680 (Fig. 4b; Methods; Supplementary Note S6). Compared with WFR-FM, the strongest baseline under this criterion, PulseBridge reduced PCIE by 31.1% and achieved a lower mean Wasserstein-1 distance (0.301 versus 0.308), indicating improved agreement with metabolic directions while preserving agreement with the observed treatment-state distributions (Fig. 4b; Supplementary Fig. 6; Methods).

PulseBridge ordered the HCT116 response as a continuous progression from day-0 states through intermediate day-1 and day-2 response states toward day-3 treated states, consistent with the temporal structure measured by Well-TEMP-seq (Fig. 4c,i). Program-level analysis along PulseBridge trajectories revealed a staged sequence of drug-response programs: DKK1/GAS5 reactivation emerged early, followed by B2M/HLA/IFITM3 immune-surface remodeling and SAT1/LGALS1/MT adaptive-metabolic activation, whereas EREG/PCNA repression became most prominent by day 3 (Fig. 4c,ii; Methods).

The trained PulseBridge dynamics also supported in silico perturbation of candidate late-response programs. Decreasing the SAT1/LGALS1/MT adaptive-metabolic program in initial cells strengthened predicted day-3 EREG/PCNA repression and improved the net desired-versus-adaptive balance while lowering DKK1/GAS5 reactivation and immune-surface remodeling, whereas increasing it shifted these programs in the opposite direction (Fig. 4c,iii; Supplementary Fig. 7). These results nominate the SAT1/LGALS1/MT program as a candidate adaptive component of the 5-AZA-CdR response whose suppression during treatment would test this predicted trade-off.

### CellCompass identifies clone-supported lineage priming among transcriptionally similar progenitors

Clonal lineage information constrains fate divergence that an RNA time course alone cannot resolve. In the LARRY mouse hematopoiesis time-course dataset, day-2 progenitors were profiled together with day-4 and day-6 descendants, sparse clone barcodes and fate annotations (Fig. 5a and Methods)^22^. Nearby day-2 progenitors can occupy similar transcriptional states yet give rise to different day-6 barcode descendants (Supplementary Fig. 8). CellCompass framed this as a clone-informed fate-dynamics problem, asking how lineage information should enter the dynamics so that future fate probabilities can be predicted from early single-cell states.

**Figure 5:**
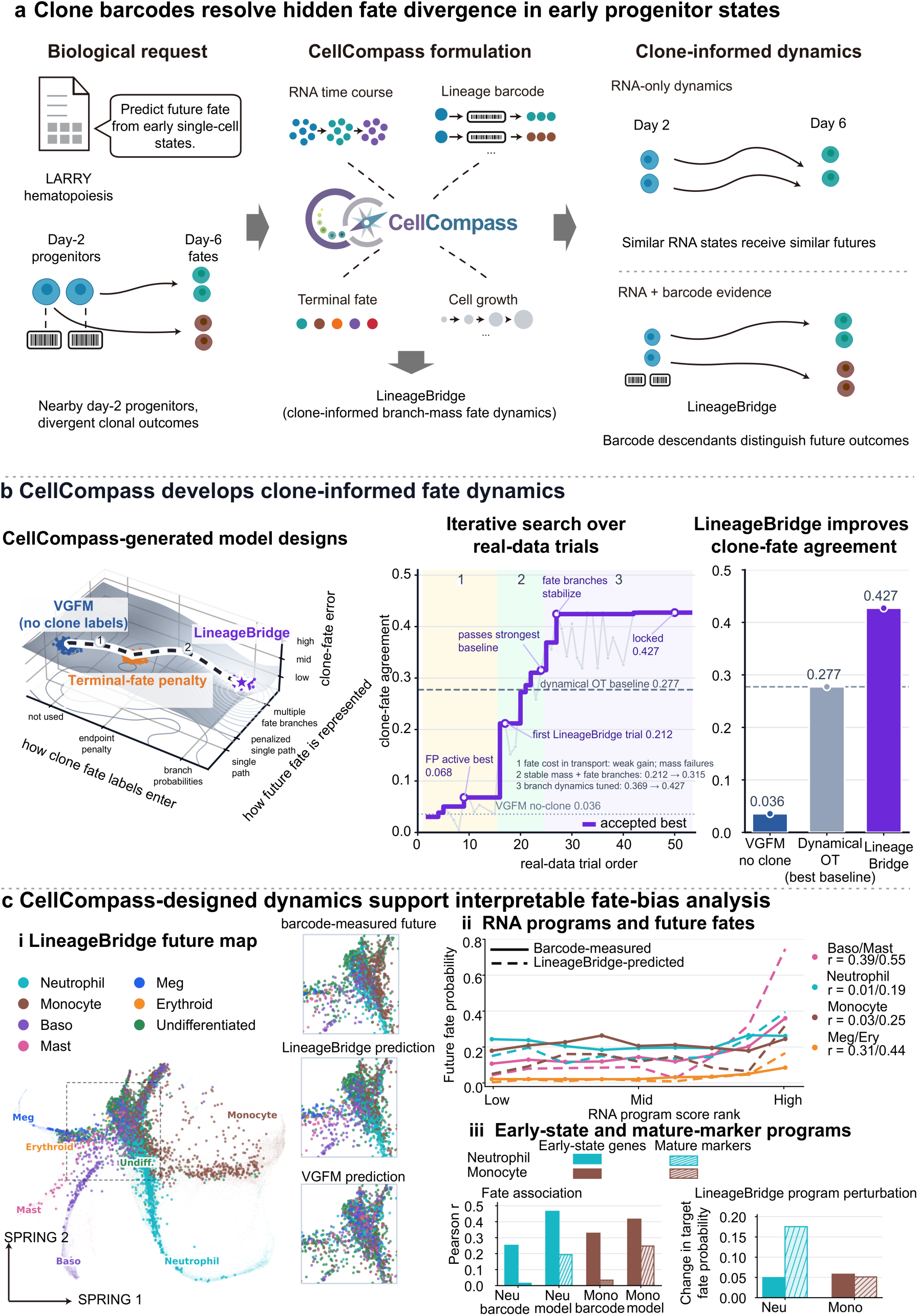
CellCompass designs LineageBridge for clone-informed fate dynamics in lineage-traced hematopoiesis. **a,** CellCompass used the LARRY hematopoiesis dataset to design LineageBridge, incorporating clone barcodes into velocity–growth dynamics to predict day-6 fates from day-2 RNA states. The schematic illustrates how clone information distinguishes future outcomes among transcriptionally similar progenitors. **b**, Model design and evaluation. Left, comparison of RNA-only velocity–growth flow matching (VGFM), a formulation with a terminal-fate penalty (FP) and LineageBridge with branch-mass fate dynamics. Middle, clone-fate agreement across candidate trials. The purple line tracks the best accepted model, background shading marks development stages and horizontal reference lines indicate baseline performance. Right, LineageBridge achieved clone-fate agreement of 0.427, compared with 0.277 for dynamical optimal transport and 0.036 for RNA-only VGFM. **c,** Fate prediction and downstream analysis of lineage-associated RNA programs. **i**, Day-2 cells colored by their most probable LineageBridge-predicted fate on the SPRING map. Insets compare barcode-measured, LineageBridge-predicted and VGFM-predicted fates within the outlined region. **ii**, Cells ranked by RNA-program score and divided into nine approximately equal-sized groups. Solid and dashed lines show mean barcode-measured and LineageBridge-predicted fate probabilities, respectively. Paired r values report Pearson correlations between program scores and barcode-measured/model-predicted probabilities. Basophil and mast fates are combined for this analysis, as are megakaryocyte and erythroid fates. **iii**, Correlations of early-state gene programs and mature-marker programs with neutrophil or monocyte fate probabilities. Lower right, changes in predicted fate probability after shifting initial RNA states by +5 times the mean principal-component loading vector of the indicated genes before simulation (Supplementary Note S7). Filled bars represent early-state gene programs and striped bars represent mature-marker programs.

CellCompass designed LineageBridge as an unbalanced velocity–growth flow-matching model in which clone barcodes restrict plausible transport between observed time points, terminal clone composition allocates transported mass among future fate branches and branch mass represents fate probabilities. The model therefore uses clone-derived future fate information to supervise hidden fate branches while preserving unbalanced population dynamics across day-2, day-4 and day-6 snapshots (Fig. 5a, Methods and Supplementary Note S7).

CellCompass next compared three ways of introducing clone information into the dynamical model: an RNA-only velocity–growth model without clone labels, a formulation that used clone information only through a terminal-fate mismatch penalty and a branch-mass formulation in which future fate is represented by multiple latent fate branches (Fig. 5b). Among the dynamical formulations tested, adding a terminal-fate penalty gave smaller gains than using clone information to allocate mass among branches. Introducing branch-mass fate dynamics produced a sharp increase in clone-fate agreement, and subsequent stabilization of branch probabilities and mass dynamics further improved the model from the first branch-mass version to the final selected model (Fig. 5b and Methods). The final LineageBridge model achieved clone-fate agreement of 0.427, compared with 0.036 for the no-clone VGFM model and 0.277 for the strongest dynamical optimal-transport baseline, corresponding to a 54% improvement (Fig. 5b; Supplementary Fig. 9). Brier error also improved, supporting better agreement between predicted fate probabilities and day-6 fate distributions estimated from clone barcodes (Supplementary Fig. 10).

The learned fate dynamics produced an interpretable future-fate map for day-2 progenitors, assigning early cells to regions biased toward neutrophil, monocyte, basophil, mast, megakaryocyte, erythroid and undifferentiated outcomes that were not recovered equally by RNA-only dynamics (Fig. 5c). In a transcriptionally similar early-state region, LineageBridge predictions better matched barcode-measured future fates than the VGFM prediction, indicating that clone-informed branch dynamics preserved divergent clonal outcomes that were weakly resolved by RNA state alone (Fig. 5c; Supplementary Figs. 8 and 11).

Focusing on the four major differentiated trajectories (neutrophil, monocyte, combined basophil/mast, and combined megakaryocyte/erythroid; Methods), LineageBridge further resolved early lineage-priming programs. Whereas basophil/mast and megakaryocyte/erythroid fates tracked continuous transcriptional gradients, neutrophil and monocyte fate biases at day 2 were associated far more strongly with early-state priming modules than with canonical mature markers (Fig. 5c). For neutrophil fate, an early-state program led by *Calr*, *Sell*, *Emb* and *Gsr* (eight genes in total, Supplementary Note S7) correlated more strongly with barcode-measured future bias than the mature neutrophil markers *Cebpe*, *S100a8*, *S100a9* and *Lyz2*, and LineageBridge predictions followed the same trend. The same held for monocyte fate, where an early-state program including Rbms1, Sirpa, *Casp6* and *Cd44* tracked future monocyte bias better than the mature markers *Irf8*, *Csf1r*, *Lyz2* and *Ly6c2*. In the fitted dynamics, shifting initial RNA states along either the early-state or the mature-marker direction increased the predicted probability of the corresponding fate (Fig. 5c and Supplementary Note S7; additional program perturbations in Supplementary Fig. 12). By representing clone composition through branch mass, LineageBridge resolved fate divergence among transcriptionally similar progenitors and linked it to early gene programs that precede mature lineage markers.

### CellCompass traces fate-specific RNA-to-surface-protein timing from initial RNA states

Surface-protein phenotypes need not change synchronously with the transcripts associated with them, so predicting future protein states from RNA requires a model that connects these molecular layers over time. CellCompass was applied to a CITE-seq hematopoietic time-course dataset of CD34+ hematopoietic progenitors measured across days 2, 3, 4, 7, and 10, with paired RNA profiles and 103 antibody-derived tag (ADT) surface-protein measurements (Fig. 6a; Methods)^76^. This task extends temporal RNA modeling from movement through RNA state space to generation of time-resolved surface-protein phenotypes from an initial RNA state. CellCompass formulated the problem as RNA-initialized protein dynamics and designed ProteinBridge, implemented as CITE paired-coordinate flow matching, to learn RNA trajectories together with a time-dependent surface-protein state (Fig. 6a; Methods; Supplementary Note S8).

**Figure 6:**
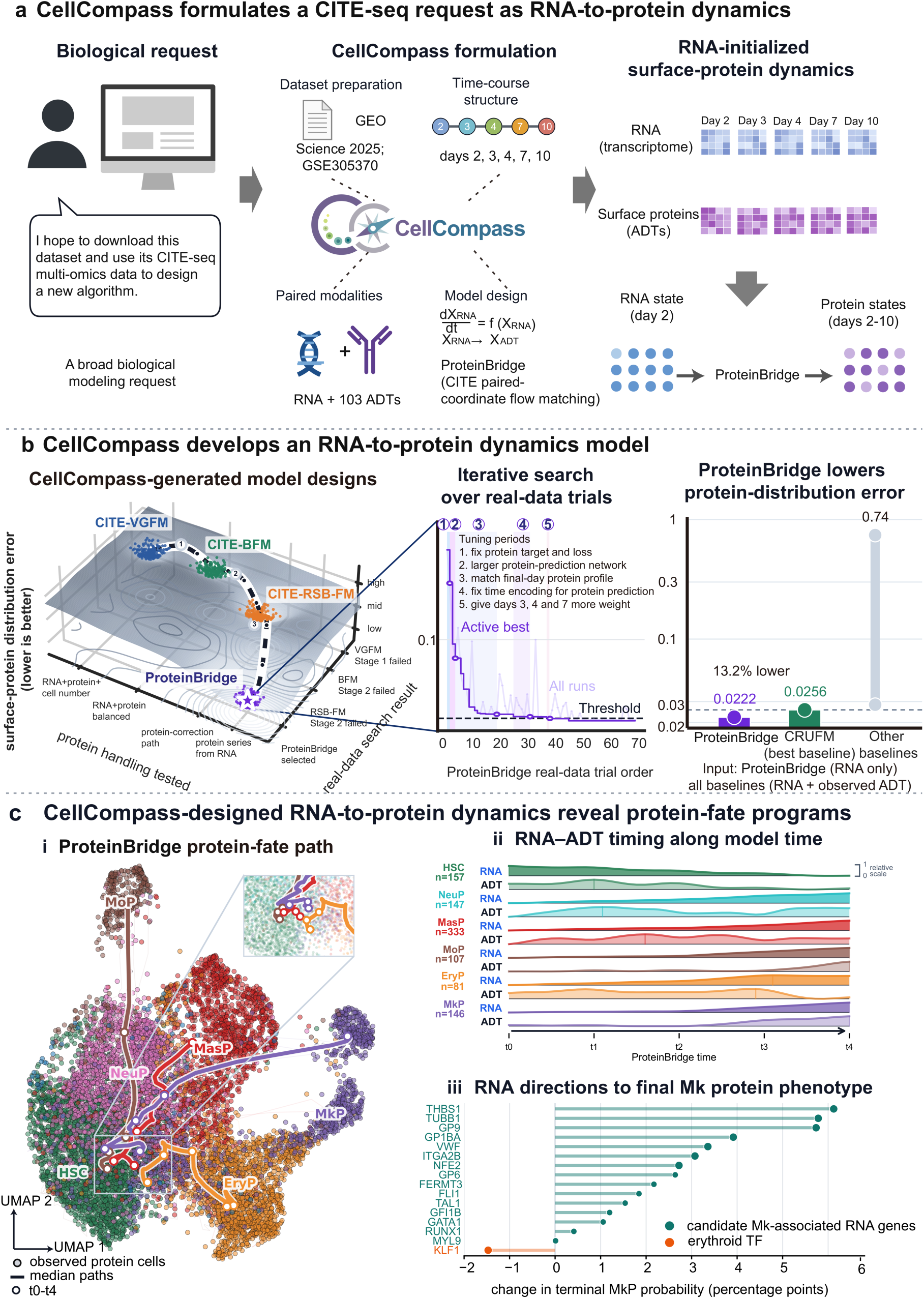
CellCompass designs ProteinBridge for RNA-initialized surface-protein dynamics from CITE-seq time-course measurements. **a,** CellCompass used paired RNA and 103 antibody-derived tag (ADT) measurements from a hematopoietic time course spanning days 2, 3, 4, 7 and 10 to design ProteinBridge. **b,** Model design and evaluation. Left, candidate formulations explored by CellCompass, including unbalanced velocity–growth (CITE-VGFM), balanced (CITE-BFM) and residual stochastic-bridge (CITE-RSB-FM) flow matching. Middle, protein-distribution error across successive trials. The light purple line shows individual trial results, and the dark purple line tracks the active best model. Right, comparison with reference methods on a logarithmic scale. ProteinBridge achieved an error of 0.0222, 13.2% lower than CRUFM. The gray line spans the minimum and maximum errors of the other baselines (0.028–0.74). ProteinBridge used RNA-only initial input, whereas the baselines used RNA and observed ADTs. **c,** Surface-protein trajectories and biological analysis. **i**, Observed cells colored by protein phenotype, with median generated paths initialized from day-2 hematopoietic stem cell (HSC) RNA states. Labels indicate neutrophil (NeuP), mast-cell (MasP), monocyte (MoP), erythroid (EryP) and megakaryocyte (MkP) progenitors. White circles mark model times along the paths. **ii**, RNA- and ADT-program profiles along generated trajectories, each scaled to 0–1 to compare their timing. Vertical marks indicate peaks, and n denotes the number of simulated HSC-derived trajectories assigned to each terminal group. **iii**, Changes in terminal megakaryocyte-progenitor (MkP) probability, in percentage points, after a +5-standard-deviation shift along each indicated gene’s RNA direction. Green denotes candidate megakaryocyte-associated genes and orange denotes the erythroid reference KLF1.

During model development, CellCompass compared several candidate formulations that differed in how RNA and protein information entered the dynamics (Fig. 6b). Early designs either treated RNA and protein as a generic joint state or failed to reproduce future surface-protein distributions. ProteinBridge instead separated RNA-state dynamics from protein prediction by propagating RNA from the initial state while using a learned RNA–time mapping to predict the corresponding surface-protein state. Revisions to protein targets, mapping capacity, time encoding and intermediate-time weighting progressively reduced surface-protein distribution error, leading CellCompass to select the paired-coordinate formulation as the final model (Fig. 6b; Methods; Supplementary Note S8).

In the evaluation, ProteinBridge achieved the lowest surface-protein distribution error among the tested CITE-seq dynamics models, with a protein energy distance of 0.0222 compared with 0.0256 for conditional regularized unbalanced flow matching (CRUFM), the strongest baseline under this metric, corresponding to a 13.2% reduction (Fig. 6b; Supplementary Figs. 13–14; Methods). ProteinBridge also reduced RNA– protein cross-covariance error relative to CRUFM (0.467 versus 0.525), indicating improved preservation of RNA–protein coordination. ProteinBridge was initialized from RNA alone at inference, whereas competing baselines used RNA together with observed ADT information as input in the benchmark setting.

On the observed CITE-seq protein landscape, trajectories initialized from hematopoietic stem cell (HSC)-like RNA states moved toward surface-protein phenotypes corresponding to neutrophil, mast-cell, monocyte, erythroid and megakaryocyte progenitors (Fig. 6c). RNA–ADT timing profiles further revealed fate-specific temporal offsets between transcriptional and surface-marker programs, which reached their maxima at distinct model times across lineages (Fig. 6c; Supplementary Note S8).

We next examined which changes in initial RNA state increased the predicted probability of a terminal megakaryocytic protein phenotype. From HSC-like initial states, increasing megakaryocyte-associated RNA directions raised terminal megakaryocytic protein-state probability, with the strongest positive effects for platelet and megakaryocyte genes including *THBS1*, *TUBB1*, *GP9*, *GP1BA*, *VWF* and *ITGA2B* (Fig. 6c). In contrast, the erythroid transcription factor KLF1 shifted predictions away from the megakaryocytic phenotype, consistent with known megakaryocyte–erythroid biology and with the observed RNA–surface-protein associations in the dataset (Supplementary Fig. 15).

## Discussion

For a virtual cell model to make useful predictions, its dynamics must reflect the biological information provided by the experiment, posing a scaling problem different from that of cellular representation learning: larger datasets and additional modalities can often be incorporated into a shared representation, whereas new measurements may uncover relations that no existing dynamical model can describe. Here we propose CellCompass, an agentic model-design framework that addresses this challenge by making dynamical formulation part of the analysis itself. The scientific agent identifies the relation exposed by the experiment, and the extensible dynamical engine provides the mathematical components needed to represent it.

By automating model design within a structured dynamical framework, CellCompass outperformed both general-purpose coding agents and a biomedical agent on DynBench100. Recent virtual-cell benchmarks have also begun to evaluate zero-shot generalization across cellular contexts^77,78^. Whereas those benchmarks focus on cross-context perturbation prediction, DynBench100 evaluates whether an agent can recover hidden dynamical structure across six dynamical tasks.

CellCompass also differs from general-purpose scientific agents, including emerging multi-agent frameworks for therapeutic discovery^66–74^, and from cellular foundation models^6,8–13^. General-purpose AI agents can inspect data, retrieve literature, and write code, but each task typically begins with a broad space of possible analyses and usually follows established pipelines. CellCompass instead operates within a structured dynamical model space where reusable mathematical modeling principles and components can be adapted to new experimental measurements to develop tailored algorithms. Meanwhile, cellular foundation models learn general representations of cell state, whereas CellCompass determines how those states evolve under the constraints supplied by an experiment. Learned representations can therefore define the state spaces on which CellCompass constructs dynamics, while dynamical models provide those representations with temporal, perturbational and fate-predictive structure.

The biological applications of CellCompass demonstrate why different experimental measurements require different dynamical structures. In the EMT study, the relevant dynamics were already captured by movement through transcriptional state space together with changes in effective abundance, and CellCompass retained the existing model, whereas the other three studies introduced information that required additional dynamical structure: metabolic labeling resolved the temporal ordering of the 5-AZA-CdR response and identified a candidate adaptive program; clonal barcodes revealed fate bias among transcriptionally similar progenitors before mature lineage markers became informative; and paired RNA– protein measurements showed that the offset between transcriptional and surface-protein peaks differs among lineages.

Several methodological limitations point to future directions. First, snapshot-derived effective growth rates capture net population change but cannot separate cell division, death and sampling efficiency without division-tracking assays or spike-in controls. Second, clone-informed branch allocation depends on experimental barcode recovery depth, and in silico perturbation predictions require prospective experimental validation. Third, while our current engine focuses on temporal trajectories across destructive snapshots, extending dynamical design to spatial and history-dependent processes will require adding spatial interaction kernels and non-Markovian state representations. Finally, although reviewer agents check the mathematical proposal, its implementation and the prediction inputs, they can still miss implicit modeling assumptions or subtle errors in the equations and code. Mathematical formulations and biological predictions therefore require expert inspection before experimental follow-up.

CellCompass is designed to expand as biological measurements evolve. In the future, new modalities can introduce new dynamical constraints, while validated formulations can be reused across related studies. Connecting model development to prospective experiments is a shared goal of recent virtual-cell frameworks, AI-enabled virtual laboratories and agentic experimental systems^79–83^. For CellCompass, an important next step is to use model-derived predictions to guide new measurements and then test whether those measurements support or require revision of the proposed dynamics. This framework creates a path toward virtual cells whose dynamical capabilities grow together with biological measurement.

## Methods

### CellCompass architecture for virtual-cell model design

CellCompass treats dynamical model design as a mapping from experimental measurements to model constraints. For each study, the system first defines the biological question and future-state prediction target, then identifies the relationship between present and future cells that is supported by the available measurements. It next determines whether this relationship can be represented by an existing formulation or requires modification of the dynamics. When a new formulation is required, CellCompass specifies which component of the model should change, how the additional measurement enters that component and how recovery of the intended relationship will be evaluated. The system combines this scientific-agent workflow with an extensible dynamical engine that provides reusable model families, fitting procedures, simulation routines and evaluation metrics (Fig. 1).

### Scientific agent and five-stage workflow

In the five-stage workflow (Fig. 1c), the scientific agent first prepares the data and separates measurements used for model fitting from inputs available at trajectory generation and observations reserved for evaluation. After mapping the experimental measurements onto dynamical constraints, the agent either applies an existing formulation without altering its mathematical structure or develops a new formulation by modifying existing components, combining components across model families or introducing new governing equations. Each new proposal is grounded in theory and prior work and reviewed before implementation, after which the resulting trajectories are benchmarked against a fixed set of reference methods using the prespecified evaluation criterion and used for biological analysis.

In CellCompass, candidate formulations can return to earlier stages during development: failure to recover the prespecified relationship returns the study to dynamics design to revise the relevant model component, whereas evidence that the assumed measurement–dynamics relationship is unsupported or insufficiently identifiable returns the study to constraint mapping.

The agent workflow separates model proposal from model approval. Within the scientific agent, a design agent specifies the biological constraint, mathematical formulation, prediction inputs and evaluation plan, while context-isolated reviewer agents check that the proposed relationship is supported by the data, that the code implements the approved formulation, that the planned comparison tests that relationship and that forward prediction uses only initial cell states and fitted parameters before the candidate is benchmarked. Throughout this workflow, the agents invoke executable tools for data preprocessing, model fitting, simulation and evaluation, and all model specifications, reviewer decisions, candidate implementations and evaluation results are recorded (Supplementary Note S1). In this study, CellCompass used GPT-5.5 with the high reasoning setting to design PulseBridge, LineageBridge and ProteinBridge.

### Extensible dynamical engine and its model family

All models in the dynamical engine share a common population-dynamics framework and allow CellCompass to modify three components: the coupling between cells at adjacent observation times, the conditional path connecting coupled cells and the quantities propagated or read out along that path (Fig. 1d). The latter can include effective mass, fate probabilities, score-based diffusion terms or molecular readouts, allowing CellCompass to incorporate additional experimental measurements by modifying the relevant components.

Because time-resolved single-cell assays capture cross-sectional snapshots across destructive sampling times, we represent each snapshot as a sample from an evolving cell-state distribution and infer the dynamics connecting these distributions. At observation time *T_k_*, the *n_k_* sampled cells define the empirical measure

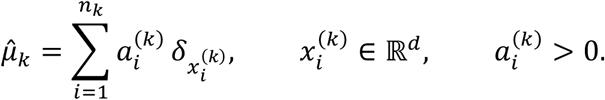

Here, 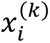 denotes the state of cell *i*, represented by its gene-expression profile or a low-dimensional embedding, and 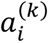 denotes its weight. Balanced formulations conserve total mass and represent each snapshot as a probability measure, whereas unbalanced formulations allow mass to vary between time points, accommodating changes in relative cell abundance arising from proliferation, death or sampling. Future populations are generated by simulating weighted particles whose states and weights evolve according to

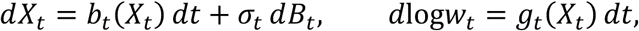

where *B_t_* is standard Brownian motion. The simulated population at time *t* is represented by 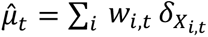. At the population level, the corresponding density *ρ_t_* satisfies a Fokker–Planck equation with a source term,

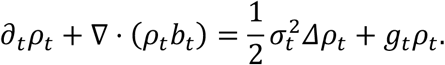

The drift *b_t_* describes cell-state transitions, *σ_t_* controls stochastic dispersion, and the source field *g_t_* determines state-dependent changes in particle weight; in stochastic formulations, the drift decomposes into the current velocity and the score-based diffusion term (Supplementary Note S2).

We use regularized unbalanced optimal transport (RUOT) to describe the relationship between the transport and bridge formulations implemented in the engine^49,50^. For each pair of successive observed populations, RUOT jointly determines the density path, drift and source field by minimizing

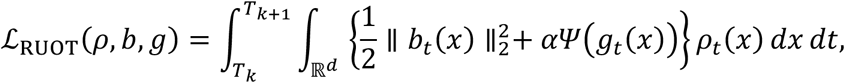

subject to the Fokker–Planck equation and endpoint constraints 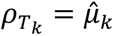 and 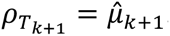. The convex function *Ψ* penalizes local mass change, and *α* sets its cost relative to transport through state space. Restricting *Ψ*(*g*) to be finite only at *g* = 0 enforces mass conservation and, with non-zero diffusion, yields a balanced Schrödinger bridge. In the zero-diffusion limit, a quadratic penalty *Ψ*(*g*) = *g*^2^, up to rescaling, recovers dynamic unbalanced optimal transport with a Wasserstein–Fisher–Rao transport–growth action^51^. Further imposing *g* = 0 and equal endpoint mass recovers Benamou–Brenier dynamic optimal transport^84^. These formulations thus differ in their treatment of cell-state transport, stochastic dispersion and effective growth within the same population equation, with the associated variational problems and limiting conditions given in Supplementary Note S2.

The engine estimates these dynamics from finite samples using conditional flow matching^53^. For each interval *T_k_* < *T_k_*_+1_, a coupling *Γ_k_* assigns mass to source–target cell pairs (*x*, *y*). Conditional on a pair, the path construction samples a normalized time *τ* ∈ [0,1], an intermediate state *Z_τ_* and the corresponding field targets. The default deterministic interpolation is

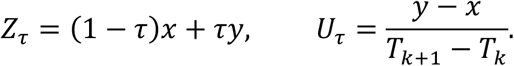

The velocity field is trained against *U_τ_*; for unbalanced formulations, the coupling also specifies the terminal mass assigned to each source particle and a conditional mass path provides the growth target *G_τ_*, whereas stochastic formulations additionally provide a denoising score target *S_τ_*. The relevant fields are fitted jointly using

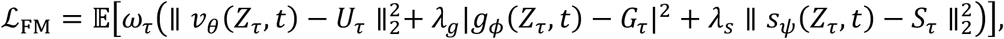

with terms omitted when the corresponding field is absent and, in stochastic formulations, the score error additionally weighted by the squared denoising scale (Supplementary Note S3). Among the balanced formulations, optimal-transport conditional flow matching combines a balanced coupling with a deterministic path and learns only the velocity field, whereas simulation-free Schrödinger bridges via score and flow matching (SF2M) learns current-velocity and score-based diffusion fields to model stochastic dynamics while conserving total population mass^54^. Among the unbalanced formulations, velocity–growth flow matching (VGFM) uses a source-relaxed unbalanced coupling to learn deterministic velocity and growth fields^55^, Wasserstein–Fisher–Rao flow matching (WFR-FM) uses Wasserstein–Fisher–Rao semi-couplings and their associated transport–growth paths^56^, and conditional regularized unbalanced flow matching (CRUFM) combines an unbalanced coupling with a stochastic conditional bridge to learn current-velocity, growth and score-based diffusion fields. Each built-in model is therefore specified by its coupling, conditional path and transported quantities.

These components are implemented as separate modules (Supplementary Notes S1 and S3), so extending the engine requires replacing only those affected by new experimental constraints while reusing the remaining training and simulation procedures. For example, PulseBridge modifies the coupling, conditional path and training loss; LineageBridge modifies the coupling and branch-mass allocation; and ProteinBridge adds paired-modality coupling and a protein readout network.

### Staged validation and model acceptance

To test recovery of inter-timepoint dynamics beyond endpoint fitting, CellCompass separates formulation design from model acceptance and evaluates each new formulation against a biological criterion specified from the measurement that motivated the model. Review of the mathematical proposal and code implementation, comparison with fixed reference methods, and a final confirmation step are applied before a newly designed formulation is accepted (Supplementary Fig. 1).

All candidates use the same processed data, initial cells, reference methods and evaluation code across three validation stages. The first stage checks numerical stability and trajectory validity, and the second tests the relationship that motivated the formulation, where acceptance requires at least a 10% improvement in the prespecified biological measure over the strongest comparable reference method without compromising agreement with observed populations. In the third stage, CellCompass further tunes the algorithm to improve agreement between predicted and observed cell populations while preserving the biological improvement established in the second stage.

After these stages, the formulation is fixed, including its equations, prediction inputs and numerical settings, and the full evaluation is repeated. Confirmed formulations are used for downstream analyses and retained in the engine for subsequent studies, whereas a failed confirmation returns the study to proposal development, implementation or evaluation according to the source of the discrepancy.

### Construction of the DynBench100 benchmark

DynBench100 evaluates whether a model-design system can recover multiple properties of a cellular dynamical process from snapshots. Each scenario was simulated from a stochastic gene-regulatory network in which self-activating, mutually inhibitory gene pairs generated alternative terminal states, while the expression of a specified subset of downstream genes determined cell birth and death. The simulator equations, gene roles and scenario settings are described in Supplementary Note S4.

DynBench100 comprises 100 scenarios (20 easy, 40 medium, 25 hard and 15 adversarial), with increasing difficulty introducing more fate modules, slow confounders and noise genes alongside longer trajectories and greater stochastic variation. Each scenario was sampled at 6–12 time points, where two or three intermediate time points were withheld from training to evaluate recovery of unobserved cell-state distributions. Detailed settings for each difficulty group are summarized in Supplementary Note S4.

For each scenario, CellCompass received the training snapshots, task description and a common terminal-state classifier. The simulator equations were withheld, together with the six ground-truth quantities that defined the benchmark evaluations: cell velocities, growth-driver genes, populations at withheld time points, root-cell fates, perturbation responses and the signed regulatory network (Supplementary Note S4).

### Comparison with general-purpose agents on DynBench100

CellCompass, OpenAI Codex, Claude Code with MiMo or DeepSeek, and Biomni^67^ received identical task descriptions, training snapshots, terminal-state classifiers and output specifications for each scenario. All submissions were assessed using the same evaluation code, with missing or invalid outputs receiving a score of zero for the affected component.

The primary CellCompass configurations used GPT-5.5 with the high reasoning setting and MiMo-v2.5-pro. Each configuration was run once across all 100 scenarios over the complete workflow from model selection or construction to generation of the six required outputs, and run-to-run stability was assessed on 20 selected scenarios using three runs per scenario with different random seeds.

The six component scores were reported separately and combined as an unweighted mean. Velocity recovery was measured by cosine agreement between predicted and simulated group-mean directions, and growth-driver and regulatory-edge recovery by area under the precision–recall curve (with regulatory direction scored separately); population prediction was evaluated at each withheld time using Wasserstein-1 distance, fate prediction by root-cell top-1 accuracy, and perturbation responses by active-response recovery and sign accuracy.

We compared predicted velocities with ground truth in the simulator’s gene-expression coordinates and time units by inverting each method’s coordinate and time transformations, including the diffusion term for continuous stochastic models. To evaluate perturbation responses, we set each selected gene in the initial cells to the simulator’s control or perturbed expression level and simulated the response with the fitted model, evaluating all six predictions from the same run (coordinate conversions, initial perturbations and score definitions in Supplementary Note S4).

We tested the contribution of the dynamical engine across all 40 medium-complexity scenarios using GPT-5.5 with high reasoning, running both versions twice per scenario with paired numerical seeds. We averaged the two runs within each scenario and compared the resulting 40 pairs using two-sided Wilcoxon signed-rank tests, with Holm correction across the total score and six component scores (Supplementary Note S4).

### EMT dynamics across cancer cell lines and stimuli

The epithelial–mesenchymal transition (EMT) analysis used GSE147405^75^, comprising A549, DU145, MCF7 and OVCA420 cells treated with EGF, TGF-β1 or TNF and sampled at 0 d, 8 h, 1 d, 3 d and 7 d. After excluding withdrawal samples, which represented a distinct experimental regime, the dataset contained 34,994 cells and 11,058 genes; counts were normalized to 10,000 per cell, log-transformed, restricted to 2,000 highly variable genes and projected onto 50 principal components.

In the first interaction round, CellCompass fitted a separate velocity–growth flow-matching (VGFM) model across the five time points for each of the 12 cell-line–stimulus combinations, preventing baseline differences between cell lines from being interpreted as within-condition transitions, and simulated trajectories forward from all earliest-time cells in each condition. In the second round, it compared the inferred EMT routes and regulatory programs across conditions.

Population distributions were compared using Wasserstein-1 distance after normalizing predicted and observed mass, whereas total simulated particle mass was retained as a measure of effective abundance. To follow pathway activity along the simulated trajectories, each gene-program score was calculated as the mean standardized expression of its constituent genes, and ridge regression was fitted on the observed cells to predict these scores from the 50-dimensional state representation. Temporal agreement was assessed using Pearson and Spearman correlations, and an additive analysis separated cell-line and stimulus effects (Supplementary Note S5).

### PulseBridge for metabolic-RNA-guided drug-response dynamics

PulseBridge was developed using Well-TEMP-seq measurements of HCT116 cells exposed to 5-aza-2′-deoxycytidine (5-AZA-CdR; decitabine), available under GSE194357^26^. The dataset contained 20,000 cells, with 4,000 cells from each of five samples: control, labeled day 0 and treatment days 1, 2 and 3. Total, newly synthesized and pre-existing RNA were represented in a shared 30-dimensional state space, where metabolic-velocity annotations provided local estimates of the direction of RNA production and turnover for the 16,000 labeled cells (12,816 used for training and 3,184 held out), while the 4,000 unlabeled control cells defined the unperturbed day-0 baseline. To evaluate directional recovery, pulse-control identifiability error (PCIE) measures the discrepancy between experimentally inferred metabolic directions and the direction of change between consecutive simulated cell states, evaluated on the 2,400 held-out directions from days 0, 1 and 2 because the simulated trajectories ended at day 3.

Metabolic RNA labeling provides information about the direction of molecular change within a sampling interval, whereas existing formulations used only the endpoint populations. PulseBridge therefore used the metabolic directions to guide both cell coupling and the intermediate paths between coupled cells while retaining balanced transport and deterministic simulation, with additional training penalties for directional disagreement and labeling-associated motion. Because labeling altered the day-0 cell-state distribution, CellCompass designed PulseBridge to first remove components of the measured directions aligned with three labeling-associated modes of variation estimated from the training cells. PulseBridge then adjusted the intermediate portion of each modeled path toward the remaining measured direction according to its confidence, with the adjustment vanishing at both endpoints so that the coupled initial and terminal cell states remained unchanged.

Model development completed in 5.3 h across seven candidate trials; the six that ran to completion were evaluated by held-out PCIE and population Wasserstein-1 distance, with forward trajectories integrated from the combined control and labeled day-0 RNA states.

The confirmed model was used to order transcriptional response programs along simulated trajectories and to evaluate program-level perturbations against random shifts of the same norm propagated through the same fitted field (Supplementary Note S6).

### LineageBridge for clone-informed fate dynamics

LineageBridge was developed using the LARRY mouse hematopoiesis dataset^22^, comprising 4,638 cells at day 2, 14,985 at day 4 and 29,679 at day 6, represented by 2,447 genes. Clone barcodes linked cells sharing clonal ancestry across these three time points, and day-6 annotations defined each clone’s terminal fate composition.

To model fate divergence among transcriptionally similar progenitors, LineageBridge used clone barcodes to constrain transport between adjacent time points and used terminal clone composition to allocate each source cell’s transported mass among fate branches. From these branch-resolved couplings, the model learned branch-specific velocity and growth fields together with fate probabilities predicted from RNA state and time, allowing total cell abundance to vary over time while conserving source-cell mass across fate branches.

Development of LineageBridge took 21.5 h and comprised 54 candidate trials spanning three formulations (RNA-only VGFM, terminal-fate penalty and branch-mass allocation), evaluated on 1,728 eligible day-2 cells from 1,243 clones with forward trajectories initialized from day-2 RNA states. Clone-fate correlation (CFCorr) and reliability-weighted multiclass Brier error measured how closely the predicted fate probabilities of each eligible day-2 cell matched the observed terminal-fate proportions within its clone, while Wasserstein-1 distance and total-mass variation assessed agreement between simulated and observed cell-state populations.

Model fitting and CFCorr evaluation retained all 11 annotated day-6 cell classes, whereas the fate map displayed the seven primary outcomes (excluding the smaller eosinophil, lymphoid, Ccr7-positive dendritic and plasmacytoid dendritic populations) and trajectory-level gene-program summaries grouped basophils with mast cells and megakaryocytes with erythroid cells. To examine early fate priming, we ranked non-marker genes by their residual correlation with barcode-derived fate probabilities after controlling for mature lineage markers in eligible day-2 cells, and assessed the effects of perturbing each module in the fitted model (Supplementary Note S7).

### ProteinBridge for RNA-initialized protein dynamics

ProteinBridge was developed using a CITE-seq^27^ study (GSE305370) of human CD34-positive hematopoietic stem and progenitor cells from four healthy donors, sampled at days 2, 3, 4, 7 and 10^76^. Across the 10,000 analyzed cells (103 antibody-derived tags, ADTs), RNA profiles were reduced to 50 components using truncated singular value decomposition and surface-protein profiles to 30 principal components using PCA, with each component standardized to zero mean and unit variance.

ProteinBridge used paired RNA–protein measurements during fitting in two ways: first, protein distance contributed to the endpoint coupling cost to reduce matches between transcriptionally similar cells with distinct protein profiles; second, the resulting paired RNA–protein paths and endpoints were used to train a time-dependent RNA-to-protein mapping alongside the RNA velocity field, enabling protein levels to be predicted along simulated RNA trajectories initialized from day-2 RNA.

Model development spanned 28.2 h, during which CellCompass compared several model families and developed the selected paired-coordinate formulation over 70 trials, each fitted across all five observed time points and evaluated by comparing predictions initialized from 2,000 day-2 RNA profiles against the four later RNA–protein populations. Model acceptance was based primarily on the energy distance between predicted and observed surface-protein distributions, alongside RNA Wasserstein-1 distance for transcriptional population agreement and relative cross-covariance error for RNA–protein coordination.

Using trajectories simulated from the 972 day-2 hematopoietic stem cells, we compared the timing of RNA and surface-protein differentiation programs across lineages and examined which initial RNA-state shifts increased terminal megakaryocyte-progenitor commitment (Supplementary Note S8).

### Numerical implementation and statistical analysis

Across all studies, distributional shape agreement was evaluated via Wasserstein-1 distance on weight-normalized empirical measures, while relative population change in unbalanced models was quantified by total-mass variation (Supplementary Note S9). Repeated DynBench100 evaluations (20 scenarios × 3 runs) were compared at the scenario-mean level using two-sided paired Wilcoxon signed-rank tests with Holm step-down correction. Language-model token consumption and API-equivalent cost over the full development of LineageBridge are detailed in Supplementary Note S1.

## Acknowledgments

This work was supported by the National Natural Science Foundation of China (NSFC Nos. 12288101, 8206100646 and T2321001 to P.Z.) and the Fundamental and Interdisciplinary Disciplines Breakthrough Plan of the Ministry of Education of China (grant no. JYB2025XDXM502). We acknowledge the support from the High-performance Computing Platform of Peking University for computation.

## Author contributions

P.Z., Q.N. and Z.Z. conceived this research. Z.Z., Z.W. and P.Z. designed CellCompass. Z.W. and Z.Z. implemented the core components of CellCompass. B.Z. and T.Z. contributed to parts of CellCompass. Z.Z. and Z.W. designed and implemented DynBench100. Z.Z., Z.W., Q.T., B.Z. and T.Z. contributed to the evaluation of DynBench100. Z.Z. and Z.W. analyzed the data. All authors interpreted the results. Z.Z. and Q.T. prepared the visualizations. Z.Z., Z.W. and Q.T. wrote the original draft. P.Z. and Q.N. reviewed and edited the manuscript. P.Z. and Q.N. supervised the work. All authors read and approved the manuscript.

## Data availability

All datasets analyzed in this study are publicly available. The epithelial-mesenchymal transition time-course single-cell RNA-seq dataset is available from the NCBI Gene Expression Omnibus (GEO) under accession GSE147405 (ref. ^75^). The Well-TEMP-seq dataset is available from GEO under accession GSE194357 (ref. ^26^). The LARRY mouse hematopoiesis lineage-tracing dataset is available from GEO under accession GSE140802 (ref. ^22^). The longitudinal CITE-seq hematopoietic dataset is available from GEO under accession GSE305370 (ref. ^76^).

