## Supplementary Information for "Building virtual cell dynamics that evolve with biological measurements"

### S1 CellCompass architecture and model development

#### S1.1 CellCompass system architecture

CellCompass combines a scientific agent with an extensible dynamical engine for virtual-cell model design. Given a biological question and time-resolved single-cell measurements, the agent identifies the relationship between present and future cells that needs to be represented, then selects an existing formulation or develops a new one, evaluates the resulting predictions and uses the generated trajectories for biological analysis. This work proceeds through five stages: question and data setup, constraint mapping, dynamics design, model evaluation, and outputs and interpretation. The dynamical engine provides reusable model families together with executable tools for preprocessing, model fitting, trajectory simulation and metric calculation; these models describe how cellular states change over time and, where required, how cell abundance and fate probabilities change along the trajectories.

When an existing formulation is sufficient, CellCompass fits it without changing its mathematical structure. Otherwise, it specifies which model component should change, how the additional measurements enter that component and how the intended improvement will be evaluated. Proposals and implementations then undergo context-isolated review, in which reviewer agents work in separate conversations without the main agent's history, and new candidate formulations pass through feasibility testing, biological evaluation and further tuning, described below. CellCompass used GPT-5.5 with the high reasoning setting to design PulseBridge, LineageBridge and ProteinBridge.

#### S1.2 Agent procedure and execution

Algorithm 1 summarizes the study procedure, from model selection through context-isolated review to iterative refinement, and Algorithm 2 gives the corresponding flow-matching training and forward-integration routines. The main agent receives the scientific instructions, the current study information and descriptions of the available models and tools, and loads stage-specific instructions as the corresponding tasks are performed. Each reasoning task runs in a bounded tool-use loop that dispatches the requested tool calls in turn, appends their outputs or error tracebacks and enforces configurable limits on tokens, retries and completion checks.

**Algorithm 1:** CellCompass study procedure

```
Input: Biological question, single-cell data, available dynamical models, development
       settings
Output: Fitted model and evaluated predictions, or an unfinished study
Prepare the data and define the biological comparison
Distinguish fitting measurements, prediction inputs and evaluation observations
Assess whether an existing formulation represents the required biological relation
if an existing formulation is sufficient then
    Fit that model, generate trajectories and evaluate the predictions
    return the fitted model and its results
end
while development remains within the configured limits do
    Propose the model components to change and the comparison to test them
    Obtain a context-isolated proposal review
    if the proposal requires revision then
        Revise it using the review and repeat proposal review
    end
    if the proposal is approved then
        Implement its equations, training objective and prediction procedure
        Review the implementation and prediction inputs
        if implementation review requires revision then
            Revise the code and repeat implementation review
        end
        if implementation review is approved then
            Fit and simulate candidates using the same data and reference methods
            if fitting or simulation fails the feasibility criteria then
                Revise the implementation or formulation
            end
            else if the proposed biological improvement is not established then
                Revise the model and repeat the biological comparison
            end
            else
                Tune population agreement while retaining the biological improvement
                Fix the selected formulation and repeat its complete evaluation
                if final confirmation succeeds then
                    Retain the model in the dynamical engine and analyze its trajectories
                    return the fitted model and its results
                end
                Return to the step responsible for the failed confirmation
            end
        end
    end
end
return the study and the development step not yet completed
```

Algorithm 2 summarizes how the dynamical engine trains a proposed model and generates predictions: endpoint couplings supply paired cells and mass targets, conditional paths provide intermediate states and rate targets for neural-field regression (Supplementary Note S3), and the fitted fields are then integrated forward from the initial cells.

---

**Algorithm 2:** Training and prediction with the dynamical engine

---

**Input:** Reviewed formulation, fitting measurements, model times $T_0 < \cdots < T_K$, designated prediction inputs and evaluation observations

**Output:** Fitted model, simulated trajectories and population and biological comparisons

Construct the cell-state representation and group the fitting cells by model time<br>
Build the model's neural fields and any additional fate or molecular networks<br>
**foreach** *adjacent pair of times* $T_k, T_{k+1}$ **do**<br>
    Set $\Delta_k = T_{k+1} - T_k$<br>
    Compute the cell-pair cost from the designated fitting measurements<br>
    Estimate the balanced or unbalanced coupling specified by the formulation<br>
    Compute any terminal-mass targets and branch allocations<br>
**end**<br>
**while** *the prescribed fitting schedule is not complete* **do**<br>
    Sample cell pairs and, where required, branches according to their coupling weights<br>
    Draw a local path position $\tau \in [0, 1]$ and any path noise $\epsilon$<br>
    Set model time $t = T_k + \tau\Delta_k$<br>
    Construct the intermediate state $Z_\tau$ using the specified conditional path<br>
    Compute the local velocity and log-mass-rate targets<br>
    Divide these rate targets by $\Delta_k$ to express them per unit model time<br>
    Compute the path regression weight $\omega_{ij}(\tau)$<br>
    **if** *the model learns a score-based diffusion term* **then**<br>
        Compute its denoising scale using the stated noise and time convention<br>
    **end**<br>
    Evaluate the neural fields at $(Z_\tau, t)$ and any additional conditioning variables<br>
    Calculate the active velocity, growth and score losses<br>
    Add or replace losses specified by the new formulation<br>
    Update the model parameters by the prescribed optimizer<br>
**end**<br>
Construct prediction inputs from the initial cells and fitted model<br>
Initialize particle weights and any branch probabilities<br>
Integrate the learned fields over the requested model times<br>
Calculate fate probabilities or molecular profiles using the fitted prediction networks<br>
Compare these simulated states and weights with the evaluation observations<br>
Return the fitted model, trajectories and prespecified comparisons

---

#### S1.3 Implementing new dynamical models

When existing dynamics cannot represent the additional biological information, CellCompass modifies only the affected modules of the dynamical engine while inheriting the remaining components from an existing formulation (Table 1). For example, PulseBridge retains balanced transport but changes its cost, paths and loss; LineageBridge adds clone-informed coupling, branch allocation and branch-specific velocity and growth; and ProteinBridge adds protein information to the coupling and trains an RNA-to-protein network alongside the RNA velocity field. A custom training loop can also replace the standard trainer if the proposed equations require a different optimization procedure.

Table 1: Components used to implement a new CellCompass formulation.

| Component | Inputs | Outputs and role |
|---|---|---|
| Training data | Preprocessed profiles, time labels, auxiliary modalities | Time-grouped state tensors and aligned cell indices. |
| Coupling and paths | Adjacent populations, auxiliary modalities | Weighted cell pairs, intermediate states $Z_\tau$, rate targets $(U_\tau, G_\tau)$, loss weights $\omega_\tau$ and noise samples. |
| Neural fields | State dimension and architecture settings | Velocity, growth or score networks, plus any branch or molecular-readout networks. |
| Training objective | Sampled intermediate states, targets and network predictions | Regression and auxiliary losses parameterizing the velocity, growth or score fields. |
| Prediction | Initial cells $X_0$, model times and fitted parameters | Forward-integrated state trajectories, particle weights and branch or molecular readouts. |
| Evaluation | Simulated trajectories and observed populations | Population distances ($W_1$, total-mass variation) and the prespecified biological metric. |

Across these modules, auxiliary modalities such as metabolic directions, clonal barcodes or paired surface proteins guide coupling construction and auxiliary readouts during training, whereas forward trajectory integration depends only on the initial cell states $X_0$, model time $t$ and the fitted network parameters.

#### S1.4 Model design, review and evaluation

Before implementing a new formulation, CellCompass prepares a scientific proposal that explains which biological relationship the existing dynamics cannot represent and how the new formulation addresses it. The proposal defines the state variables and governing equations, including assumptions about population change and stochasticity, and specifies the measurements used for fitting, how predictions will be generated, what inputs they require and which comparison will test the proposed improvement.

An independent reviewer agent assesses whether the available data support the proposed biological relation, whether the equations are consistent with the intended prediction and whether the planned comparison tests that relation. After implementation, a second review checks that the code, fitting objective and simulation procedure agree with the proposal. Each reviewer receives a task brief, the relevant proposal or code and the numerical results, and its assessment is returned to the main agent to guide revision.

Candidate models are compared using the same processed data, initial cells, reference methods and evaluation code, and CellCompass retains improved candidates during development. A candidate proceeds to further tuning only after meeting the feasibility and biological criteria. If a revision changes the biological relationship represented by the model, the proposal and evaluation plan are reviewed again before development continues.

Table 2: Evaluation and revision of a newly proposed CellCompass dynamics model.

| Step | Purpose | Evaluation | Next action |
|---|---|---|---|
| Scientific proposal | Define how the measurements constrain the dynamics. | Review the equations, assumptions, prediction inputs and planned comparison before implementation. | Implement the approved proposal or revise it following review. |
| Stage 1: feasibility | Establish stable fitting and valid trajectory generation. | Check finite losses, valid simulated states and acceptable population-prediction error. | Proceed if feasible, otherwise revise the implementation or formulation. |
| Stage 2: biological test | Test recovery of the biological relation that motivated the model. | Compare the prespecified biological measure with the strongest comparable reference method while assessing population agreement. | Proceed if the criterion is met, otherwise revise the model. |
| Stage 3: tuning | Improve population agreement while retaining the Stage 2 improvement. | Reduce Wasserstein-1 distance while monitoring the biological measure on the Stage 2 data. | Submit the selected model for final confirmation. |
| Final confirmation | Confirm the selected formulation before reporting results. | Repeat the complete evaluation with equations, prediction inputs and numerical settings fixed. | Retain the confirmed model or revisit the failed step. |

In Stage 2 (Table 2 and Supplementary Fig. 1), the primary biological measure must improve by at least 10% over the strongest comparable reference method without a substantial loss of future-population agreement. Stage 3 tuning ends when its performance target is met or the allowed number of trials is reached, and the selected model then proceeds to final confirmation before it is retained in the dynamical engine.

Supplementary Notes S2 and S3 describe the common mathematical framework and training procedures, Supplementary Note S4 presents DynBench100 and the agent comparisons, and Notes S5–S8 explain how CellCompass selected or developed models for the four single-cell studies, followed by their methods and evaluation. Researcher requests and excerpts from the model proposals are quoted where available, accompanied by summaries of the corresponding CellCompass outputs.

The researcher defines the biological question and can request additional analyses as a study develops. In the EMT study, a second request asked for an explanation of the differences between conditions (Supplementary Note S5), and the ProteinBridge follow-up requested downstream analyses and comparisons with reference methods (Supplementary Note S8). The LineageBridge example in Supplementary Note S7 follows the model proposal, reviewer feedback, code revisions, fitting and final evaluation within one development session.

##### S1.5 Language-model usage during algorithm development

We estimated the language-model usage over the full development of LineageBridge, from the initial research request through the 54 development trials to the final evaluation run (21.5 hours in total; Table 3). Design, implementation, review and analysis together used 125.24 million tokens, including 1.01 million output tokens, corresponding to an estimated cost of US$127 assuming that 116.6 million of the 124.2 million input tokens were cached at GPT-5.5 API rates of US$5, US$0.50 and US$30 per million uncached input, cached input and output tokens, respectively.

Table 3: Language-model use and estimated cost of developing LineageBridge.

| Quantity | Value |
|---|---:|
| Development time | 21.5 h |
| Model-development trials | 54 |
| Final training and evaluation run | 1 |
| Total tokens | 125.24 million |
| Output tokens | 1.01 million |
| Estimated API-equivalent cost | US$127 |

### S2 Mathematical framework for cell-state dynamics and population change

The dynamical engine represents cell-state transitions, stochastic variation and changes in effective abundance within a common population equation. Its model families differ in their treatment of these processes and in the constraints imposed between observed populations. Below, we describe the corresponding Fokker–Planck equation and variational formulations of optimal transport, Wasserstein–Fisher–Rao transport, Schrödinger bridges and regularized unbalanced optimal transport. Their finite-sample flow-matching implementations are given in Supplementary Note S3.

#### S2.1 Representing cell populations and their dynamics

**Empirical probability and positive measures.** Let $\mathcal{X} \subseteq \mathbb{R}^d$ be the state space used for one analysis. At experimental time $T_k$, observed cells $x_i^{(k)} \in \mathcal{X}$ and non-negative weights $a_i^{(k)}$ define the empirical positive measure

$$\widehat{\mu}_k = \sum_{i=1}^{n_k} a_i^{(k)} \delta_{x_i^{(k)}}, \qquad M_k = \widehat{\mu}_k(\mathcal{X}) = \sum_{i=1}^{n_k} a_i^{(k)}.$$

Because capture and sequencing efficiency can decouple observed cell counts $n_k$ from population abundance, balanced formulations normalize $\widehat{\mu}_k$ to a probability measure ($a_i^{(k)} = 1/n_k$, $M_k = 1$), whereas unbalanced formulations represent relative abundance changes through effective mass $M_k$.

When a density exists, we write $\mu_t(dx) = \rho_t(x)\,dx$. All equations below also have weak measure-valued forms. For a smooth compactly supported test function $\varphi$, a deterministic velocity $v_t$ and an effective-growth rate $g_t$ (the source per unit mass) satisfy

$$\frac{d}{dt}\int_{\mathcal{X}} \varphi(x)\,d\mu_t(x) = \int_{\mathcal{X}} \nabla\varphi(x)^{\mathsf{T}} v_t(x)\,d\mu_t(x) + \int_{\mathcal{X}} \varphi(x) g_t(x)\,d\mu_t(x). \tag{S2.1}$$

This weak form remains meaningful for empirical and singular measures. Setting $\varphi \equiv 1$ yields

$$\frac{dM_t}{dt} = \int g_t(x)\,d\mu_t(x),$$

so only the population average of $g_t$ controls total mass. Variation of the growth rate across cell states changes composition even when the net mass derivative is zero.

**A unified Fokker–Planck equation with growth.** Consider weighted particles obeying the Itô dynamics

$$dX_t = b_t(X_t)\,dt + \sigma\,dW_t, \qquad d\log w_t = g_t(X_t)\,dt, \tag{S2.2}$$

where $b_t$ is a drift, $\sigma \geq 0$ is a constant isotropic noise amplitude, $W_t$ is standard Brownian motion (denoted $B_t$ in the main Methods) and $w_t$ is an effective particle weight. Under standard integrability conditions, the weighted law $\mu_t(A) = \mathbb{E}[w_t \mathbf{1}\{X_t \in A\}]$ has density satisfying

$$\partial_t \rho_t(x) + \nabla\cdot\big(\rho_t(x) b_t(x)\big) = \frac{1}{2}\sigma^2 \Delta\rho_t(x) + g_t(x)\rho_t(x). \tag{S2.3}$$

Equation (S2.3) contains the principal families used in snapshot dynamics: setting $\sigma = 0$ and $g_t = 0$ gives balanced deterministic transport, $\sigma > 0$ with $g_t = 0$ gives a mass-conserving diffusion or stochastic bridge, $g_t \neq 0$ introduces unbalanced dynamics, and combining $\sigma > 0$ with $g_t \neq 0$ gives an unbalanced stochastic model.

For constant $\sigma$, define the current velocity

$$v_t = b_t - \frac{1}{2}\sigma^2 \nabla \log \rho_t$$

on the positive-density region. The diffusion equation can then be rewritten as $\partial_t \rho_t + \nabla \cdot (\rho_t v_t) = g_t \rho_t$. The current velocity combines the drift with the contribution of diffusion to population movement. Recovering the drift of the stochastic process therefore requires both the diffusion coefficient $\sigma^2/2$ and the score function $\nabla \log \rho_t$, even when the current velocity is known.

#### S2.2 Transport and bridge formulations

**Static balanced optimal transport.** For probability measures $\mu_0$ and $\mu_1$, let $\Pi(\mu_0, \mu_1)$ be the set of couplings with those marginals. The Kantorovich problem with cost $c$ is

$$\mathcal{T}_c(\mu_0, \mu_1) = \inf_{\pi \in \Pi(\mu_0, \mu_1)} \int_{\mathcal{X} \times \mathcal{X}} c(x, y)\, d\pi(x, y). \tag{S2.4}$$

For $c(x, y) = \|x - y\|_2^2$, this is the squared Wasserstein-2 distance. An endpoint coupling $\pi \in \Pi(\mu_0, \mu_1)$ specifies a statistical correspondence between the two populations, and its biological meaning depends on the transport cost $c$ and auxiliary constraints.

For finite measures with weights $a \in \mathbb{R}_+^n$ and $b \in \mathbb{R}_+^m$ of equal total mass, a commonly used entropic approximation is [1]

$$\min_{\Gamma \geq 0} \sum_{i,j} C_{ij} \Gamma_{ij} + \varepsilon \sum_{i,j} \Gamma_{ij} \left( \log \frac{\Gamma_{ij}}{a_i b_j} - 1 \right), \quad \Gamma \mathbf{1}_m = a, \quad \Gamma^\mathsf{T} \mathbf{1}_n = b. \tag{S2.5}$$

The regularization $\varepsilon$ spreads mass across endpoint pairs and permits solution by Sinkhorn scaling. Waddington-OT uses population couplings to infer transitions between time-resolved single-cell samples [2]. Here, coupled cells provide endpoint pairs for training continuous-time dynamics, as described in Supplementary Note S3.

**Benamou–Brenier dynamic transport.** For absolutely continuous probability measures with finite second moment, the Benamou–Brenier formulation represents squared Wasserstein-2 distance by minimum kinetic action [3]:

$$W_2^2(\mu_0, \mu_1) = \inf_{\rho, v} \left\{ \int_0^1 \int_{\mathcal{X}} \rho_t(x) \|v_t(x)\|_2^2 \, dx\, dt : \partial_t \rho_t + \nabla \cdot (\rho_t v_t) = 0,\ \rho_0 = \mu_0,\ \rho_1 = \mu_1 \right\}. \tag{S2.6}$$

Under suitable regularity, the minimizer is a constant-speed Wasserstein geodesic, and the static and dynamic formulations have the same optimal value, with the former determining an endpoint coupling and the latter a density path and momentum $m_t = \rho_t v_t$. In the convex momentum formulation, the cost $\|m_t\|^2/\rho_t$ is infinite for nonzero momentum at zero density.

Minimum kinetic action selects among solutions of the continuity equation. Additional biological information can change the inferred paths through the state representation, transport cost, measured velocities or intermediate population constraints.

**Wasserstein–Fisher–Rao and dynamic unbalanced transport.** When endpoint masses differ, the balanced coupling set is empty. Dynamic unbalanced transport augments the continuity equation with a source and penalizes both displacement and local mass change. One common scaling of the Wasserstein–Fisher–Rao (WFR) action is [4]

$$\mathrm{WFR}_\kappa^2(\mu_0, \mu_1) = \inf_{\rho, v, g} \int_0^1 \int_{\mathcal{X}} \rho_t(x) \left\{ \|v_t(x)\|_2^2 + \kappa^2 g_t(x)^2 \right\} \, dx \, dt, \tag{S2.7}$$

subject to

$$\partial_t \rho_t + \nabla \cdot (\rho_t v_t) = g_t \rho_t, \qquad \rho_0 = \mu_0, \quad \rho_1 = \mu_1.$$

The scale $\kappa$ sets the relative cost of reaction and transport. As the penalty on $g$ tends to infinity and the endpoint masses agree, admissible minimizers approach balanced transport under appropriate compactness conditions. At finite $\kappa$, removal and recreation can cost less than transport over a long state-space distance.

The WFR source describes changes in effective population mass. In the empirical models below, growth targets can be derived either from source-relaxed transport couplings, as in VGFM, or from WFR semi-couplings and their transport–growth paths, as in WFR-FM. Supplementary Note S3 describes these constructions and their endpoint-recovery properties.

**Schrödinger bridges.** Let $R^\sigma$ be the path measure of the reference diffusion $dX_t = b_t^0(X_t)dt + \sigma \, dW_t$, initialized at $\mu_0$, with constant isotropic noise amplitude $\sigma > 0$. Let $P_0$ and $P_1$ denote the endpoint marginals of a candidate path measure $P$. The dynamic Schrödinger problem is

$$\inf_{P:\, P_0 = \mu_0,\, P_1 = \mu_1} \mathrm{KL}(P \,\|\, R^\sigma). \tag{S2.8}$$

It selects the path law closest in relative entropy to the reference process while exactly matching the endpoint distributions [5]. If $P$ has controlled drift $b_t^0 + u_t$ and the same diffusion amplitude, Girsanov's theorem gives, under the usual absolute-continuity and integrability conditions,

$$\mathrm{KL}(P \,\|\, R^\sigma) = \mathbb{E}_P \int_0^1 \frac{1}{2\sigma^2} \|u_t(X_t)\|_2^2 \, dt, \quad \partial_t \rho_t + \nabla \cdot \{\rho_t (b_t^0 + u_t)\} = \frac{1}{2} \sigma^2 \Delta \rho_t.$$

The bridge therefore preserves a stochastic reference and changes it minimally to satisfy the endpoint constraints. The reference drift can encode prior dynamics when such a prior is scientifically justified [6].

Schrödinger bridges connect entropic transport, stochastic control and score-based modeling. For a Brownian reference with zero drift, their limit as $\sigma \to 0$ recovers quadratic optimal transport under suitable scaling and regularity. At nonzero noise ($\sigma > 0$), rewriting the control action using current velocity produces a kinetic term together with a Fisher-information contribution and boundary-entropy terms, distinguishing a Schrödinger bridge from deterministic dynamic OT with added Gaussian noise. Simulation-free Schrödinger bridges via score and flow matching (SF2M) learns velocity fields and score-based diffusion terms from endpoint couplings and tractable conditional bridges to model stochastic dynamics while conserving total population mass [7].

**Regularized unbalanced optimal transport (RUOT).** RUOT combines displacement, diffusion and mass change [8, 9]. For constant isotropic noise amplitude $\sigma$, the density satisfies

$$\partial_t \rho_t + \nabla \cdot (\rho_t b_t) = \frac{1}{2} \sigma^2 \Delta \rho_t + g_t \rho_t, \qquad \rho_0 = \mu_0, \quad \rho_1 = \mu_1. \tag{S2.9}$$

The corresponding RUOT action is

$$\mathcal{L}_{\mathrm{RUOT}}(\rho, b, g) = \int_0^1 \int_{\mathcal{X}} \rho_t(x) \left\{ \frac{1}{2} \|b_t(x)\|_2^2 + \alpha \Psi\big(g_t(x)\big) \right\} dx\, dt, \tag{S2.10}$$

where $\Psi : \mathbb{R} \to [0, +\infty]$ is convex and $\alpha > 0$ weights the cost of effective population growth relative to state movement. Choosing $\Psi(g) = +\infty$ for $g \neq 0$ enforces mass conservation and recovers a balanced stochastic bridge formulation under the stated diffusion and reference-process convention. When $\sigma \to 0$ and $\Psi(g) = g^2$ up to scale, equations (S2.9)–(S2.10) reduce to dynamic unbalanced transport with a WFR action. Imposing $g = 0$ as well, with equal endpoint mass, gives Benamou–Brenier dynamic transport. Other convex source penalties, reference birth–death laws, relaxed endpoint marginals and explicit Fisher regularization define related RUOT variants.

#### S2.3 Connections among the implemented model families

**Finite-data implementation.** In the dynamical engine, these formulations determine how cells are coupled across observation times and how paths are constructed between coupled cells. Training samples intermediate states along these paths and uses their known rates of change to learn the velocity field, together with growth fields and score-based diffusion terms where required. The learned fields then generate trajectories from the initial population. This simulation-free conditional flow-matching framework decomposes the continuous variational problems in equations (S2.6), (S2.7), (S2.8) and (S2.10) into finite-sample endpoint coupling followed by conditional vector-field regression along tractable bridge paths, avoiding trajectory integration during training.

Table 4: Couplings, learned fields and conditional paths in the implemented flow-matching models. Finite-sample constructions and conditions for population recovery are given in Supplementary Note S3.

| Model | Endpoint coupling | Learned fields and simulation | Conditional paths |
|---|---|---|---|
| Balanced OT conditional flow matching (OT-CFM) | Balanced coupling with both endpoint marginals fixed | Velocity field with deterministic ODE simulation and fixed particle weights | Linear state interpolation with constant mass |
| SF2M | Balanced coupling (entropic or exact OT) with both endpoint marginals fixed | Current velocity and score-based diffusion term with stochastic or probability-flow simulation and fixed particle weights | Conditional Brownian bridges with constant mass |
| VGFM | Unbalanced coupling with the source marginal relaxed and target marginal fixed | Velocity and growth fields with deterministic weighted-particle ODE simulation | Linear state interpolation with log-linear mass interpolation |

| Model | Endpoint coupling | Learned fields and simulation | Conditional paths |
|---|---|---|---|
| WFR-FM | Entropy-transport plan with both marginals relaxed, converted to two semi-couplings that match the source and target masses, respectively | Velocity and growth fields with deterministic weighted-particle ODE simulation | WFR transport paths with quadratic mass interpolation |
| CRUFM | Unbalanced coupling with the source marginal relaxed and target marginal fixed, as in VGFM | Current velocity, score-based diffusion and growth fields with stochastic or weighted probability-flow simulation | Conditional Brownian bridges with log-linear mass interpolation |

Flow matching is the training procedure shared by these models, while the transport formulation determines the coupling and the paths used for training: VGFM learns deterministic changes in cell state and effective abundance, WFR-FM relates the costs of these changes through a Wasserstein–Fisher–Rao endpoint problem, and CRUFM additionally uses score matching to learn the score-based diffusion term for stochastic dynamics. For each family, exact recovery refers to the population path defined by its chosen coupling and conditional paths (Table 4).

**Relations among transport, bridge and flow-matching formulations.** Table 5 summarizes the limiting relationships that connect these variational transport, stochastic bridge and flow-matching formulations.

Table 5: Relations among transport, bridge and flow-matching formulations. Conditions specific to each relation are listed below, in addition to the regularity and existence assumptions of the cited results.

| Relation | Mathematical statement | Conditions |
|---|---|---|
| Static OT $\leftrightarrow$ dynamic OT | Quadratic Kantorovich transport and Benamou–Brenier kinetic action have the same minimum value [3] | Equal endpoint mass, finite second moments and squared Euclidean transport cost over a unit time interval |
| WFR $\to$ OT | As the penalty on mass change increases without bound, WFR approaches balanced dynamic OT [4] | Equal endpoint mass and suitable compactness. In equation (S2.7), $\kappa \to \infty$ while the transport cost remains fixed |
| Schrödinger bridge $\to$ OT | The small-noise limit recovers quadratic dynamic OT [5, 6] | Brownian reference with zero drift, fixed endpoint distributions and the appropriate rescaling of the entropy objective as $\sigma \to 0$ |

| Relation | Mathematical statement | Conditions |
|---|---|---|
| SF2M $\rightarrow$ Schrödinger bridge marginals | Flow matching and score matching recover the bridge marginals in the population exact-fit limit [7] | Brownian conditional bridges and an entropic endpoint coupling matched to the same reference diffusion. The score function is scaled by the diffusion coefficient when constructing the stochastic drift |
| RUOT $\rightarrow$ WFR, Schrödinger bridge or OT | The RUOT action reduces to WFR at zero diffusion with a quadratic source penalty, to a balanced bridge with zero source, and to dynamic OT when both vanish [8] | WFR: $\sigma = 0$ and $\Psi(g) = g^2$, with coefficients rescaled. Bridge: $g = 0$, $\sigma > 0$ and a zero-drift Brownian reference. OT: $g = 0$ and $\sigma = 0$. The balanced cases require equal endpoint mass |

### S3 Flow-matching training and trajectory generation

Single-cell time courses sample different cells at successive times, so the measured data do not directly provide trajectories for training. The dynamical engine first estimates a coupling between adjacent populations, then constructs conditional paths between the coupled cells. The states and rates of change along these paths provide training examples for the velocity field and, where required, the growth field and score-based diffusion term. Once trained, these fields generate trajectories from the initial cells.

#### S3.1 Learning population dynamics from coupled cells

**Finite endpoint coupling and model time.** Consider adjacent observations at model times $T_k < T_{k+1}$, with interval length $\Delta_k = T_{k+1} - T_k$. Time may be represented by elapsed physical time or by an ordinal index, and the convention used for each application is stated in its case-specific note. Write

$$\widehat{\mu}_k = \sum_{i=1}^{n_k} a_i \delta_{x_i}, \qquad \widehat{\mu}_{k+1} = \sum_{j=1}^{n_{k+1}} b_j \delta_{y_j}.$$

A non-negative matrix $\Gamma = (\Gamma_{ij})$ assigns mass or sampling weight to endpoint pairs $(x_i, y_j)$. In balanced flow matching,

$$\Gamma \mathbf{1} = a, \qquad \Gamma^{\mathsf{T}} \mathbf{1} = b. \tag{S3.1}$$

VGFM and CRUFM use a source-relaxed, target-constrained UOT problem:

$$\min_{\Gamma \geq 0} \langle C, \Gamma \rangle + \varepsilon \mathcal{H}(\Gamma) + \tau_{\mathrm{s}} \, \mathrm{KL}(\Gamma \mathbf{1} \,\|\, a), \qquad \Gamma^{\mathsf{T}} \mathbf{1} = b, \tag{S3.2}$$

where $\mathcal{H}$ is the entropic regularizer. The row marginal $r_i = \sum_j \Gamma_{ij}$ may differ from $a_i$, whereas the target column marginal is fixed. The analyses use unit empirical reference masses within a transport block.

Training uses a local coordinate $\tau \in [0, 1]$, while the neural fields receive model time

$$t = T_k + \tau \Delta_k.$$

We denote targets in local time by $\widetilde{U}_\tau$ and $\widetilde{G}_\tau$. The corresponding targets per unit model time, used in the main Methods, are $U_\tau = \widetilde{U}_\tau / \Delta_k$ and $G_\tau = \widetilde{G}_\tau / \Delta_k$. For multi-time-point series ($T_0 < \cdots < T_K$), couplings and conditional paths are constructed within each interval $[T_k, T_{k+1}]$, while a single time-conditioned neural field is trained jointly across all intervals.

**Conditional paths and population regression.** Let $E$ denote a sampled pair of cells at adjacent times, together with any mass information assigned to their connection. For a given pair $E = e$, $p_\tau(z \mid e)$ describes the distribution of intermediate states and $q_\tau(e) > 0$ gives the mass carried by that connection relative to its sampling weight. Suppose this conditional path obeys

$$\partial_\tau \{q_\tau(e) p_\tau(z \mid e)\} + \nabla \cdot \{q_\tau(e) p_\tau(z \mid e) u_\tau(z \mid e)\} = q_\tau(e) p_\tau(z \mid e) \gamma_\tau(e), \tag{S3.3}$$

with $\gamma_\tau(e) = \partial_\tau \log q_\tau(e)$. The field $u_\tau$ is the conditional current velocity, and for a smooth stochastic path, adding its score-based diffusion term gives the forward drift. If $E$ is sampled from a reference measure $\eta(de)$, the marginal positive measure is

$$\rho_\tau(z) = \int q_\tau(e) p_\tau(z \mid e) \, \eta(de). \tag{S3.4}$$

**Proposition S3.1 (population minimizers).** Assume finite second moments, $\rho_\tau(z) > 0$, and sufficient regularity of $(q_\tau, p_\tau, u_\tau, \gamma_\tau)$ to permit differentiation under the integral sign. The minimizers of the mass-weighted squared losses

$$\mathcal{L}_v(v) = \mathbb{E}\big[q_\tau(E)\|v(Z_\tau, \tau) - u_\tau(Z_\tau \mid E)\|_2^2\big]\,, \tag{S3.5}$$
$$\mathcal{L}_g(g) = \mathbb{E}\big[q_\tau(E)\{g(Z_\tau, \tau) - \gamma_\tau(E)\}^2\big] \tag{S3.6}$$

are the mass-weighted conditional expectations

$$v^*(z, \tau) = \frac{\int q_\tau(e)p_\tau(z \mid e)u_\tau(z \mid e)\,\eta(de)}{\rho_\tau(z)}, \tag{S3.7}$$
$$g^*(z, \tau) = \frac{\int q_\tau(e)p_\tau(z \mid e)\gamma_\tau(e)\,\eta(de)}{\rho_\tau(z)}. \tag{S3.8}$$

The mixture in equation (S3.4) satisfies

$$\partial_\tau \rho_\tau + \nabla \cdot (\rho_\tau v^*) = \rho_\tau g^*. \tag{S3.9}$$

*Proof.* At a fixed state and time, write $d\nu_z(e) = q_\tau(e)p_\tau(z \mid e)\eta(de)$. Its total mass is $\rho_\tau(z)$. If $\bar{u} = \int u_\tau(z \mid e)d\nu_z(e)/\rho_\tau(z)$, completing the square gives

$$\int \|v - u_\tau\|^2 d\nu_z = \rho_\tau(z)\|v - \bar{u}\|^2 + \int \|u_\tau - \bar{u}\|^2 d\nu_z.$$

The second term is independent of $v$, so the minimum is attained at $v = \bar{u}$. Applying the same calculation to the scalar growth target gives $g^*$ in equation (S3.8).

For a smooth compactly supported test function $\varphi$, the conditional continuity equation implies

$$\frac{d}{d\tau}\int \varphi(z)q_\tau(e)p_\tau(z \mid e)\,dz = \int \{\nabla\varphi \cdot u_\tau + \varphi\gamma_\tau\}q_\tau(e)p_\tau(z \mid e)\,dz.$$

Integrating over $e$, with differentiation and integration interchangeable, replaces the conditional targets by their mass-weighted means. Hence

$$\frac{d}{d\tau}\int \varphi\rho_\tau = \int (\nabla\varphi \cdot v^* + \varphi g^*)\rho_\tau,$$

which is the weak form of equation (S3.9). □

At each state and time, the optimal fields are mass-weighted averages of the conditional velocity and growth targets, and neural-network fitting approximates these conditional expectations over the training distribution.

**Proposition S3.2 (population endpoint recovery under exact fitting).** Suppose the conditional mixture has $\rho_0 = \widehat{\mu}_k$ and $\rho_1 = \widehat{\mu}_{k+1}$. If the marginal continuity equation with fields $(v^*, g^*)$ has a unique solution in the considered class, simulation of that population law from $\widehat{\mu}_k$ with the exact fields recovers $\widehat{\mu}_{k+1}$ at $T_{k+1}$. A weighted-particle realization uses

$$\frac{dX_t}{dt} = \frac{v^*(X_t, \tau)}{\Delta_k}, \qquad \frac{d\log w_t}{dt} = \frac{g^*(X_t, \tau)}{\Delta_k} \tag{S3.10}$$

for deterministic families, where $\tau = (t - T_k)/\Delta_k$ and $(v^*, g^*)$ are the local-time fields above. Stochastic families add the diffusion and score-based diffusion term specified by their conditional bridge.

*Proof.* Proposition S3.1 makes the conditional mixture a solution of the population equation. It has the specified initial measure, so uniqueness identifies it with the simulated law. In particular, their terminal measures agree.

In model time, write $v_t(x) = v^*(x, \tau)/\Delta_k$ and $g_t(x) = g^*(x, \tau)/\Delta_k$. For the deterministic particle representation, the product rule gives

$$\frac{d}{dt}\{w_t\varphi(X_t)\} = w_t\{\nabla\varphi(X_t) \cdot v_t(X_t) + \varphi(X_t)g_t(X_t)\}.$$

Averaging over initial particles yields the weak continuity equation. For a stochastic realization with smooth positive density, constant diffusion amplitude $\beta$, drift $b$ and the same weight equation, Itô's formula adds $\beta^2\Delta\varphi/2$. Its population equation is therefore

$$\partial_t\rho = -\nabla \cdot (\rho b) + \frac{\beta^2}{2}\Delta\rho + g\rho.$$

When $b = v + \beta^2\nabla \log \rho/2$, the score contribution satisfies $\nabla \cdot (\rho\beta^2\nabla \log \rho/2) = \beta^2\Delta\rho/2$, which cancels the diffusion term and leaves the same marginal continuity equation with current velocity $v$. □

**Balanced OT conditional flow matching (OT-CFM).** Balanced conditional flow matching [10, 11] samples $(X_0, X_1)$ from a coupling satisfying equation (S3.1). For the default deterministic linear path,

$$Z_\tau = (1 - \tau)X_0 + \tau X_1, \qquad \widetilde{U}_\tau = X_1 - X_0, \qquad q_\tau = 1, \qquad \widetilde{G}_\tau = 0. \tag{S3.11}$$

The velocity field, expressed in model time, minimizes

$$\mathcal{L}_{\mathrm{FM}}(\theta) = \mathbb{E}\left[\left\|v_\theta(Z_\tau, T_k + \tau\Delta_k) - \frac{X_1 - X_0}{\Delta_k}\right\|_2^2\right]. \tag{S3.12}$$

Since the coupling has the observed marginals and the path reaches its paired endpoints, Propositions S3.1–S3.2 give marginal endpoint recovery in the population exact-fit limit.

A residual conditional path can change the route without changing this endpoint statement. For example,

$$Z_\tau = (1 - \tau)X_0 + \tau X_1 + \tau(1 - \tau)R, \quad \widetilde{U}_\tau = X_1 - X_0 + (1 - 2\tau)R.$$

The added term is zero at $\tau = 0, 1$, so it changes the intermediate path while leaving both endpoint states unchanged. PulseBridge uses this construction to incorporate metabolic RNA directions after removing components associated with day-0 labeling differences (Supplementary Note S6).

#### S3.2 VGFM: unbalanced deterministic flow matching

VGFM uses equation (S3.2) [12]. Put $r_i = \sum_j \Gamma_{ij}$ and assume $a_i, r_i > 0$. Conditional on source $i$, the target is sampled with probability $P(j \mid i) = \Gamma_{ij}/r_i$. A source particle starts with mass $a_i$ and receives terminal multiplier

$$m_i = \frac{r_i}{a_i}, \qquad \ell_i = \log m_i.$$

VGFM uses the linear state path in equation (S3.11), a log-linear mass path $w_i(\tau) = m_i^\tau$, and the local targets

$$\widetilde{U}_{ij,\tau} = y_j - x_i, \qquad \widetilde{G}_{i,\tau} = \frac{d}{d\tau}\log w_i(\tau) = \ell_i. \tag{S3.13}$$

Edges are sampled in proportion to $\Gamma_{ij}$, and the regression loss is multiplied by

$$\omega_{ij}(\tau) = m_i^{\tau-1}. \tag{S3.14}$$

This weight makes the effective edge mass follow

$$q_{ij}(\tau) = \Gamma_{ij} m_i^{\tau-1} = \Gamma_{ij} \left(\frac{a_i}{r_i}\right)^{1-\tau}. \tag{S3.15}$$

At $\tau = 0$, $\sum_j q_{ij}(0) = a_i$. At $\tau = 1$, $q_{ij}(1) = \Gamma_{ij}$ and $\sum_i q_{ij}(1) = b_j$ because the target is constrained. This gives the endpoint-mass recovery result for the source-relaxed convention.

The velocity and growth networks are trained jointly to reproduce the rates of state change and mass change along these paths, using the objective

$$\begin{aligned}\mathcal{L}_{\mathrm{VGFM}}(\theta, \phi) = \mathbb{E}_{(i,j),\tau}\Bigg[\omega_{ij}(\tau)\Bigg\{ \left\| v_\theta(Z_\tau, t) - \frac{y_j - x_i}{\Delta_k} \right\|_2^2 \\ + \lambda_g \left( g_\phi(Z_\tau, t) - \frac{\ell_i}{\Delta_k} \right)^2 \Bigg\}\Bigg].\end{aligned} \tag{S3.16}$$

The coefficient $\lambda_g$ weights the growth error relative to the velocity error. VGFM simulates equation (S3.10) from the observed starting cells and their specified initial weights using the fitted velocity and growth fields, which under exact fitting realize the UOT-induced mixture in equation (S3.15).

#### S3.3 WFR-FM: semi-coupling and traveling-Gaussian targets

WFR-FM [13] differs from VGFM in both its endpoint problem and its conditional mass path. For scale $\delta > 0$, define the truncated WFR endpoint cost

$$c_\delta(x, y) = -2 \log \cos_+ \left( \frac{\|x - y\|_2}{2\delta} \right).$$

Here $\cos_+(a) = \cos(a)$ for $0 \leq a < \pi/2$ and is zero for $a \geq \pi/2$. In computation, the angle is kept below $\pi/2$ by a small positive margin $\epsilon_{\mathrm{ang}}$. With entropic coefficient $\varepsilon$ and marginal coefficients $\tau_0, \tau_1 > 0$, the finite optimal entropy-transport (OET) problem has the form

$$\min_{\gamma \geq 0} \langle c_\delta, \gamma \rangle + \varepsilon \mathcal{H}(\gamma) + \tau_0 \, \mathrm{KL}_+(\gamma \mathbf{1} \,\|\, a) + \tau_1 \, \mathrm{KL}_+(\gamma^\top \mathbf{1} \,\|\, b). \tag{S3.17}$$

The generalized Kullback–Leibler divergence $\mathrm{KL}_+(r \| a) = \sum_i [r_i \log(r_i/a_i) - r_i + a_i]$ penalizes differences between non-negative masses. Thus both marginals are relaxed in the OET plan. If $\gamma$ is its solution with row and column sums $\bar{\gamma}_0$ and $\bar{\gamma}_1$, it is converted into source- and target-side semi-couplings

$$\Gamma^0_{ij} = \gamma_{ij} \frac{a_i}{\bar{\gamma}_{0,i}}, \qquad \Gamma^1_{ij} = \gamma_{ij} \frac{b_j}{\bar{\gamma}_{1,j}}. \tag{S3.18}$$

They satisfy $\Gamma^0 \mathbf{1} = a$ and $(\Gamma^1)^\top \mathbf{1} = b$. WFR-FM samples an edge from $\Gamma^0$ and assigns the pair-specific terminal mass

$$m_1^{ij} = \frac{\Gamma^1_{ij}}{\Gamma^0_{ij}}. \tag{S3.19}$$

This two-semi-coupling construction, rather than the row sum of a single UOT plan, is what connects the finite problem to dynamic WFR [4, 13].

For one sampled edge, let $d = \|y-x\|_2$, $e = (y-x)/d$ for $d > 0$ (with $e = 0$ when $x = y$), and

$$\alpha = \min\left\{\frac{d}{2\delta}, \frac{\pi}{2} - \epsilon_{\text{ang}}\right\}, \quad \tau_d = \tan\alpha, \quad s = \sqrt{\frac{m_1}{1+\tau_d^2}}. \tag{S3.20}$$

Define

$$A = 1 + m_1 - 2s, \qquad B = 1 - s, \qquad \Omega = 2\delta\tau_d s\, e, \qquad m(\tau) = A\tau^2 - 2B\tau + 1. \tag{S3.21}$$

Then $m(0) = 1$ and $m(1) = m_1$. The traveling center and conditional targets are

$$\eta_\tau = x + \Omega \int_0^\tau \frac{du}{m(u)}, \qquad Z_\tau = \eta_\tau + \sigma\epsilon, \tag{S3.22}$$

$$\widetilde{U}_\tau = \frac{\Omega}{m(\tau)}, \qquad \widetilde{G}_\tau = \frac{2A\tau - 2B}{m(\tau)} = \frac{d}{d\tau}\log m(\tau). \tag{S3.23}$$

Here $\epsilon \sim \mathcal{N}(0, I)$ and $\sigma$ is the standard deviation of the Gaussian perturbation; the deterministic WFR-FM model sets $\sigma = 0$, whereas a nonzero constant $\sigma$ in equation (S3.22) yields Gaussian-smoothed endpoint measures. The integral has an arctangent closed form, so the training target is evaluated without numerical integration. The WFR-FM loss weights both velocity and growth regression by $m(\tau)$, and targets per unit model time are obtained by dividing equation (S3.23) by $\Delta_k$.

**Proposition S3.3 (WFR-FM exact-fit recovery, adapted from [13]).** Assume the optimal-entropy solution is converted exactly to semi-couplings, $\sigma = 0$, no positive-mass edge activates the angle clip, the conditional rays are defined by equations (S3.19)–(S3.23), and the population velocity and growth regressions attain their minima under the realization conditions of Proposition S3.2. Then their marginal fields solve the unbalanced continuity equation of the mixture of WFR rays, whose endpoints are $\widehat{\mu}_k$ and $\widehat{\mu}_{k+1}$.

*Proof.* Without angle clipping, write $\alpha = \|y-x\|/(2\delta)$ and $s = \sqrt{m_1}\cos\alpha$. The mass path can be written as

$$m(\tau) = \{1 - \tau + s\tau\}^2 + \{\sqrt{m_1}\sin\alpha\,\tau\}^2.$$

For positive endpoint mass and $0 < \alpha < \pi/2$, this expression stays positive. Let $\phi(\tau)$ be the continuous angle of the two-dimensional vector inside these squares. It starts at zero and ends at $\alpha$, and differentiation gives

$$\phi'(\tau) = \frac{\sqrt{m_1}\sin\alpha}{m(\tau)}.$$

Since $\Omega = 2\delta\sqrt{m_1}\sin\alpha\, e$, the traveling center is $x + 2\delta\phi(\tau)e$ and therefore reaches $y$ at $\tau = 1$. At $\alpha = 0$, both centers coincide and the mass path is the positive square $(1 - \tau + \sqrt{m_1}\tau)^2$.

Each edge contributes mass $\Gamma^0_{ij}$ initially and $\Gamma^0_{ij} m_1^{ij} = \Gamma^1_{ij}$ finally, so summing these contributions recovers the two observed endpoint measures by the semi-coupling constraints. Differentiating the center and log mass gives the conditional velocity and growth targets, and Propositions S3.1 and S3.2 then give the marginal equation and endpoint recovery. □

The scale $\delta$ is estimated from latent distances between adjacent observations. In practice, angle clipping ($\alpha < \pi/2 - \epsilon_{\text{ang}}$), Gaussian smoothing ($\sigma > 0$) and blockwise entropic regularization make endpoint recovery approximate, and the resulting deviations are assessed with Wasserstein-1 distance and total-mass variation.

##### S3.4 SF2M: Schrödinger bridges via score and flow matching

Simulation-free Schrödinger bridges via score and flow matching (SF2M) [7] is a method for solving Schrödinger-bridge problems without simulating the learned dynamics during training. It combines an entropy-regularized endpoint coupling matched to the reference diffusion with score matching and flow matching. The coupling links cells across observation times, and conditional Brownian bridges provide intermediate training states between the paired endpoints, with total population mass held constant. The dynamical engine also implements the exact-OT minibatch approximation described by Tong et al. [11].

The stochastic formulas below use a unit model-time interval. For an interval of length $\Delta_k$, a Brownian bridge with local noise amplitude $\sigma$ has model-time diffusion amplitude $\beta_k = \sigma/\sqrt{\Delta_k}$. Its current-velocity target is $U_\tau = \widetilde{U}_\tau/\Delta_k$ and its score-based diffusion term is $\beta_k^2 \nabla \log \rho/2$. The denoising scale is correspondingly $2\sigma_\tau/\beta_k^2$. We therefore rescale the velocity target, diffusion amplitude and score-based diffusion term together when changing the model-time interval.

For a paired source cell $x$ and target cell $y$, the intermediate training state is sampled as

$$\mu_\tau = (1-\tau)x + \tau y, \qquad \sigma_\tau = \sigma\sqrt{\tau(1-\tau)}, \qquad Z_\tau = \mu_\tau + \sigma_\tau \epsilon, \quad \epsilon \sim \mathcal{N}(0, I). \tag{S3.24}$$

Here $\tau$ is the relative position between the two observation times, and $\sigma_\tau$ is the noise standard deviation around the interpolated mean; this standard deviation vanishes at both endpoints and is largest halfway between them, with its overall amplitude set by $\sigma$. The conditional local velocity is

$$\widetilde{U}_\tau = y - x + \frac{1-2\tau}{2\tau(1-\tau)}(Z_\tau - \mu_\tau), \tag{S3.25}$$

with a small numerical denominator stabilization near the endpoints. The velocity field is fitted to $U_\tau = \widetilde{U}_\tau/\Delta_k$ by minimizing squared prediction error. Both the intermediate states and their targets are sampled directly, without integrating the learned dynamics during training.

The score function, $\nabla_z \log p_\tau(z)$, describes how the log density of intermediate states changes with cell state. The engine represents the corresponding score-based diffusion term in the stochastic drift as $s_\psi(z,t) = \nabla_z F_\psi(z,t)$, where $F_\psi$ is a scalar potential, and this term includes the factor $\sigma^2/2$ multiplying the score function. For $\sigma > 0$, the denoising score-matching loss uses the scale

$$\lambda(\tau) = \frac{2\sigma_\tau}{\sigma^2}. \tag{S3.26}$$

For SF2M, the score model minimizes

$$\mathcal{L}_s(\psi) = \mathbb{E}\left[\|\lambda(\tau)s_\psi(Z_\tau, t) + \epsilon\|_2^2\right], \tag{S3.27}$$

with the potential regularizer added during fitting. By the Gaussian denoising identity, minimizing $\mathcal{L}_s(\psi)$ yields the population optimal score field

$$s^*(z,t) = \frac{\sigma^2}{2}\nabla \log p_\tau(z). \tag{S3.28}$$

After fitting, stochastic trajectories are generated from initial cells using

$$dX_t = \{v_\theta(X_t,t) + s_\psi(X_t,t)\}\,dt + \sigma\,dW_t. \tag{S3.29}$$

SF2M keeps particle weights constant. For the unit model-time interval used in these equations, the simulation noise amplitude $\sigma$ is the same as in the conditional bridge, and for a general interval the diffusion amplitude and score-based diffusion term use the conversion given above. Adding $s_\psi$ to the current velocity $v_\theta$ gives the stochastic drift, whose contribution offsets diffusion in the population equation and so preserves the fitted marginal path. The corresponding deterministic probability-flow model uses $dX_t = v_\theta(X_t,t)dt$, with both $s_\psi$ and diffusion omitted.

##### S3.5 CRUFM: Conditional regularized unbalanced flow matching

CRUFM combines the same stochastic path with the source-relaxed UOT coupling and effective-growth learning used by VGFM. The transported mass of source cell $i$ relative to its initial mass is $m_i = r_i/a_i$, giving the growth target $\log m_i/\Delta_k$. The velocity, growth and score-matching losses are weighted by $\omega_{ij}(\tau) = m_i^{\tau-1}$, as in equation (S3.14); for score matching, the weight multiplies the squared error inside the expectation in equation (S3.27), so the score relationship in equation (S3.28) applies to the normalized, mass-weighted density of intermediate states.

During simulation, states follow equation (S3.29), while particle weights evolve according to $d\log w_t = g_\phi(X_t, t)\,dt$, so the predicted population can expand or decline as its state distribution changes. At zero noise, CRUFM uses the deterministic VGFM path and disables score matching, avoiding the singular scaling in equation (S3.26). As for VGFM, endpoint recovery requires the coupling and growth targets to use the same mass convention.

### S4 DynBench100 and agent evaluation

#### S4.1 Construction of DynBench100

DynBench100 evaluates whether a model-design system can recover multiple properties of a cellular dynamical process from time-resolved snapshots. Each scenario is simulated from a stochastic gene-regulatory network with state-dependent population change, and the simulator provides six classes of ground truth: cell velocities, growth-driver genes, populations at withheld time points, initial-cell fates, perturbation responses and the signed regulatory network. Agents receive only the training snapshots, task description and a common terminal-state classifier, while the simulator equations and ground-truth quantities are withheld for evaluation.

**Regulatory dynamics and population change.** For each cell, the non-negative state $x_i$ represents the expression of gene $i$. Expression evolves according to

$$\mathrm{d}x_i = v_i(x)\,\mathrm{d}t + \sigma\,\mathrm{d}W_i, \qquad v_i(x) = a_i F_i(x) - d_i x_i, \tag{S4.1}$$

where $a_i$ is the production rate, $d_i$ is the degradation rate and the $W_i$ are independent Wiener processes. Regulatory inputs are combined multiplicatively,

$$F_i(x) = \prod_{j\in\mathcal{A}_i} \frac{x_j^{n_{ji}}}{K_{ji}^{n_{ji}} + x_j^{n_{ji}}} \prod_{j\in\mathcal{R}_i} \frac{K_{ji}^{n_{ji}}}{K_{ji}^{n_{ji}} + x_j^{n_{ji}}},$$

where $\mathcal{A}_i$ and $\mathcal{R}_i$ are the activators and repressors of gene $i$, $K_{ji}$ is the half-saturation constant and $n_{ji}$ is the Hill coefficient. For a gene without regulatory inputs, $F_i = 0.5$.

Each toggle module contains two self-activating transcription factors that inhibit one another; their relative expression defines a binary fate choice, so $m$ modules specify $2^m$ possible terminal fates, and each factor also activates one reporter gene. Confounder genes receive inputs from between two and all available reporters and degrade more slowly, allowing them to correlate with time without directly controlling growth, whereas noise genes have constitutive production and no regulatory connections. Production and degradation rates $(a_i, d_i)$ were $(1.2, 0.3)$ for transcription factors, $(0.8, 0.25)$ for reporters, $(0.6, 0.12)$ for confounders and $(0.5, 0.3)$ for noise genes. Toggle-edge $K$ and Hill coefficients were sampled uniformly from $[0.45, 0.55]$ and $[2.7, 3.3]$, whereas reporter and confounder edges had Hill coefficient 2 and $K$ sampled from $[0.25, 0.35]$ and $[0.17, 0.23]$, respectively.

Effective growth depended on one reporter or the mean expression of two reporters associated with the first toggle module. With $G$ denoting this set of growth-driver genes, the rate was

$$g(x) = -0.02 + \frac{0.06}{1 + \exp[-5(\bar{x}_G - 0.3)]}, \qquad \bar{x}_G = \frac{1}{|G|}\sum_{j\in G} x_j. \tag{S4.2}$$

Cells were advanced by Euler–Maruyama integration with time step $\Delta t = 0.1$, with negative expression values set to zero. At each step, a cell divided with probability $\min(0.1, \max(g, 0)\Delta t)$ or was removed with probability $\min(0.1, \max(-g, 0)\Delta t)$. A daughter inherited its parent's state with independent Gaussian noise of standard deviation $0.3\sigma$. Initial expression was sampled with standard deviation 0.12 around means of 0.5 for transcription factors and noise genes, 0.3 for reporters and 0.2 for confounders, again truncated at zero.

**Scenario settings and ground truth.** The 100 scenarios comprise 20 easy, 40 medium, 25 hard and 15 adversarial cases, which vary in network size, noise, initial population size and sampling times (Table 6); increasing difficulty introduces more toggle modules, confounders and noise genes, together with greater stochastic variation. Each scenario was sampled at 6–12 equally spaced simulation times, with two or three intermediate snapshots withheld in their entirety to evaluate temporal interpolation between observed time points, and scenario-specific seeds determine the regulatory parameters and connections, initial cells, stochastic fluctuations and population events.

Table 6: DynBench100 simulation settings. Ranges describe variation among scenarios within each group. The simulated time span is measured in model time units. Withheld time points are included in the sampled total and are used only for evaluation. Fate counts denote the alternatives specified by the toggle modules.

| Setting | Easy | Medium | Hard | Adversarial |
|---|---|---|---|---|
| Scenarios | 20 | 40 | 25 | 15 |
| Genes | 5–7 | 10–18 | 18–28 | 25–36 |
| Terminal fates | 2 | 4 or 8 | 8 or 16 | 8 or 16 |
| Confounder genes | 1–2 | 2–4 | 4–6 | 7–10 |
| Noise genes | 0–1 | 0–2 | 2–6 | 6–10 |
| Initial cells | 650–850 | 800–1,100 | 900–1,300 | 900–1,500 |
| Noise amplitude $\sigma$ | 0.08–0.12 | 0.11–0.15 | 0.15–0.19 | 0.18–0.22 |
| Simulated time span (model units) | 9–10 | 11–13 | 13–15 | 15–18 |
| Number of sampled time points | 6–7 | 8–9 | 9–11 | 10–12 |
| Number of withheld time points | 2 | 2 | 2–3 | 2–3 |

The simulator records the drift $v(x)$ at each sampled cell, the identities of the genes in $G$ and the signed regulatory edges, while populations at withheld times provide the reference distributions. Root-cell fate probabilities were estimated from 50 stochastic trajectories per initial cell, simulated without birth or death events and classified by the terminal toggle states. For each gene perturbation, initial expression was set to the root-population mean in the control and to five standard deviations below that mean, truncated at zero, in the perturbed cells, with the remaining genes unchanged, and 30 stochastic trajectories per root cell and condition were used to estimate the change in mean fate probabilities under matched random seeds. For each scenario, agents received the log-transformed expression profiles, observation times and cell-type labels at the training snapshots, together with the prediction times and the common terminal-state classifier.

The six scores measure velocity-field accuracy (M1), growth-driver recovery (M2), holdout distribution matching (M3), per-cell fate prediction (M4), perturbation-response recovery (M5) and gene-regulatory-network recovery (M6), with fate prediction evaluated for the initial cells (root cells in the simulator). The overall score is the unweighted mean of M1–M6, with missing or invalid task outputs scored as zero.

**Definition of the six scores.** Table 7 specifies how each prediction is compared with simulator ground truth. All six scores range from zero to one, with higher values indicating better recovery. For M2 and M6, area under the precision–recall curve (AUPRC) was computed as average precision, using absolute predicted scores to rank genes or edges.

Table 7: Definitions of the DynBench100 component scores.

| Task | Score calculation |
|---|---|
| M1: velocity-field accuracy | Cells are grouped by observation-time bin and true fate. Within each group containing at least five cells, predicted and true velocities are averaged separately and their cosine similarity is calculated. Negative similarities and comparisons with a zero-norm vector receive zero. M1 is the mean across eligible groups. |
| M2: growth-driver recovery | AUPRC for ranking the simulator's growth-driver genes above the other measured genes. |
| M3: holdout distribution matching | Wasserstein-1 distance between predicted and withheld expression distributions, converted to a score relative to within-group sampling variation and between-group differences. The conversion is defined below. |
| M4: per-cell fate prediction | Fraction of initial cells for which the most probable predicted terminal fate agrees with the most probable simulator-derived fate. |
| M5: perturbation-response recovery | Equal-weight mean of response-detection F1 and sign accuracy across gene–fate pairs. A response is detected when the absolute change in fate probability is at least 0.01. F1 compares detected pairs with the ground truth. Sign accuracy is the fraction of truly responsive pairs with the correct predicted direction of change. |
| M6: regulatory-network recovery | Equal-weight mean of edge-existence AUPRC and regulatory-sign accuracy. AUPRC considers all ordered gene pairs, including self-regulation, with unreported edges assigned zero. Sign accuracy compares activation or repression among reported edges that are present in the true network. |

For M3, we compared predicted and observed expression distributions separately at each withheld time in log-transformed gene-expression coordinates. At time $t_k$, we normalized weights within each population and calculated the Euclidean Wasserstein-1 distance $d_k$ using at most 2,000 cells from each population. The sampling reference $d_{\text{noise},k}$ was the median distance between bootstrap samples within fate groups at that time, using 30 repetitions per eligible group. The mismatch reference $d_{\text{mismatch},k}$ was the median distance between different fate groups at the same time. For $K$ withheld times, the score was

$$r_k = \frac{d_k - d_{\text{noise},k}}{d_{\text{mismatch},k} - d_{\text{noise},k}}, \qquad M3 = \frac{1}{K}\sum_{k=1}^{K} \max\{0, 1 - \max(0, r_k)\}.$$

Each time point contributes equally. Scores decrease continuously from one at the sampling reference to zero at the mismatch reference and remain zero for larger distances.

**Velocity coordinates and initial perturbations.** We compared predicted velocities with ground truth in the simulator's untransformed gene-expression coordinates and time units by reversing each method's coordinate and time transformations, including the Itô correction for diffusion when the model defined a continuous stochastic differential equation and using the chain rule for deterministic velocity fields. To evaluate perturbation responses, we applied the control and perturbed initial states defined above in each method's fitting coordinates, generated predictions using its fitted model, simulation procedure and fate calculation under matched random seeds, and evaluated all six task outputs from the same numerical run.

#### S4.2 Agent comparison

We evaluated CellCompass, OpenAI Codex, Claude Code (with MiMo or DeepSeek), and Biomni [14] across all 100 scenarios under identical input and output specifications (Table 8). CellCompass achieved a mean total score of 0.712 with GPT-5.5 high and 0.667 with MiMo-v2.5-pro. To assess run-to-run variation, we also evaluated 20 scenarios three times each across the six primary agent configurations (omitting the lower-reasoning Codex setting), on which CellCompass with GPT-5.5 high achieved a mean score of 0.672. The three runs were averaged within each scenario, and the resulting 20 pairs were compared using two-sided Wilcoxon signed-rank tests, with Holm correction for the five comparisons against CellCompass with GPT-5.5 high; error bars in Fig. 2f show the standard deviation across the 60 runs.

Table 8: DynBench100 mean component scores across 100 scenarios. Each row evaluates a complete agent system. Higher is better.

| Method | M1 | M2 | M3 | M4 | M5 | M6 | Total |
|---|---|---|---|---|---|---|---|
| CellCompass (GPT-5.5 high) | 0.946 | 0.271 | 0.833 | 0.690 | 0.759 | 0.774 | 0.712 |
| CellCompass (MiMo-v2.5-pro) | 0.883 | 0.283 | 0.738 | 0.649 | 0.723 | 0.724 | 0.667 |
| Codex (GPT-5.5 high) | 0.875 | 0.280 | 0.831 | 0.450 | 0.721 | 0.635 | 0.632 |
| Biomni | 0.878 | 0.273 | 0.804 | 0.442 | 0.728 | 0.536 | 0.610 |
| Codex (GPT-5.5 medium) | 0.841 | 0.256 | 0.785 | 0.413 | 0.692 | 0.536 | 0.587 |
| Claude Code (DeepSeek) | 0.844 | 0.257 | 0.737 | 0.432 | 0.673 | 0.556 | 0.583 |
| Claude Code (MiMo) | 0.851 | 0.269 | 0.758 | 0.402 | 0.655 | 0.464 | 0.566 |

#### S4.3 Contribution of the dynamical engine

We compared the complete CellCompass system with a version that lacked the dynamical engine and its training and simulation tools but retained scientific reasoning, proposal review, the same public inputs and general-purpose scientific Python. Both versions used GPT-5.5 with high reasoning and the same per-run budget, and each was run twice on all 40 medium-complexity scenarios with paired numerical seeds (160 runs in total). Their predictions were evaluated using the same six metrics as in the agent comparison.

We averaged the two runs within each scenario and compared the resulting 40 pairs using two-sided Wilcoxon signed-rank tests, with Holm correction across the total score and the six component scores, and obtained confidence intervals by bootstrap with 20,000 resamples. The mean six-task score was 0.736 with the engine and 0.672 without it (paired difference 0.064, 95% confidence interval 0.048–0.080, adjusted $P = 2.0 \times 10^{-8}$), and CellCompass scored higher in 35 of 40 scenarios (Fig. 2g and Supplementary Fig. 2). Fate and regulatory-network recovery showed the largest gains, and velocity, distribution and perturbation-response scores were also significantly higher after multiplicity correction, whereas growth-driver recovery did not differ significantly.

### S5 Context-dependent EMT dynamics with an existing velocity–growth model

#### S5.1 Model selection and data preparation

The EMT study required a model of transcriptional state change and effective abundance across cancer-cell lines and stimuli. Velocity–growth flow matching (VGFM) already represents both quantities, so CellCompass selected this formulation and fitted it separately to the 12 conditions. Summaries of the two interaction rounds in the EMT study are given below.

**Round 1 (condition-specific dynamics, summarized).** Given the Cook and Vanderhyden EMT time-course scRNA-seq dataset (GSE147405), the researcher asked CellCompass to fit and evaluate separate dynamical models across the 12 cell-line $\times$ stimulus induction time courses using a shared preprocessing and gene-program framework, compare generated and observed trajectories, classify shared versus context-specific EMT routes, and rank early, intermediate and late EMT-associated programs. CellCompass obtained GSE147405, excluded the withdrawal samples, prepared twelve condition-specific datasets with a shared normalization, highly-variable-gene and PCA representation and a common set of gene-program scores, and judged that an existing velocity–growth model (VGFM) already represented what the experiment constrains. It fitted one VGFM model per condition, compared generated with observed states at every time point, and reported that EMT routes were strongly condition-specific: full mesenchymal conversion was clearest in DU145 under TGF-$\beta$1, partial or hybrid routes in MCF7 under TGF-$\beta$1 and OVCA420 under EGF, and several conditions retained epithelial character. It noted that growth and depletion differed sharply between conditions, which supported the choice of a model with a growth term.

**Round 2 (mechanism analysis, summarized).** The researcher then asked CellCompass to investigate the mechanism behind the strongly condition-specific EMT routes. CellCompass organized the fitted differences into three layers: stimulus-proximal signaling (TNF/NF$\kappa$B, TGF-$\beta$/SMAD and EGFR/AP-1) set the early direction of each response; the initial state of each cell line (epithelial, interferon or stress, partial-basal, proliferative) determined whether that direction led to full EMT-TF/ECM conversion, a partial or hybrid state, or an epithelial-retaining response; and the growth term separated expanding routes (A549 and DU145 under EGF, DU145 and OVCA420 under TNF) from depleted, stress-associated routes (MCF7 under TGF-$\beta$1 or TNF, A549 under TGF-$\beta$1). Stimulus identity explained most of the variation in pathway and remodeling modules, whereas epithelial-junction, interferon and cell-cycle modules carried the largest cell-line effects.

**Data preprocessing and VGFM fitting.** The experiment profiles A549, DU145, MCF7 and OVCA420 cancer cells exposed to EGF, TGF-$\beta$1 or TNF at 0 d, 8 h, 1 d, 3 d and 7 d [15]. After excluding withdrawal samples from GSE147405, the remaining 34,994 cells and 11,058 genes across the 12 cell-line $\times$ stimulus time courses were normalized to 10,000 counts per cell, $\log(1 + x)$-transformed and projected from 2,000 highly variable genes to 50 principal components, with the same preprocessing and gene-program definitions used in all conditions. Each condition was fitted independently across ordinal model times $0, 1, 2, 3$ and 4 using deterministic VGFM (Supplementary Note S3) and simulated forward from all of its earliest-time cells, with total simulated particle weight describing changes in effective abundance.

#### S5.2 Evaluation and biological interpretation

Generated and observed populations were compared at all measured times, and gene-program scores summarized epithelial identity, core EMT, partial or basal hybrid state, extracellular-matrix remodeling, TGF-$\beta$/SMAD, TNF/NF-$\kappa$B, EGFR/AP-1, interferon/JAK–STAT, stress and cell-cycle programs. Condition-level mean Wasserstein-1 distance ranged from 10.16 to 14.96, and mean total-mass variation ranged from 0.0023 to 0.0908. Observed and generated time courses and endpoint comparisons are shown in Supplementary Figs. 3 and 4.

The predicted EMT responses differed among stimuli and cell lines. Under TGF-$\beta$1, A549 and OVCA420 showed the largest increases in the core-EMT and extracellular-matrix modules while retaining baseline epithelial-junction expression (high partial/hybrid scores), DU145 combined a more moderate core-EMT shift with clearer downregulation of canonical epithelial markers (*CDH1, EPCAM* and *CLDN4*), and MCF7 retained a stronger partial or basal and stress-associated character. Accordingly, the first-round screen—which required both a mesenchymal-score increase above 0.25 and an epithelial-score decrease of more than 0.05—highlighted DU145 under TGF-$\beta$1, whereas the two-axis route map in Fig. 3c classifies endpoints by core-EMT versus hybrid-score shifts. EGF and TNF responses varied substantially between cell lines. Population change and the prevalence of each endpoint program across the 12 conditions are reported in Supplementary Fig. 5.

**Gene-program scores and variation across conditions.** We standardized each gene's log-normalized expression across the observed cells from all 12 conditions and defined each program score as the mean standardized expression of its constituent genes, using nine route-summary programs in the first analysis round (Table 9; Fig. 3c and Supplementary Figs. 3–5) and ten signaling and cell-state modules in the second mechanism-focused round (Table 10; Fig. 3e–g). To follow each program along simulated trajectories, we fitted ridge regression (penalty 10, standardized inputs) to predict program scores from the 50-dimensional RNA representation on observed cells and applied the fitted models to simulated states, weighting population means by the simulated particle weights.

For each program and condition, we calculated the change from day 0 to day 7 using the weighted simulated means and, separately, the observed means. Denoting either set of changes by $y_{ls}$ for cell line $l$ and stimulus $s$, we used an additive model to separate variation associated with cell line, stimulus and their combination. With bars denoting averages over the indicated index, the sums of squares were

$$\mathrm{SS}_{\text{cell line}} = 3\sum_{l=1}^{4}(\overline{y}_{l\cdot} - \overline{y})^2,$$

$$\mathrm{SS}_{\text{stimulus}} = 4\sum_{s=1}^{3}(\overline{y}_{\cdot s} - \overline{y})^2,$$

$$\mathrm{SS}_{\text{combination}} = \sum_{l=1}^{4}\sum_{s=1}^{3}(y_{ls} - \overline{y}_{l\cdot} - \overline{y}_{\cdot s} + \overline{y})^2.$$

We divided each sum of squares by the total to obtain its fraction of the variation. The last component measures departures from additive cell-line and stimulus effects.

Route classes in Fig. 3c and Supplementary Figs. 4 and 5 were assigned from the generated day-0-to-day-7 changes in the core EMT and hybrid programs: conditions with strong core-EMT induction ($\geq 0.30$, labeled mesenchymal conversion in Fig. 3c, which in A549 and OVCA420 co-occurred with retained epithelial-junction expression), remaining conditions with a hybrid increase of at least 0.18 (partial or hybrid EMT), and conditions with a core-EMT change below 0.08 in magnitude and a hybrid change below 0.12 (epithelial-retaining responses). All other conditions were classed as mixed or weak routes.

Table 9: Measured genes in the nine programs used to describe EMT routes.

| Program | Measured genes |
|---|---|
| Epithelial | *EPCAM, KRT8, KRT18, KRT19, KRT7, CDH1, CLDN3, CLDN4, CLDN7, OCLN, TJP1, MUC1, DSP, DSG2, JUP* |
| Hybrid | *ITGA6, ITGB4, LAMC2, LAMB3, KRT17, VIM, CDH1, EPCAM, JAG1, NOTCH1, AREG, TGFBI, SERPINE1, S100A2* |
| Mesenchymal | *VIM, FN1, SNAI1, TAGLN, COL1A1, ITGA5, MMP9* |
| Core EMT | *VIM, FN1, SNAI1, TGFBI, SERPINE1, MMP9, ITGA5, COL1A1* |
| Immediate response | *FOS, JUN, JUNB, JUND, FOSB, EGR1, ATF3, IER2, DUSP1, DUSP5, DUSP6, NR4A1, NR4A2, KLF4, KLF6, BTG2* |
| Inflammatory TNF response | *NFKBIA, NFKBIZ, TNFAIP3, CXCL8, CXCL1, CXCL2, CXCL3, ICAM1, BIRC3, RELA, JUNB, CCL2* |
| Partial EMT | *TGFBI, SERPINE1, LAMC2, LAMB3, ITGA6, ITGB4, JAG1, NOTCH1, S100A2, KRT17, AREG, EREG, PLAUR, ANXA1* |
| Late ECM–mesenchymal | *COL1A1, COL5A2, FN1, THBS1, ITGA5, TAGLN, MMP9, TIMP1* |
| Growth and cell cycle | *MKI67, TOP2A, PCNA, MCM2, MCM3, MCM4, MCM5, MCM6, MCM7, CDK1, CCNB1, CCNB2, CDC20, AURKA, AURKB, UBE2C, TYMS, RRM2* |

Table 10: Measured genes in the ten programs used to compare signaling and cell-state changes across conditions.

| Program | Measured genes |
|---|---|
| EGFR/AP-1/MAPK | *EGFR, ERBB2, ERBB3, AREG, EREG, HBEGF, FOS, FOSB, JUN, JUNB, JUND, EGR1, DUSP1, DUSP5, DUSP6* |
| TNF/NF-$\kappa$B | *NFKBIA, NFKBIZ, NFKB1, NFKB2, RELA, RELB, TNFAIP3, BIRC3, ICAM1, CXCL1, CXCL2, CXCL3, CXCL8, CCL2* |
| TGF-$\beta$/SMAD targets | *TGFB1, TGFB2, TGFBR1, TGFBR2, SMAD2, SMAD3, SMAD4, SMAD7, TGFBI, SERPINE1, CTGF, PMEPA1, ID1, ID2, JUNB* |
| EMT transcription factors | *SNAI1, SOX4, HMGA2* |
| ECM remodeling | *COL1A1, COL5A2, FN1, THBS1, ITGA5, MMP9, TIMP1, TAGLN* |
| Epithelial junctions | *EPCAM, CDH1, KRT8, KRT18, KRT19, KRT7, CLDN3, CLDN4, CLDN7, OCLN, TJP1, DSP, DSG2, JUP* |
| Partial/basal hybrid | *LAMC2, LAMB3, ITGA6, ITGB4, KRT17, S100A2, JAG1, NOTCH1, TGFBI, SERPINE1, PLAUR, ANXA1* |
| Cell cycle and growth | *MKI67, TOP2A, PCNA, MCM2, MCM3, MCM4, MCM5, MCM6, MCM7, CDK1, CCNB1, CCNB2, CDC20, AURKA, AURKB, UBE2C, TYMS, RRM2* |
| Interferon/JAK–STAT | *STAT1, STAT2, IRF1, IRF7, ISG15, IFIT1, IFI6, IFI27, OAS1* |
| Stress/p53/apoptosis | *TP53, CDKN1A, MDM2, GADD45A, DDIT3, ATF3, BAX, BBC3, PMAIP1, BTG2, IER3, HMOX1* |

### S6 PulseBridge for metabolic-RNA-guided drug-response dynamics

#### S6.1 Incorporating metabolic RNA labeling into the dynamics

Metabolic RNA labeling provides information about the direction of molecular change within a sampling interval, whereas the existing formulations used in this comparison were fitted to populations at its endpoints. CellCompass designed PulseBridge to use this directional information to guide cell coupling, paths and training while accounting for differences between labeled and unlabeled cells at day 0. Summaries of the problem definition, the limitation of current methods and the proposal review from the CellCompass design session are given below.

**CellCompass proposal (problem formulation, summarized).** Metabolic-labeling temporal scRNA-seq provides same-cell new- and old-RNA measurements, but in the Well-TEMP-seq HCT116 5-AZA-CdR study the day-0 baseline includes both an unlabeled control group and a metabolically labeled (IAA-treated) group. Prior metabolic-direction formulations used new-to-old RNA estimates to guide local velocity directions without using the day-0 control group to separate treatment-response dynamics from labeling-associated baseline shifts. The modeling goal is to learn a continuous balanced dynamics model whose trajectories reflect metabolic-pulse-informed treatment progression rather than day-0 labeling differences.

**CellCompass proposal (limitation of current methods, summarized).** Balanced OT-CFM and prior metabolic-direction methods can fit endpoint distributions or align raw new-to-old RNA directions while retaining an unquantified component along the day-0 control-versus-labeled variation axis, so apparent early 5-AZA-CdR progression can partly reflect day-0 labeling differences.

**Proposal reviewer (summarized).** The reviewer judged the idea ready for algorithm development and asked that the next proposal estimate the labeling-associated subspace from training cells only, keep held-out and future data out of fitting, retain a balanced model, and limit causal claims about treatment to what the pulse-control identifiability evidence supports.

**Estimating metabolic directions.** PulseBridge was developed using Well-TEMP-seq measurements of HCT116 cells exposed to 5-aza-2′-deoxycytidine (5-AZA-CdR, decitabine), available under GSE194357 [16]. The analysis included 20,000 cells, with 4,000 cells from each of five samples (unlabeled control, labeled day 0 and treatment days 1, 2 and 3), whose RNA states were represented in 30 dimensions. Among the 16,000 labeled cells, metabolic directions from 12,816 cells were used for training and those from 3,184 cells were held out for evaluation, while the RNA states of the held-out cells remained available during fitting.

For each gene, the estimated change is $n/L - \gamma o$, where $n$ and $o$ are newly synthesized and pre-existing RNA counts, $L$ is the labeling duration and $\gamma$ is a decay coefficient; the first term estimates synthesis and the second accounts for loss of pre-existing RNA. The decay coefficient is estimated from the ratio of summed new to old RNA in labeled day-0 training cells, divided by $L$. Projecting these changes through the RNA loading matrix gives a direction $g$ in the same coordinates as the RNA state, and its confidence increases with the square root of the cell's total new-RNA count.

Because labeling altered the day-0 cell-state distribution, CellCompass designed PulseBridge to remove components of the measured directions aligned with labeling-associated variation. To estimate this variation, each labeled day-0 training cell was matched to its nearest unlabeled control in RNA space, and a confidence-weighted singular value decomposition of the differences between these cells identified three modes of variation, represented by the orthonormal columns of $N$. The projection $P = I - NN^{\mathsf{T}}$ removes these components from the metabolic direction $g$, giving $g^{\perp} = Pg$. For any direction $z$, the fraction $\ell(z) = \|N^{\mathsf{T}} z\|^2/\|z\|^2$ measures the contribution of labeling-associated variation, and its tolerance $b$ is estimated from the same day-0 differences.

**Cell coupling and conditional paths.** A coupling between observed populations determines which early and later cells are connected, but does not specify how cell state changes between them. PulseBridge uses the metabolic direction to adjust this intermediate path. For a candidate pair $x, y$ from adjacent times, the conditional state at position $\tau$ is

$$\psi_\tau = (1-\tau)x + \tau y + \tau(1-\tau)r, \qquad 0 \le \tau \le 1. \tag{S6.1}$$

The first two terms interpolate linearly between the paired RNA states, whereas the additional term changes the intermediate path and is zero at both endpoints, leaving the initial and terminal states unchanged.

Let $c_{xy}$ be the direction's confidence after accounting for support and agreement with the endpoint displacement. The full adjustment $r_0 = 0.3 c_{xy} g^{\perp}$ is capped in length at $\min(\|y-x\|/2, 5)$. PulseBridge compares $r \in \mathcal{R}_{xy} = \{0, r_0/2, r_0\}$ using the tangent $d_r = y - x + (1-2\tau_*)r$ at $\tau_* = 0.25$ and the score

$$J_{xy}(r) = c_{xy}\left[0.65\{1-\cos(Pd_r, g^{\perp})\}_+ + 0.15\{\ell(d_r) - b\}_+^2 + 0.01\|r\|^2\right],$$

where $\{z\}_+ = \max(z, 0)$, and the three terms measure directional disagreement, excess labeling-associated motion and adjustment size. Unsupported or conflicting directions receive $c_{xy} = 0$, giving a linear path.

The selected adjustment and cell-pair cost are

$$r^*_{xy} = \underset{r \in \mathcal{R}_{xy}}{\operatorname{argmin}}\ J_{xy}(r), \qquad C_{xy} = \|y-x\|^2 + J_{xy}(r^*_{xy}).$$

Balanced entropic transport uses $C_{xy}$ to assign cell-pair weights, and training samples pairs with these weights and uses $r^*_{xy}$ in equation (S6.1). Metabolic directions thus guide both cell pairing and the intermediate path, while balanced transport preserves endpoint population proportions.

**Training the velocity field and generating trajectories.** For each sampled pair, training draws $\tau$ uniformly along its path. Differentiating equation (S6.1) gives the velocity target over an interval of duration $\Delta t$,

$$u_\tau = \frac{y - x + (1-2\tau)r}{\Delta t}. \tag{S6.2}$$

The velocity network is trained to predict $u_\tau$ from the intermediate RNA state $\psi_\tau$ and its corresponding time. The objective combines this flow-matching loss with two terms that retain the measured directional information,

$$\mathcal{L}_{\text{Pulse}} = \mathcal{L}_{\text{velocity}} + \lambda_{\text{dir}}\mathcal{L}_{\text{direction}} + \lambda_{\text{base}}\mathcal{L}_{\text{baseline}}, \tag{S6.3}$$

where the velocity loss measures the difference between the predicted rate of RNA-state change and the target defined by the conditional path, the direction loss encourages agreement with the projected metabolic direction and the baseline loss penalizes the fraction of predicted motion associated with labeling when it exceeds the tolerance $b$.

The two additional terms are weighted by confidence, so weakly supported metabolic directions contribute less to fitting; their definitions and coefficients are given at the end of this note. Because these terms also shape the learned velocity field, agreement with the observed populations is evaluated from the simulated trajectories (Table 11).

Forward trajectories are integrated from the 8,000 control and labeled day-0 RNA states under the fitted velocity field $v_\theta(\psi, t)$ with uniform particle weights.

#### S6.2 Evaluation and biological interpretation

**Pulse-control identifiability error.** Pulse-control identifiability error (PCIE) measures agreement between simulated changes in RNA state and held-out metabolic directions, with an additional penalty for excess labeling-associated motion. To compute it, each cell $i$ with a held-out direction $g_i$ was matched to the nearest simulated RNA state at the same time, and the displacement of that particle to its next recorded state, divided by elapsed model time, defined $\widehat{v}_i$. Nearest-state searches used a random subset of 4,096 particles when more were available, with particle identities retained at the next time. For the evaluated held-out cells $\mathcal{H}$, with count-based confidence $c_i$, the score is

$$\mathrm{PCIE} = \frac{\sum_{i\in\mathcal{H}} c_i \left[1 - \cos(P\widehat{v}_i, Pg_i) + 0.25\{\ell(\widehat{v}_i) - b\}_+\right]}{\sum_{i\in\mathcal{H}} c_i}. \tag{S6.4}$$

The first term compares the projected directions $P\widehat{v}_i$ and $Pg_i$, where $P = I - NN^\mathsf{T}$ removes the labeling-associated directions estimated during training. The second penalizes the part of $\ell(\widehat{v}_i)$, the fraction of squared predicted motion along those directions, that exceeds the tolerance $b$. Lower PCIE therefore indicates closer agreement with the held-out directions and less excess labeling-associated motion. Scores were pooled across evaluation times using the held-out cells' count-based confidence, rather than the pair-dependent confidence used for path selection.

PCIE was evaluated using 2,400 of the 3,184 held-out directions, from 825 cells at day 0, 791 at day 1 and 784 at day 2. The 784 day-3 directions were excluded because the simulated trajectories end at day 3 and therefore have no later state from which to calculate a forward displacement. The 2,400 evaluation directions were withheld from parameter fitting but were used to compare candidate models during development.

Table 11: PulseBridge evaluation on the HCT116 Well-TEMP-seq panel. Lower values are better.

| Method | Mean Wasserstein-1 distance | PCIE |
|---|---|---|
| PulseBridge | 0.3011 | 0.6804 |
| WFR-FM | 0.3079 | 0.9882 |
| CRUFM | 0.3785 | 1.0108 |
| VGFM | 0.3878 | 1.0089 |
| SF2M | 0.4729 | 0.9975 |
| Dynamical OT | 0.5316 | 1.0159 |
| Unbalanced OT | 0.5829 | 1.0067 |
| RUOT | 0.5881 | 1.0210 |
| Cyto simulation | 0.6156 | 1.0143 |

PulseBridge reduced PCIE by 31.1% relative to WFR-FM and achieved the lowest mean Wasserstein-1 distance, slightly below that of WFR-FM (Table 11). Supplementary Fig. 6 adds comparisons with observed response states and, along individual gene axes, with held-out metabolic directions and random directions of the same norm.

Along the simulated trajectories, DKK1/GAS5 reactivation appeared early and was followed by B2M/HLA/IFITM3 immune-surface remodeling and SAT1/LGALS1/MT adaptive-metabolic activation, whereas EREG/PCNA repression was most prominent by day 3. We then perturbed these gene programs in the initial RNA states, simulated the modified cells through the same fitted model and compared each perturbation with random shifts of the same norm applied to the same starting cells (Supplementary Fig. 7).

**Gene programs used in the perturbation analysis.** Each gene's log-normalized expression was standardized across the 20,000 observed cells, and expression at the initial and final simulated RNA states was estimated by averaging the standardized expression of their 30 nearest observed cells at day 0 and day 3, respectively. Using the gene sets in Table 12, reactivation, immune-surface and adaptive-metabolic scores were the means of their listed genes, and proliferation repression was minus the mean of *EREG* and *PCNA*. The anti-tumor-like score combined reactivation and immune-surface genes with proliferation repression as

$$S_{\text{anti-tumor}} = \tfrac{1}{2}\left(\frac{1}{7}\sum_{g\in G_+} z_g - \frac{1}{2}\sum_{g\in G_-} z_g\right),$$

where $G_+$ contains the seven positive genes and $G_-$ the two negative genes in the table, and the balance between anti-tumor-like and adaptive responses was defined by their score difference.

Table 12: Genes and signs used in PulseBridge perturbation readouts.

| Program | Contributing genes |
|---|---|
| Anti-tumor-like response | Positive: *DKK1, GAS5, B2M, HLA-A, HLA-B, HLA-C, IFITM3*. Negative: *EREG, PCNA* |
| Reactivation | Positive: *DKK1, GAS5* |
| Immune-surface response | Positive: *B2M, HLA-A, HLA-B, HLA-C, IFITM3* |
| Proliferation repression | Negative: *EREG, PCNA* |
| Adaptive-metabolic response | Positive: *SAT1, LGALS1, MT-CO1, MT-CO2, MT-CO3, MT-ND4, MT-CYB* |

#### S6.3 Training losses and numerical settings

For a batch of $M$ sampled paths and positions, let $v_n = v_\theta(\psi_{\tau_n}, t_n)$ and let $u_n$ be the corresponding target from equation (S6.2). The source cell supplies the projected metabolic direction $g_n^\perp$. Its confidence $c_n$ includes the adjustments for directional support and agreement with the endpoint displacement described above. With $\mathcal{A}$ denoting samples with positive confidence and a nonzero projected direction, the loss terms are

$$\begin{aligned}
\mathcal{L}_{\text{velocity}} &= \frac{1}{M}\sum_{n=1}^{M}\frac{\|v_n - u_n\|^2}{d},\\
\mathcal{L}_{\text{direction}} &= \frac{\sum_{n\in\mathcal{A}} c_n\{1-\cos(Pv_n, g_n^\perp)\}_+}{\sum_{n\in\mathcal{A}} c_n},\\
\mathcal{L}_{\text{baseline}} &= \frac{\sum_{n\in\mathcal{A}} c_n\{\ell(v_n)-b\}_+}{\sum_{n\in\mathcal{A}} c_n}.
\end{aligned}\tag{S6.5}$$

Here $d = 30$ is the RNA dimension, and velocity error is averaged over all sampled paths and RNA coordinates, whereas $\mathcal{L}_{\text{direction}}$ and $\mathcal{L}_{\text{baseline}}$ are confidence-weighted means over $\mathcal{A}$. Unlike path selection, which penalizes the square of excess labeling-associated motion, the baseline loss is linear in it.

**Numerical settings.** The loss weights were $\lambda_{\text{dir}} = 0.02$ and $\lambda_{\text{base}} = 0.01$, and balanced entropic transport used blocks of 1,000 cells with entropic regularization 0.05. The labeling duration was $L = 2/24$ days, and genes with at least ten total counts determined the 10th and 90th percentile bounds for clipping decay coefficients.

Confidence was the square root of the new-RNA count divided by the square root of its positive-count 90th percentile, capped at one. The day-0 baseline subspace was estimated by uncentered singular value decomposition of nearest-neighbor displacements weighted by the square root of confidence, and the tolerance $b$ was the confidence-weighted 90th percentile of baseline fractions, restricted to $[0.02, 0.50]$.

Directions with initial confidence below 0.05 were excluded, and each time point required at least 100 usable directions. Confidence was then multiplied by $(a - 0.1)/0.9$, restricted to $[0, 1]$, where $a$ is the median ratio of projected to original direction length. Finally, confidence was set to zero for any candidate pair whose projected direction had cosine agreement below $-0.25$ with the endpoint displacement.

### S7 LineageBridge for clone-informed fate dynamics

#### S7.1 Using clonal relationships to model fate divergence

Clonal lineage information connects early cells to their descendants and can distinguish future outcomes among transcriptionally similar progenitors. CellCompass designed LineageBridge to use this information during fitting while predicting future fate from early RNA states. During development, it compared a model without clone information, a formulation that penalized errors in terminal-fate predictions and a formulation that used clone composition to allocate transported mass among fate branches.

A summary of the interaction in the LineageBridge study is given below.

**Algorithm development and evaluation (summarized).** Given the LARRY mouse hematopoiesis scRNA-seq dataset (days 2, 4 and 6) with clone barcodes, cell annotations and SPRING coordinates, the researcher asked CellCompass to design a dynamical model that incorporates sparse and incomplete clonal lineage information, explicitly determines whether population growth and death are needed rather than assuming balanced transport, and evaluates whether inferred day-2 fate probabilities agree better with clone-derived fate outcomes than a comparable no-clone baseline. CellCompass reported a completed branch-mass formulation, internally designated Clone-CUVGF-BMF and later named LineageBridge, together with its final evaluation on the day-2/4/6 data. It defined clone-fate correlation (CFCorr) as the primary measure, selected an unbalanced velocity–growth formulation to account for expanding cell numbers across days 2, 4 and 6, summarized the quantitative comparison against the no-clone VGFM and dynamical OT baselines (Table 13), and produced the requested SPRING-coordinate maps of clone-derived and predicted fate probabilities.

#### S7.2 Development of the branch allocation

During LineageBridge development, the proposal and implementation reviews shaped how clone information entered the model. The proposal reviewer noted that day-4 cells connect the day-2-to-day-4 and day-4-to-day-6 intervals, so each cell needs a consistent allocation to fate branches in both intervals:

**Proposal reviewer.** The branch-resolved day-4 measure produced by the first interval must also be the branch-resolved source measure expected by the day4→day6 interval.

CellCompass revised the proposal to allocate branch mass jointly across the two intervals, and the implementation then went through four rounds of code review:

**Implementation reviewer (round 1, summarized).** Although the code followed the approved training schedule (3,000 epochs), it split branches independently within each time interval and averaged branch endpoints into a single barycentric state during population evaluation, rather than maintaining the joint branch-boundary coupling and branch-resolved trajectories specified in the proposal.

The subsequent code revision replaced the separate branch allocations with a joint adjustment and retained the branch particles during simulation:

**Implementation reviewer (round 2, summarized).** The updated implementation retained branch-specific heads, branch-resolved forward simulation and chunked sparse transport, but the joint branch projection did not weight clone priors by their empirical reliability: it computed clone-fate reliabilities without applying them during projection, treating raw sparse clone proportions as hard row targets rather than reliability-weighted priors.

CellCompass then changed the branch preferences to combine clone-derived fate proportions with population frequencies according to clone reliability and removed the repeated rescaling to raw clone-derived source proportions:

**Implementation reviewer (round 3, summarized).** After sparse support pruning of the transport plan, the joint branch projection no longer preserved the assigned target-cell column masses at days 4 and 6 required by the proposal.

Finally, CellCompass restored the target mass totals and added missing connections so that each target cell received its assigned mass:

**Implementation reviewer (round 4, summarized).** The revised implementation matched the approved specification, using a sparse approximation to the joint branch allocation and 3,000-epoch training from scratch, and was approved for quantitative testing.

After approval, the implementation reached a CFCorr of 0.310 in Stage 2 testing, exceeding the strongest reference method, dynamical OT (0.277), by more than 10%. Subsequent Stage 3 tuning of the branch dynamics produced the final model in Table 13 (CFCorr 0.427) while keeping Brier error, Wasserstein-1 distance and total-mass variation low.

#### S7.3 LineageBridge formulation and branch-resolved prediction

**Principle of lineage-informed dynamics.** LineageBridge models both changes in effective abundance and the distribution of future fates. The unbalanced transport calculation determines the total mass contributed by each source cell to the later population. Clone composition then guides how this mass is divided among branches, without changing the source cell's total contribution. Each branch represents a possible future lineage and has its own state-dependent dynamics. The model learns how RNA states and particle weights evolve within these branches, together with branch probabilities predicted from RNA and time.

**Using clone information to connect populations.** LineageBridge was developed using the LARRY mouse hematopoiesis dataset [17], comprising 4,638 cells at day 2, 14,985 at day 4 and 29,679 at day 6. Lineage-barcoded cells were represented by 50 principal components of a 2,447-gene expression matrix, which were used to estimate cell couplings, fit the dynamics and assign terminal fates by nearest-neighbor comparison; SPRING coordinates were used for visualization.

Each clone's observed day-6 descendants define its terminal fate composition, which is smoothed towards the day-6 population frequencies and weighted by the number of recovered descendants to reduce the influence of sparsely sampled clones. Earlier cells receive the resulting fate proportions for their clone, cells without observed terminal descendants receive the population frequencies, and day-6 cells receive fate proportions based on their annotations. These distributions then guide branch allocation during fitting.

Fitting and CFCorr evaluation use all 11 annotated day-6 classes, whereas the main fate map shows seven primary outcomes: neutrophil, monocyte, basophil, mast, megakaryocyte, erythroid and undifferentiated (the remaining four minor classes are eosinophil, lymphoid, *Ccr7*-positive dendritic and plasmacytoid dendritic cells). Program summaries combine basophil with mast and megakaryocyte with erythroid, and the lineage-program analysis focuses on the four differentiated groups.

Cell coupling is estimated separately between days 2 and 4 and between days 4 and 6. The cost $C_{ij}$ of connecting source cell $i$ to target cell $j$ combines their scaled squared RNA distance with clone information. When barcode coverage is sufficient, shared clonal ancestry lowers the cost and different clone identities increase it. The transport matrix $A$ for each interval is obtained by solving

$$\min_{A\geq 0} \sum_{ij} C_{ij} A_{ij} + \varepsilon \mathcal{H}(A) + \lambda_m \, \mathrm{KL}(A\mathbf{1} \,\|\, \mathbf{1}), \qquad A^{\mathsf{T}}\mathbf{1} = \mathbf{1}. \tag{S7.1}$$

Here $A_{ij}$ is the mass assigned to a connection, $\mathbf{1}$ denotes a vector of ones and each cell has unit reference mass. The constraint fixes the mass arriving at every target cell, while the KL penalty allows each source total $m_i = \sum_j A_{ij}$ to differ from one. Thus $m_i$ determines how strongly source cell $i$ contributes to the next population and supplies its effective-growth target.

In this objective, the transport term favors low-cost connections, the entropy term $\mathcal{H}(A) = \sum_{ij} A_{ij}(\log A_{ij} - 1)$ regularizes their allocation, and $\lambda_m$ controls the penalty for changing source mass, with $\mathrm{KL}(r\|\mathbf{1}) = \sum_i (r_i \log r_i - r_i + 1)$ for unit reference mass. The problem is solved by generalized Sinkhorn iterations [18], and branch allocation is performed afterwards using this transport solution and the clone-derived fate distributions.

Let $p_{if}$ and $p_{jf}$ denote the fate distributions assigned to source cell $i$ and target cell $j$. The initial weight of their connection on branch $f$ is

$$\Gamma^{\mathrm{init}}_{ijf} = A_{ij}\, p_{if}^{\beta_{\mathrm{s}}} p_{jf}^{\beta_{\mathrm{t}}}. \tag{S7.2}$$

The exponents $\beta_{\mathrm{s}}$ and $\beta_{\mathrm{t}}$ control the contributions of source and target fate information. These weights are then adjusted to enforce consistent day-4 fate proportions and retain the cell-level mass totals from transport, although individual cell-to-cell weights may change.

**Consistent fate allocation at day 4.** LineageBridge assigns each day-4 cell the same fate proportions in both intervals, while allowing its incoming and outgoing mass to differ. Write the branch weights as $\Gamma^{(2,4)}_{ijf}$ and $\Gamma^{(4,6)}_{jkf}$, where $i$, $j$ and $k$ index cells at days 2, 4 and 6. The incoming and outgoing mass on branch $f$ for day-4 cell $j$ are

$$I_{jf} = \sum_i \Gamma^{(2,4)}_{ijf}, \qquad O_{jf} = \sum_k \Gamma^{(4,6)}_{jkf}.$$

Let $M_{\mathrm{in},j}$ and $M_{\mathrm{out},j}$ be its totals from the initial transport solutions. Dividing by these totals separates fate composition from abundance, and averaging and normalizing the two resulting vectors over all branches $h$ gives the common fate proportions $s_{jf}$:

$$\widetilde{s}_{jf} = \frac{1}{2}\left(\frac{I_{jf}}{M_{\mathrm{in},j}} + \frac{O_{jf}}{M_{\mathrm{out},j}}\right), \qquad s_{jf} = \frac{\widetilde{s}_{jf}}{\sum_h \widetilde{s}_{jh}},$$

Multiplying these common proportions by the original transport totals gives the desired branch masses,

$$B_{\mathrm{in},jf} = M_{\mathrm{in},j} s_{jf}, \qquad B_{\mathrm{out},jf} = M_{\mathrm{out},j} s_{jf}. \tag{S7.3}$$

Connections are then rescaled by the ratio of desired to current branch mass:

$$\Gamma_{ijf}^{(2,4)} \leftarrow \Gamma_{ijf}^{(2,4)} \frac{B_{\text{in},jf}}{I_{jf}}, \qquad \Gamma_{jkf}^{(4,6)} \leftarrow \Gamma_{jkf}^{(4,6)} \frac{B_{\text{out},jf}}{O_{jf}}.$$

This update matches fate proportions without equating total masses and alternates with rescaling to the original outgoing day-2, incoming day-4 and incoming day-6 totals; when only one interval has nonzero transport mass at a day-4 cell, its proportions are used. The adjusted weights then determine how cell pairs and branches are sampled during training.

**Learning dynamics and branch probabilities.** Training samples a cell pair and a branch according to their adjusted weights. A position $\tau$ is sampled uniformly between the endpoints, giving the RNA state $z_\tau = (1-\tau)x + \tau y$ and its corresponding time. For an interval of length $\Delta t$ and a source with original transported mass $m$, the targets are

$$u = \frac{y - x}{\Delta t}, \qquad a = \frac{\log m}{\Delta t}. \tag{S7.4}$$

The velocity network learns the rate of RNA-state change $u$, while the growth network learns the rate $a$ at which log particle weight changes. Both networks receive the intermediate RNA state, time and branch identity, and their squared prediction errors are weighted by $m^{\tau-1}$ to account for the mass represented along the sampled path, as in VGFM (Supplementary Note S3, equations (S3.13)–(S3.16)).

To predict fate without clone information, a separate network $q_\psi(f \mid x, t)$ learns branch probabilities from RNA state and time. Its training targets are obtained by summing the adjusted connection weights, which combine clone-derived proportions with the transport constraints, for each branch and normalizing across branches. For day-2 cell $i$ and day-4 cell $j$, respectively,

$$\widehat{q}_{if}^{(2)} = \frac{\sum_j \Gamma_{ijf}^{(2,4)}}{\sum_h \sum_j \Gamma_{ijh}^{(2,4)}}, \qquad \widehat{q}_{jf}^{(4)} = \frac{I_{jf}}{\sum_h I_{jh}}.$$

The complete objective combines learning of velocity, growth and branch probabilities,

$$\mathcal{L}_{\text{Lineage}} = \mathcal{L}_{\text{velocity}} + \mathcal{L}_{\text{growth}} + \lambda_{\text{branch}} \mathcal{L}_{\text{branch}}, \tag{S7.5}$$

where the velocity and growth terms are mass-weighted squared errors and the branch term is the squared Hellinger distance between fate proportions predicted from source RNA and those assigned using clone information; the complete expressions are given at the end of this note.

**Prediction from early RNA.** Simulation begins by using $q_\psi$ to predict branch probabilities from an early cell's RNA state and time. The cell is represented by 11 particles, one for each branch, with initial weights proportional to these probabilities. All particles start from the same RNA state, but their states and weights subsequently evolve according to the corresponding branch-specific fields,

$$\frac{dX_f}{dt} = v_\theta(X_f, t, f), \qquad \frac{d \log W_f}{dt} = g_\phi(X_f, t, f). \tag{S7.6}$$

Here $X_f$ is the RNA state on branch $f$ and $W_f$ is its contribution to the predicted population. State and weight evolve together, while branch identity remains fixed.

For starting cell $i$, the initial weight on branch $f$ is $W_{if}(T_0) = q_\psi(f \mid x_i, T_0)/n_0$, where $n_0$ is the number of starting cells. After simulation to the final time $T$, let $K_a(X_{if}(T))$ be the probability of terminal fate $a$ assigned to that branch endpoint by inverse-distance weighting of the annotations of its 30 nearest observed day-6 RNA states. The reported fate probability is

$$\widehat{\pi}_{ia} = \frac{\sum_f W_{if}(T)\, K_a(X_{if}(T))}{\sum_f W_{if}(T)}.$$

The final prediction therefore depends on the states reached by the branches and their relative effective growth.

#### S7.4 Evaluation of clone-informed fate prediction

Clone-fate correlation (CFCorr) measures agreement between predicted fate probabilities and the observed terminal-fate proportions of each cell's clone, and the reliability-weighted multiclass Brier error measures the squared discrepancy between them. Both were evaluated on 1,728 eligible day-2 cells from 1,243 clones, and Wasserstein-1 distance (evaluated on a fixed 5,950-particle representative subset; Table 18) and total-mass variation were computed from the same generated trajectory. Because clone-derived fate proportions guided both model fitting and the CFCorr calculation, CFCorr assesses recovery of the clonal relationships in this dataset.

For day-2 cell $i$, let $p_{if}$ be its clone-derived terminal-fate proportion and $\widehat{\pi}_{if}$ the model prediction. The evaluation weight $r_i$ is the clone reliability divided by the number of eligible day-2 cells from that clone, so that large early clones do not dominate the comparison. For fate $f$, define weighted means $\bar{p}_f = \sum_i r_i p_{if} / \sum_i r_i$ and $\bar{\pi}_f = \sum_i r_i \widehat{\pi}_{if} / \sum_i r_i$. The fate-specific correlation and Brier error are

$$C_f = \frac{\sum_i r_i (p_{if} - \bar{p}_f)(\widehat{\pi}_{if} - \bar{\pi}_f)}{\sqrt{\sum_i r_i (p_{if} - \bar{p}_f)^2 \sum_i r_i (\widehat{\pi}_{if} - \bar{\pi}_f)^2}}, \qquad B = \frac{\sum_i r_i \sum_{f=1}^{11} (\widehat{\pi}_{if} - p_{if})^2}{\sum_i r_i}.$$

CFCorr is the mean of $C_f$ over all 11 terminal fates.

Table 13: LARRY evaluation at days 2, 4 and 6. Higher is better for CFCorr. Lower is better for the other columns.

| Method | CFCorr ↑ | Brier error ↓ | Mean $W_1$ ↓ | Mean TMV ↓ |
|---|---|---|---|---|
| LineageBridge | 0.4271 | 0.3542 | 6.4280 | 0.0340 |
| VGFM, no clone supervision | 0.0355 | 0.8242 | 6.7209 | 0.0137 |
| Dynamical OT | 0.2773 | 0.5999 | 6.6490 | 3.8150 |

All methods used the same day-6 reference cells to assign terminal fates and were compared against the same clone-derived fate proportions (Table 13). Supplementary Fig. 8 examines transcriptionally similar early cells with different clonal outcomes, and Supplementary Fig. 9 compares predictions for six selected lineages with those of VGFM and dynamical OT. Supplementary Figs. 10 and 11 compare the distribution of predicted fates with the outcomes of individual clones, and responses to RNA-program perturbations are shown in Supplementary Fig. 12.

#### S7.5 Early-state fate programs and model perturbations

Across the four differentiated trajectories in Fig. 5c (basophil/mast, megakaryocyte/erythroid, neutrophil and monocyte), we first scored the 1,728 eligible day-2 cells using canonical mature-marker programs (Table 14), with expression standardized separately for each measured gene; cells were divided into nine equal-sized groups after ranking by program score, and the mean barcode and

LineageBridge future probabilities were calculated in each group. To identify early expression signals for neutrophil and monocyte fate beyond mature lineage markers, we then regressed the barcode-derived fate probability on the corresponding mature-marker module score and ranked all genes other than the mature markers by their correlation with the residuals. The top eight genes defined each early-state module (Table 14), and module scores were compared with both the barcode-derived fate probabilities and the LineageBridge predictions.

For the model perturbation analysis, each RNA program in Table 14 was represented by the mean principal-component direction of its measured genes. Writing $\ell_g$ for the principal-component loading vector of measured gene $g$, a program direction was $d_G = |G|^{-1} \sum_{g \in G} \ell_g$, without normalization to one standard deviation. The eligible day-2 cells were shifted to $x_i^{\pm} = x_i \pm 5d_G$ (for the five lineage perturbation programs in Supplementary Fig. 12, the same shifts were applied to 360 day-2 cells), and the modified and unmodified cells were propagated through the same fitted LineageBridge velocity, growth and branch fields at model times 0, 1 and 2. Terminal states were converted to fate probabilities with the same 30-nearest-neighbor day-6 readout and branch-weighted aggregation used in the main evaluation. The plotted effects are the changes in predicted fate probabilities relative to the predictions from the unmodified starting cells; because $d_G$ is an unnormalized average over module genes ($5\|d_G\|_2 = 0.722$ and 0.567 for the neutrophil and monocyte mature-marker modules versus 0.542 and 0.476 for the eight-gene early-state modules) and the mature markers directly define the day-6 terminal states recognized by the nearest-neighbor readout, shifting day-2 progenitors along the mature-marker directions produces a larger direct shift in predicted terminal probability despite their weak endogenous variation in unperturbed day-2 progenitors (Fig. 5c,iii).

Table 14: Measured genes in the LineageBridge mature-marker, early-state and perturbation programs.

| Program | Measured genes |
|---|---|
| Basophil/mast mature markers | *Gata2, Ms4a2, Cpa3, Prss34, Mcpt8* |
| Megakaryocyte/erythroid mature markers | *Pf4, Ppbp, Itga2b, Car1, Car2* |
| Neutrophil mature markers | *Cebpe, Lyz2, S100a8, S100a9* |
| Monocyte mature markers | *Irf8, Csf1r, Lyz2, Ly6c2* |
| Neutrophil early-state module | *Calr, F630028O10Rik, Gsr, Sell, Emb, Golga3, Sgms2, Dstn* |
| Monocyte early-state module | *Rbms1, Sirpa, Casp6, Pou2f2, Emb, Egfl7, Cd44, B4galt5* |
| Stem/progenitor (perturbation program) | *Meis1, Mecom, Procr, Mpl, Kit* |
| Neutrophil (perturbation program) | *Cebpe, Lyz2, S100a8* |
| Monocyte (perturbation program) | *Irf8, Csf1r, Lyz2* |
| Basophil/mast (perturbation program) | *Gata2, Ms4a2, Cpa3* |
| Megakaryocyte/erythroid (perturbation program) | *Pf4, Ppbp, Itga2b, Car1, Car2* |

#### S7.6 Training losses and numerical settings

A training batch contains $M$ cell-pair and branch samples drawn according to the adjusted connection weights $\Gamma$, with a uniformly sampled position on each path. For sample $n$, denote the predicted velocity and effective growth at the intermediate state by $v_n$ and $g_n$. Their targets $u_n$

and $a_n$ are given in equation (S7.4), and their weight is $w_n = m_n^{\tau_n - 1}$. Here $m_n$ is the source cell's total mass from the initial transport solution, before branch allocation. The branch prediction $q_{nf} = q_\psi(f \mid x_n, t_n^{\text{source}})$ is evaluated at the source cell, with target $\widehat{q}_{nf}$ obtained from its normalized branch weights. The loss terms are

$$\begin{aligned}
\mathcal{L}_{\text{velocity}} &= \frac{1}{M} \sum_{n=1}^{M} w_n \frac{\|v_n - u_n\|^2}{d}, \\
\mathcal{L}_{\text{growth}} &= \frac{1}{M} \sum_{n=1}^{M} w_n (g_n - a_n)^2, \\
\mathcal{L}_{\text{branch}} &= \frac{1}{M} \sum_{n=1}^{M} \left( 1 - \sum_{f=1}^{F} \sqrt{q_{nf} \widehat{q}_{nf}} \right).
\end{aligned} \tag{S7.7}$$

Here $d$ is the RNA representation dimension and $F = 11$ is the number of branches; the velocity and growth losses average the mass-weighted errors over the batch, whereas the branch loss compares fate distributions at source cells with equal weight per sampled source.

**Numerical settings.** The velocity, growth and branch losses were assigned unit coefficients ($\lambda_{\text{branch}} = 1$). Clone fate distributions were smoothed using five pseudocounts distributed according to the day-6 population frequencies. Clone reliability was the observed terminal-descendant count divided by that count plus five, and the smoothed distribution was mixed with the population frequencies using weights equal to reliability and one minus reliability. Day-6 preferences placed 95% weight on the annotated fate and 5% on the population distribution.

RNA matching costs were divided by their block-wise 90th percentile, and the transport objective used $\varepsilon = 1$ and $\lambda_m = 50$. Clone support was the geometric mean of its counts in adjacent populations divided by that mean plus two. When both endpoint clones had support of at least 0.1, a shared barcode reduced cost by 0.75 times the smaller support, and different barcodes increased cost by 0.25 times the geometric mean of their supports; otherwise, cost used RNA distance alone. Within each block, the 160 largest positive connections per source were retained, adding connections where needed for target coverage and rescaling to preserve source totals.

Days 2, 4 and 6 used model times 0, 1 and 2 and fate-preference powers 2, 0.25 and 1.25, respectively. Each of the 28 projection iterations restored outgoing day-2, incoming day-4 and incoming day-6 totals, in that order, before updating the shared day-4 proportions. Forward Euler integration used a step size of 0.0125, and each starting cell contributed a total initial weight of $1/n_0$ across its branches.

### S8 ProteinBridge for RNA-initialized protein dynamics

#### S8.1 Learning protein dynamics from paired RNA–protein measurements

Surface-protein abundance can change asynchronously with the corresponding transcripts. Predicting future protein states from RNA therefore requires a model that connects these molecular layers over time. CellCompass designed ProteinBridge to use paired CITE-seq measurements during fitting in two ways: protein information contributes to cell coupling, and paired RNA–protein paths are used to learn how protein levels depend on RNA state and time. Once trained, the model generates RNA trajectories and predicts the corresponding protein profiles from initial RNA.

Summaries of the two interaction rounds in the ProteinBridge study are given below.

**Round 1 (algorithm development, summarized).** Given the longitudinal CD34-positive CITE-seq dataset (GSE305370, days 2, 3, 4, 7 and 10), the researcher asked CellCompass to use its multi-omic measurements to design a new algorithm. CellCompass formulated the task as predicting future surface-protein states from initial RNA states and reported that a balanced paired-coordinate flow-matching model, later named ProteinBridge, had completed development and 30 hyperparameter refinement runs, achieving the lowest surface-protein energy distance and RNA–protein cross-covariance error averaged over the four future sampling times (Table 15).

**Round 2 (downstream analysis, summarized).** The researcher then requested downstream analyses and figures comparing the model with baseline methods. CellCompass generated the quantitative comparisons against CRUFM, VGFM, balanced OT-CFM and dynamical OT (Table 15), together with time-resolved protein energy distance, cross-covariance error over time, interval-wise RNA Wasserstein-1 distance and simulated trajectories in the paired RNA–protein space.

During model development, CellCompass compared model families that differed in how RNA and protein information entered the dynamics, including unbalanced velocity–growth (CITE-VGFM), balanced (CITE-BFM) and residual stochastic-bridge (CITE-RSB-FM) flow matching, and developed the paired-coordinate formulation described below over 70 trials.

**Using paired measurements to connect populations.** ProteinBridge was developed using a longitudinal CITE-seq [19] study of human CD34-positive hematopoietic stem and progenitor cells from four healthy donors, sampled at days 2, 3, 4, 7 and 10 (GSE305370) [20], which are indexed by ordinal model times $t \in \{0, 1, 2, 3, 4\}$. The analysis included 10,000 cells with measurements of 103 antibody-derived tags (ADTs). RNA profiles were reduced to 50 components using truncated singular value decomposition and ADT profiles to 30 principal components, and the resulting components, denoted $R$ and $P$, were standardized to zero mean and unit variance.

Transcriptionally similar cells can have distinct surface-protein profiles. ProteinBridge therefore includes protein distance in the cost of coupling cells $i$ and $j$ at adjacent observation times,

$$C_{ij} = \|R_i - R_j\|^2 + \lambda_P \frac{\|P_i - P_j\|^2}{d_P}. \tag{S8.1}$$

The protein term, averaged over its $d_P$ coordinates and weighted by $\lambda_P$, raises the cost of matching cells with different protein profiles even when their RNA states are similar. Balanced entropic transport then uses the combined cost to estimate cell-pair weights while preserving the observed population proportions.

Training samples pairs according to their transport weights, and a position $\tau$ drawn uniformly between zero and one gives matched RNA and protein targets along each connection,

$$R_\tau = (1-\tau)R_i + \tau R_j, \qquad P_\tau = (1-\tau)P_i + \tau P_j,$$

with the same interpolation position used for both modalities.

**Learning RNA dynamics and the RNA-to-protein mapping.** The RNA velocity field $v_\theta(R,t)$ is fitted to $(R_j - R_i)/\Delta t$, where $\Delta t$ is the model-time interval, while the RNA-to-protein mapping $h_\eta(R,t)$ is fitted to the interpolated protein state $P_\tau$ at the corresponding RNA state $R_\tau$ and time. The mapping is also trained directly on measured RNA–protein pairs sampled from the same transport plan, with prediction errors at the source and target endpoints calculated separately and averaged. This endpoint loss receives greater weight than the loss on interpolated profiles, so the measured protein profiles remain the main constraint during training. The combined objective is

$$\mathcal{L}_{\text{Protein}} = \mathcal{L}_{\text{RNA}} + \lambda_{\text{path}}\mathcal{L}_{\text{path}} + \lambda_{\text{endpoint}}\mathcal{L}_{\text{endpoint}}, \tag{S8.2}$$

where $\mathcal{L}_{\text{RNA}}$ measures the difference between the predicted rate of RNA-state change and the rate defined by the conditional path, and the two protein losses compare the predicted profiles with the interpolated profiles along that path ($\mathcal{L}_{\text{path}}$) and with measured profiles at its endpoints ($\mathcal{L}_{\text{endpoint}}$). Both protein losses give greater weight to interior observation times; their definitions are given at the end of this note.

**Prediction from initial RNA.** Starting from a day-2 RNA state, the model simulates the RNA trajectory using the learned velocity field, and the RNA-to-protein mapping predicts a surface-protein profile at each simulated state and time,

$$\frac{dR_t}{dt} = v_\theta(R_t, t), \qquad \widehat{P}_t = h_\eta(R_t, t). \tag{S8.3}$$

The mapping predicts protein profiles at all times, including the initial time, along trajectories integrated from 2,000 day-2 RNA states with uniform particle weights (1/2,000). Unlike the joint-modality reference models, which require both initial RNA and initial ADT coordinates, Protein-Bridge initializes trajectories from day-2 RNA alone.

#### S8.2 Evaluation and biological interpretation

All five observed time points contributed to model fitting, and the four later RNA–protein populations were used to compare candidate formulations and select the model during development. Model acceptance was based primarily on the energy distance between predicted and observed surface-protein distributions in the 30-dimensional protein representation (Supplementary Note S9, equation (S9.3)), averaged over the four future sampling times. RNA Wasserstein-1 distance measured how closely the simulated RNA populations matched the observations, and relative RNA–protein cross-covariance error assessed how well the model preserved the relationship between the two molecular layers.

We measured RNA–protein coordination using cross-covariance after centering each modality. With $n$ cells, $R \in \mathbb{R}^{n\times d_R}$ and $P \in \mathbb{R}^{n\times d_P}$, the matrix is $C = (R-\bar{R})^{\mathsf{T}}(P-\bar{P})/(n-1)$, and the relative cross-covariance error is

$$\frac{\|C_{\text{pred}} - C_{\text{obs}}\|_{\text{F}}}{\|C_{\text{obs}}\|_{\text{F}}},$$

Table 15: Performance on the temporal CITE-seq benchmark. Lower values indicate closer agreement between generated and observed data.

| Method | RNA $W_1$ | Protein energy distance | Cross-covariance error |
|---|---|---|---|
| ProteinBridge | 8.236 | 0.02220 | 0.4668 |
| CRUFM | 8.452 | 0.02558 | 0.5250 |
| VGFM | 8.415 | 0.02826 | 0.5172 |
| Balanced OT-CFM | 9.693 | 0.17054 | 0.6990 |
| Dynamical OT | 7.446 | 0.73525 | 0.5373 |

evaluated using at most 1,500 paired cells per population and averaged equally across the four future sampling times.

ProteinBridge reduced protein energy distance by 13.2% and cross-covariance error by 11.1% relative to CRUFM, while dynamical OT had the lowest RNA Wasserstein-1 distance (Table 15). Supplementary Fig. 13 compares predicted terminal protein distributions, and Supplementary Fig. 14 reports errors over time and examines whether generated cells reproduce the observed protein states, phenotype proportions and variation within the population.

We used simulated trajectories to compare the timing of RNA and surface-protein programs and to examine responses to changes in initial RNA state, focusing on the 972 annotated hematopoietic stem cells (HSCs) among the 2,000 day-2 starting cells. Their unperturbed trajectories were grouped by predicted terminal phenotype (neutrophil, mast-cell, monocyte, erythroid or megakaryocyte progenitor), with cells retaining an HSC phenotype forming a separate category in population-composition comparisons. Within neighborhoods of initial RNA states, future-phenotype probabilities described the predicted outcomes, and their entropy summarized uncertainty.

To compare RNA and protein timing, we fitted linear decoders to the observed paired cells, used them to reconstruct selected RNA genes and ADT markers at each of the 41 simulated times and compared peak times in model-time coordinates, in which successive observation times are one unit apart.

**RNA and ADT programs for comparing peak times.** Table 16 lists the measured features used for the timing profiles. Only available genes and ADTs contributed, so *CD34* RNA entered the HSC program although CD34 ADT was unavailable, and CD71 served as the EryP surface marker because CD235a (glycophorin A) was not included in the antibody panel. For each predicted terminal group with at least 35 trajectories, decoded feature values were averaged at each of the 41 model times, and each feature was scaled between its minimum and maximum over time, with a constant feature profile assigned 0.5. The scaled features were then averaged within the RNA or ADT program, and after Gaussian smoothing with width 1.15 time bins, each program profile was again scaled between zero and one.

Table 16: Measured RNA genes and ADTs used in ProteinBridge timing profiles. HSC, hematopoietic stem cell; NeuP, neutrophil progenitor; MasP, mast-cell progenitor; MoP, monocyte progenitor; EryP, erythroid progenitor; MkP, megakaryocyte progenitor.

| Phenotype | RNA genes | ADTs |
|---|---|---|
| HSC | *CD34, MEIS1, MECOM, HOXA9, PROM1, SPINK2, PROCR* | CD49f, CD123, CD45RA, HLA-DR, CD62L, CD31 |

Table 16 (continued)

| Phenotype | RNA genes | ADTs |
|---|---|---|
| NeuP | *MPO, CTSG, AZU1, ELANE, PRTN3, CEBPA, SPI1, LYZ, S100A8, S100A9* | CD11b, CD11c, CD13, CD16, CD64, CD115, CD172a, CD88, HLA-DR |
| MasP | *GATA2, CPA3, TPSAB1, MS4A2, KIT, HDC, CLC, FCER1A* | Fc$\varepsilon$RI$\alpha$, CD88 |
| MoP | *SPI1, IRF8, LYZ, LST1, S100A8, S100A9, CEBPA, MS4A3* | CD11b, CD11c, CD13, CD64, CD115, CD172a, HLA-DR |
| EryP | *GATA1, KLF1, HBA1, HBA2, HBB, ALAS2, GYPB* | CD71 |
| MkP | *ITGA2B, GP9, GP1BA, VWF, NFE2, FLI1, TUBB1, GFI1B, RUNX1* | CD41, CD42b, CD62P, CD49b |

**RNA–ADT associations in observed cells.** We examined RNA–protein associations in the observed paired cells using four RNA programs (Table 17). Gene expression and ADT abundances were standardized across all 10,000 observed cells, and each RNA-program score was the mean of its available standardized genes. We then ranked cells by RNA-program score and divided them into ten equal-sized groups to plot mean ADT abundance, and also compared the upper and lower quartiles by subtracting the mean ADT abundance in cells at or below the 25th percentile from that in cells at or above the 75th percentile.

Table 17: Measured features in the observed RNA–ADT comparisons.

| Program | RNA genes | ADTs compared |
|---|---|---|
| HSC | *CD34, MEIS1, MECOM, HOXA9, PROM1, SPINK2, AVP, MLLT3, PROCR* | HLA-DR, CD62L, CD31, CD123, CD32, CD41, CD71 |
| Myeloid | *MPO, CTSG, AZU1, ELANE, PRTN3, CEBPA, SPI1, LYZ, LST1, S100A8, S100A9* | CD11c, CD64, CD45RA, CD71, CD41, CD88 |
| Erythroid | *GATA1, KLF1, HBA1, HBA2, HBB, ALAS2, GYPB* | CD41, CD32, CD71, CD62L, HLA-DR, CD13 |
| Megakaryocytic | *ITGA2B, GP9, GP1BA, VWF, NFE2, FLI1, TUBB1, GFI1B, RUNX1* | CD41, CD62P, CD32, CD49f, HLA-DR, CD45RA |

For the single-gene RNA-direction screen, a one-standard-deviation increase in each measured RNA feature was projected through the RNA loadings and divided by the standard deviation of each RNA coordinate, giving a gene-specific direction in the 50-dimensional RNA space. Each direction was applied at +5 standardized units to the same 972 HSC starting particles, which were then simulated through the fitted RNA dynamics and RNA-to-protein mapping. Terminal surface-protein coordinates were classified with a surface-protein-phenotype classifier fitted to the observed annotations, and a separate linear decoder reconstructed a megakaryocytic ADT module from CD41, CD42b, CD62P and CD49b. All effects were calculated relative to the predictions from the same unmodified HSC starting states.

The largest increases in terminal megakaryocyte-progenitor (MkP) probability were obtained for *THBS1* (+6.1 percentage points), *TUBB1* (+5.8), *GP9* (+5.7), *GP1BA* (+3.9), *VWF* (+3.4) and *ITGA2B* (+3.1). The same directions increased the decoded megakaryocytic ADT module by +0.20, +0.17, +0.17, +0.13, +0.11 and +0.10 standardized units, respectively. In the same screen, *KLF1* decreased terminal MkP probability by 1.5 percentage points and increased erythroid-progenitor (EryP) probability by 3.0 percentage points. The observed RNA–surface-protein associations used to interpret the fitted readout are examined separately in Supplementary Fig. 15.

##### S8.3 Training losses and numerical settings

For each of $M$ training samples, a cell pair is drawn from the transport plan and interpolated at a uniformly sampled position. Write its intermediate RNA and protein states as $R_n$ and $P_n$, its time as $t_n$, and its RNA velocity target as $u_n = (R_{j_n} - R_{i_n})/\Delta t_n$. A separately sampled batch of endpoint pairs supplies measured states $(R_n^a, P_n^a)$ at times $t_n^a$, where $a = 0$ denotes the source and $a = 1$ the target. With $w(t)$ denoting the protein time weight, the three loss terms are

$$\begin{aligned}
\mathcal{L}_{\text{RNA}} &= \frac{1}{M}\sum_{n=1}^{M} \frac{\|v_\theta(R_n, t_n) - u_n\|^2}{d_R}, \\
\mathcal{L}_{\text{path}} &= \frac{\sum_{n=1}^{M} w(t_n)\|h_\eta(R_n, t_n) - P_n\|^2}{d_P \sum_{n=1}^{M} w(t_n)}, \\
\mathcal{L}_{\text{endpoint}} &= \frac{1}{2}\sum_{a=0}^{1} \frac{\sum_{n=1}^{M} w(t_n^a)\|h_\eta(R_n^a, t_n^a) - P_n^a\|^2}{d_P \sum_{n=1}^{M} w(t_n^a)}.
\end{aligned} \tag{S8.4}$$

Here $d_R = 50$ and $d_P = 30$, and each squared error is averaged over the coordinates of its modality. The protein losses are also divided by the summed time weights within the batch, separately for source and target cells in $\mathcal{L}_{\text{endpoint}}$. The time weight $w(t)$ emphasizes the intermediate stages of differentiation during protein fitting.

**Numerical settings.** Using the preprocessed RNA expression matrix from GSE305370, no further library-size normalization or log transformation was applied. Truncated singular value decomposition of RNA did not subtract gene-wise means, and both this decomposition and the subsequent coordinate standardization were fitted before the 10,000-cell subset was selected. The paired cost used $\lambda_P = 0.25$, and transport used blocks of 1,000 cells and entropic regularization 0.05 after dividing each block's cost by its maximum.

The training-loss weights were $\lambda_{\text{path}} = 3$ and $\lambda_{\text{endpoint}} = 1500$, with the RNA velocity loss assigned unit weight. The protein time weight $w(t)$ was a baseline of one plus unit-height Gaussian peaks at model times 1, 2 and 3, with standard deviation 0.84 and the total capped at three.

The velocity network had hidden width 256 and five layers. The protein readout had two hidden layers of 1,024 units and SiLU activation, taking RNA, time and sine and cosine features with periods four and two model-time units as input. During simulation, RNA states were advanced in steps of 0.1 model-time units using the velocity evaluated at the current state and the midpoint time, and protein profiles were predicted at 41 times spanning model time 0 to 4.

### S9 Evaluation, statistical analysis and numerical settings

#### S9.1 Evaluation metrics and statistical analyses

**Cell-state distributions and effective abundance.** Models were evaluated using trajectories simulated from the observed initial cells to assess each study's biological predictions alongside cell-state distributions and effective abundance. Let $\widehat{\mu}_{t_k} = \sum_i \widehat{w}_i^{(k)} \delta_{\widehat{x}_i^{(k)}}$ denote a model prediction and $\mu_{t_k} = \sum_j w_j^{(k)} \delta_{x_j^{(k)}}$ the observed empirical measure.

To compare the shapes of predicted and observed cell-state distributions, we normalized their weights separately before calculating Wasserstein-1 distance,

$$\widetilde{\mu}_{\text{pred}} = \sum_i \frac{\widehat{w}_i}{\sum_\ell \widehat{w}_\ell} \delta_{\widehat{x}_i}, \qquad \widetilde{\mu}_{\text{obs}} = \frac{1}{n} \sum_{j=1}^{n} \delta_{x_j}.$$

Wasserstein-1 distance was then computed in the analysis representation, using the same estimator and preprocessing for all methods within a case:

$$W_1(\widetilde{\mu}_{\text{pred}}, \widetilde{\mu}_{\text{obs}}) = \inf_{\gamma \in \Pi(\widetilde{\mu}_{\text{pred}}, \widetilde{\mu}_{\text{obs}})} \int \|x - y\|_2 \, d\gamma(x, y). \tag{S9.1}$$

For unbalanced formulations, total-mass variation (TMV) evaluates effective abundance by comparing the unnormalized predicted change in total particle weight with the observed change in cell count relative to the initial time $t_0$:

$$\text{TMV}(t) = \left| \frac{\sum_i \widehat{w}_i(t)}{\sum_i \widehat{w}_i(t_0)} - \frac{n_t}{n_{t_0}} \right|. \tag{S9.2}$$

Energy distance measures the difference between predicted and observed distributions without requiring one-to-one cell matching [21]. Let $X$ and $X'$ be independent draws from the predicted distribution, and $Y$ and $Y'$ independent draws from the observed distribution, all expressed in the same coordinates. We estimated

$$\mathcal{E}(X, Y) = 2\mathbb{E}\|X - Y\|_2 - \mathbb{E}\|X - X'\|_2 - \mathbb{E}\|Y - Y'\|_2, \tag{S9.3}$$

using empirical pairwise averages, including diagonal pairs within each population, where the first term measures distances between predicted and observed cells and the two subtracted terms account for variation within each population.

#### S9.2 Numerical implementation

Table 18 summarizes the starting populations and simulation settings.

Table 18: Numerical settings of the reported case-study trajectories.

| Case | Starting population | Integrator and time grid |
|---|---|---|
| EMT VGFM | Every earliest-time cell in each of 12 conditions<br>counts listed below | Trajectories recorded at intervals of 0.1 model-time units<br>one deterministic simulation per condition |
| PulseBridge | 8,000 starting particles with equal weights | Trajectories recorded at intervals of 0.1 model-time units |
| LineageBridge | 4,638 $\times$ 11 = 51,018 branch particles<br>5,950 representative particles for distribution comparisons<br>all branches for fate evaluation | Forward Euler for state and log-weight<br>step size 0.0125<br>recorded at model times 0, 1, 2<br>one deterministic branch expansion |
| ProteinBridge | 2,000 RNA particles with uniform weights | 41 recorded times from 0 to 4<br>step size 0.1, with velocity evaluated at the current state and midpoint time<br>one deterministic simulation of the initial population |

The EMT initial counts for EGF, TGF-$\beta$1 and TNF were 985, 577 and 1,267 for A549; 343, 249 and 464 for DU145; 400, 523 and 319 for MCF7; and 238, 68 and 237 for OVCA420.

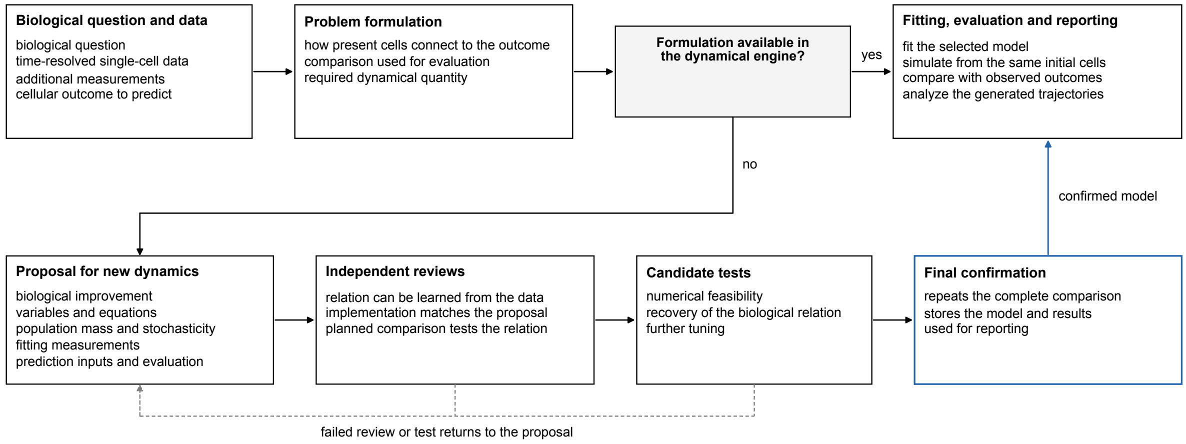

Supplementary Fig. 1: **CellCompass model selection and development of new dynamics.** Workflow from user query and experimental data to biological discovery. CellCompass selects an existing formulation from the dynamical engine when it is sufficient, or proposes and implements a task-specific extension whose proposal, code and prediction procedure undergo context-isolated review. New formulations are then tested for numerical feasibility (Stage 1), compared with reference methods on the prespecified biological measure (Stage 2) and tuned for population agreement (Stage 3), and the selected model is retained in the dynamical engine after final confirmation (Supplementary Note S1).

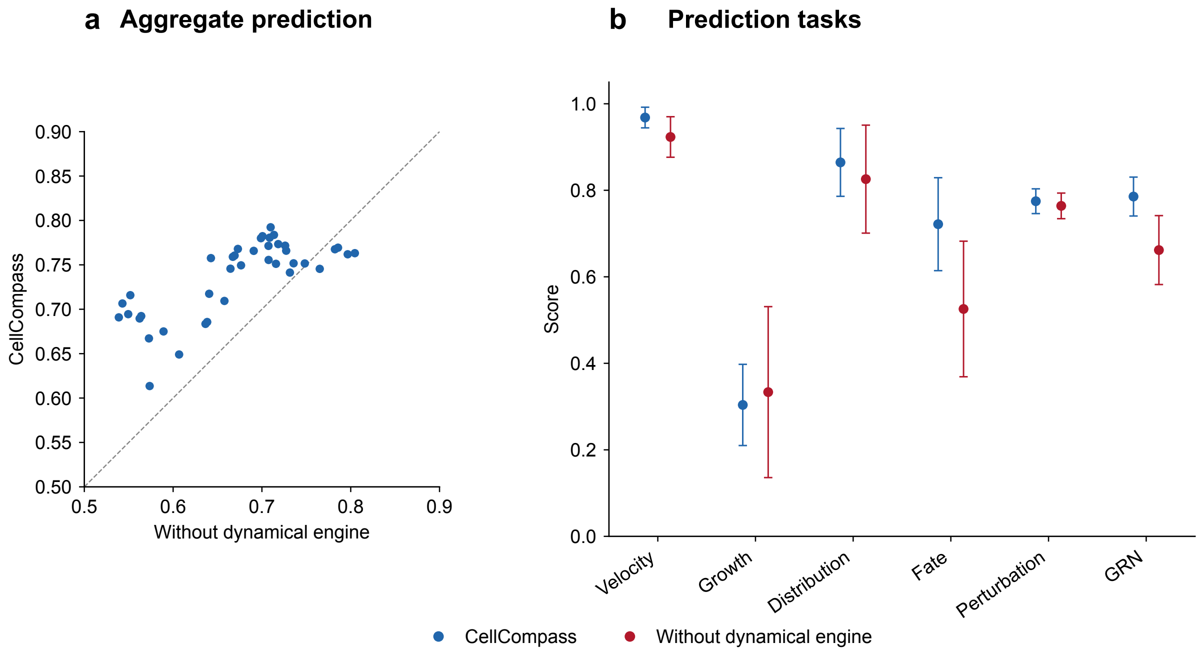

Supplementary Fig. 2: **Dynamical-engine ablation across 40 medium-complexity DynBench100 scenarios. a,** Six-task mean scores with and without the dynamical engine. Each point represents one scenario, averaged over two runs per condition. The dashed line indicates equal scores. **b,** Scores for velocity, growth-driver recovery, distribution, fate, perturbation response and gene regulatory network (GRN) recovery. Points and error bars show the mean and standard deviation across the 40 scenario means. The paired statistical comparison is reported in Supplementary Note S4.3.

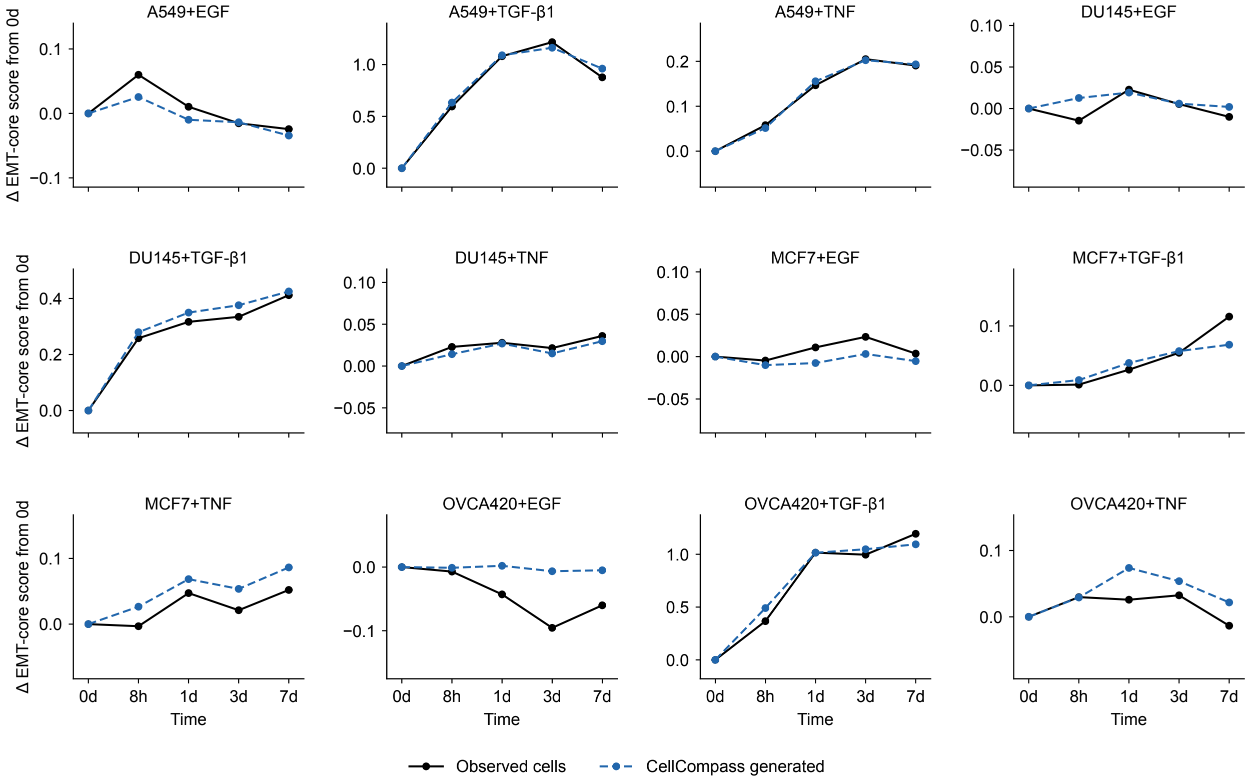

Supplementary Fig. 3: **Observed and generated EMT-core dynamics across cancer-cell contexts.** Changes in the EMT-core program are shown relative to day 0 for the 12 combinations of A549, DU145, MCF7 and OVCA420 cells with EGF, TGF-$\beta$1 or TNF. Black lines show observed cells and blue dashed lines show trajectories generated by the separately fitted VGFM model for each condition.

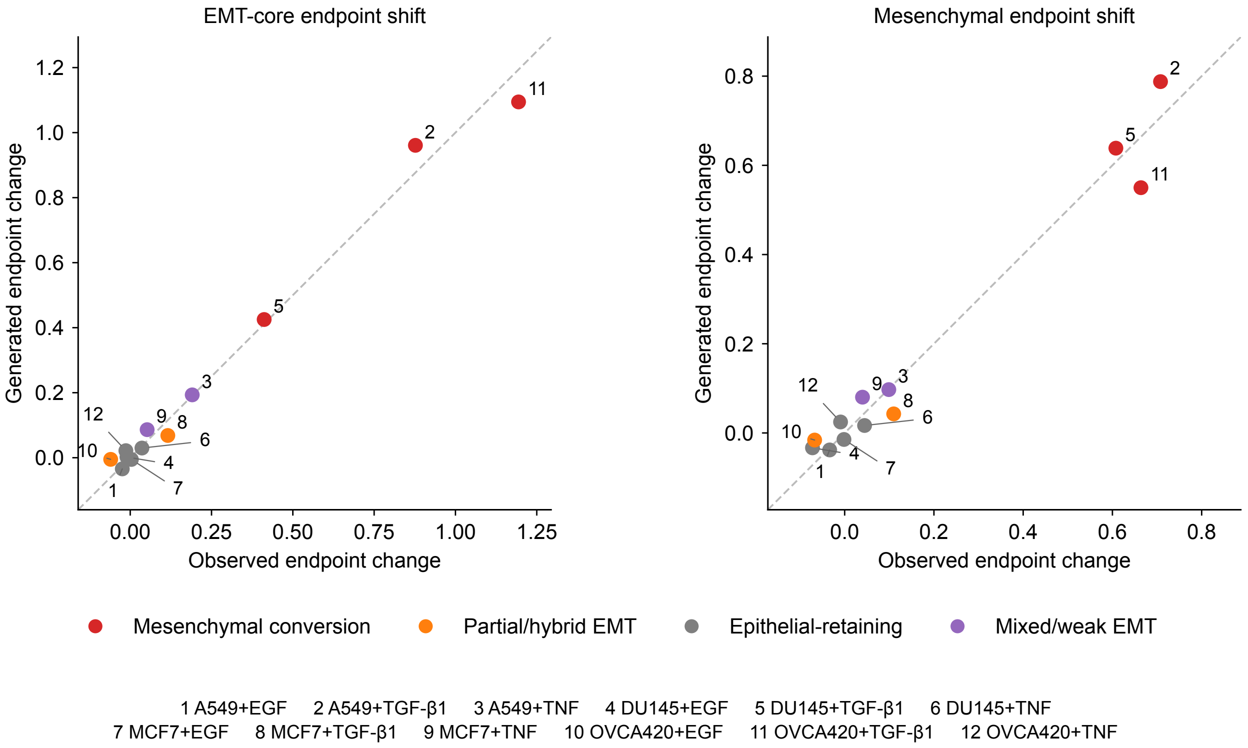

Supplementary Fig. 4: **Generated endpoints recover observed EMT program shifts.** Each numbered point represents one cell-line–stimulus combination. Predicted and observed changes from the initial to the final time point are compared for the EMT-core and mesenchymal programs. Colors indicate the EMT route classes. Pearson correlations are 0.99 for the EMT-core program and 0.98 for the mesenchymal program.

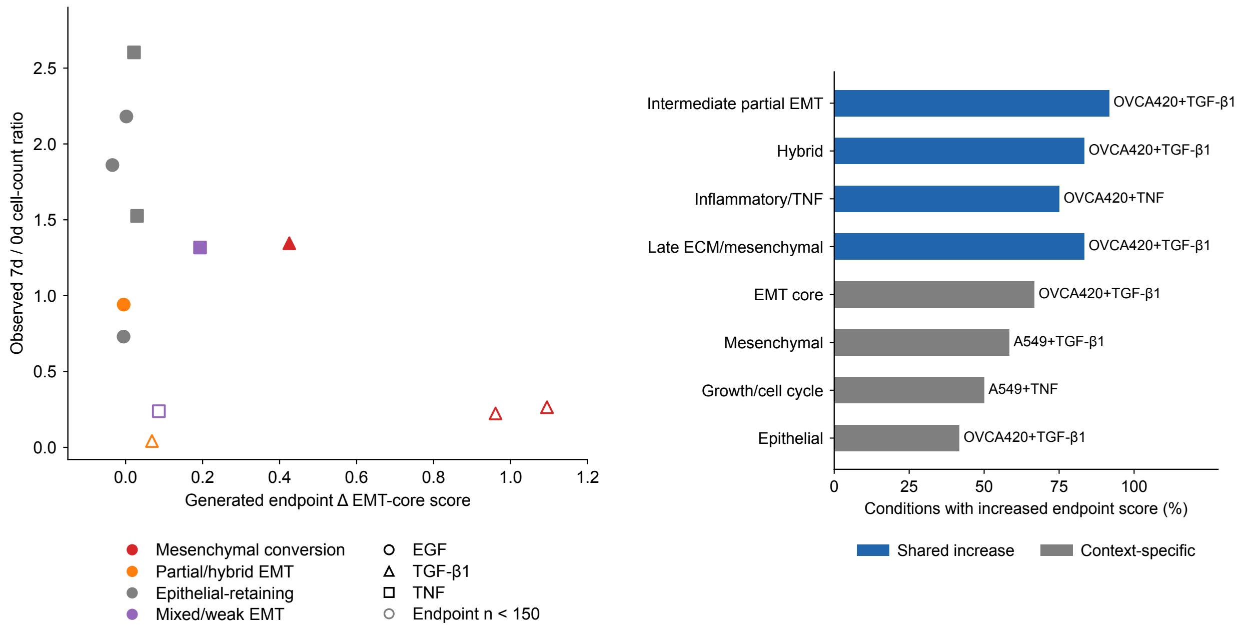

Supplementary Fig. 5: **Population change and program prevalence across EMT contexts. Left,** The generated EMT-core endpoint shift is compared with the observed day-7 to day-0 cell-count ratio for every cell-line–stimulus condition. Symbols identify the stimulus and colors identify the route class. Open symbols mark conditions with fewer than 150 endpoint cells. **Right,** The fraction of conditions with an observed endpoint increase is shown for partial EMT, hybrid, inflammatory, extracellular-matrix, core EMT, mesenchymal, growth and epithelial programs. Labels identify the condition with the strongest increase.

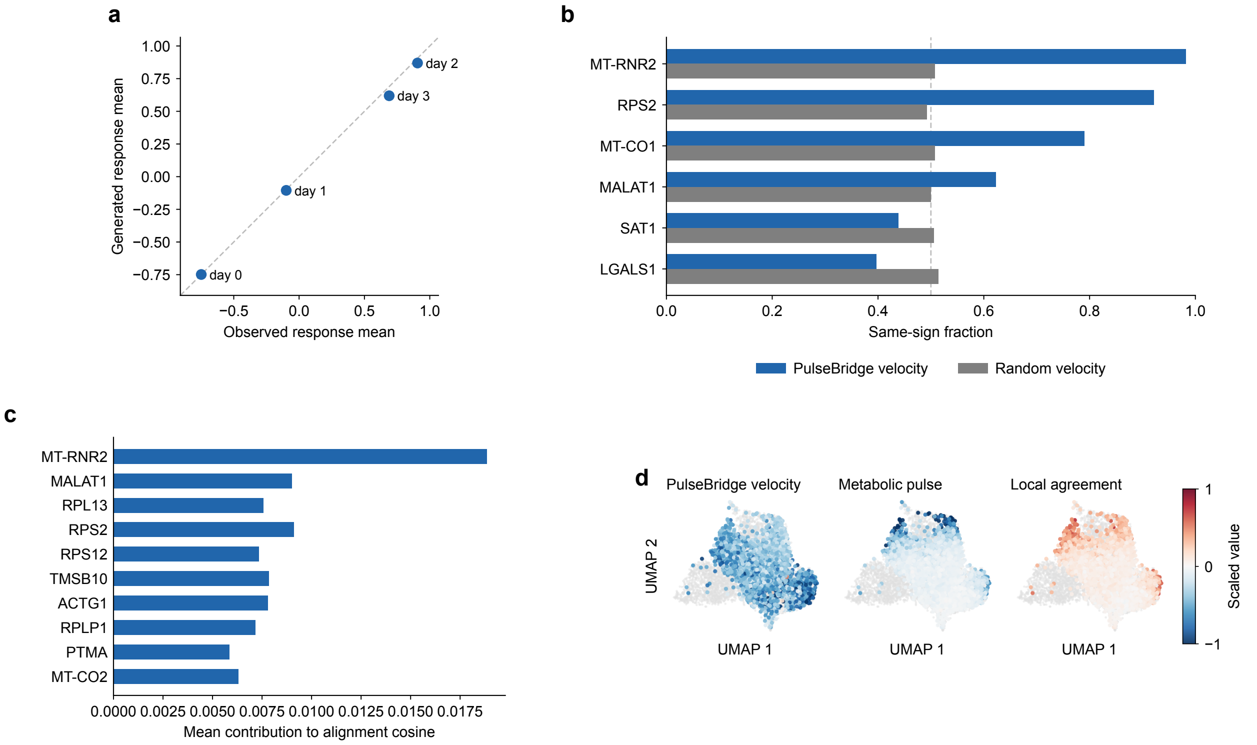

Supplementary Fig. 6: **Population and gene-level comparisons of PulseBridge dynamics.** **a,** Generated mean response states are compared with observed states across the four treatment times. **b,** For six genes, the fraction of velocity directions agreeing with the metabolic pulse is compared with same-sized random directions. **c,** Gene axes are ranked by their contribution to the pulse-alignment score. **d,** MT-RNR2 directions are shown separately for PulseBridge velocity, the metabolic pulse and their local agreement. These gene-level comparisons describe individual components of the predicted direction, whereas PCIE evaluates the joint direction in the RNA representation (Supplementary Note S6).

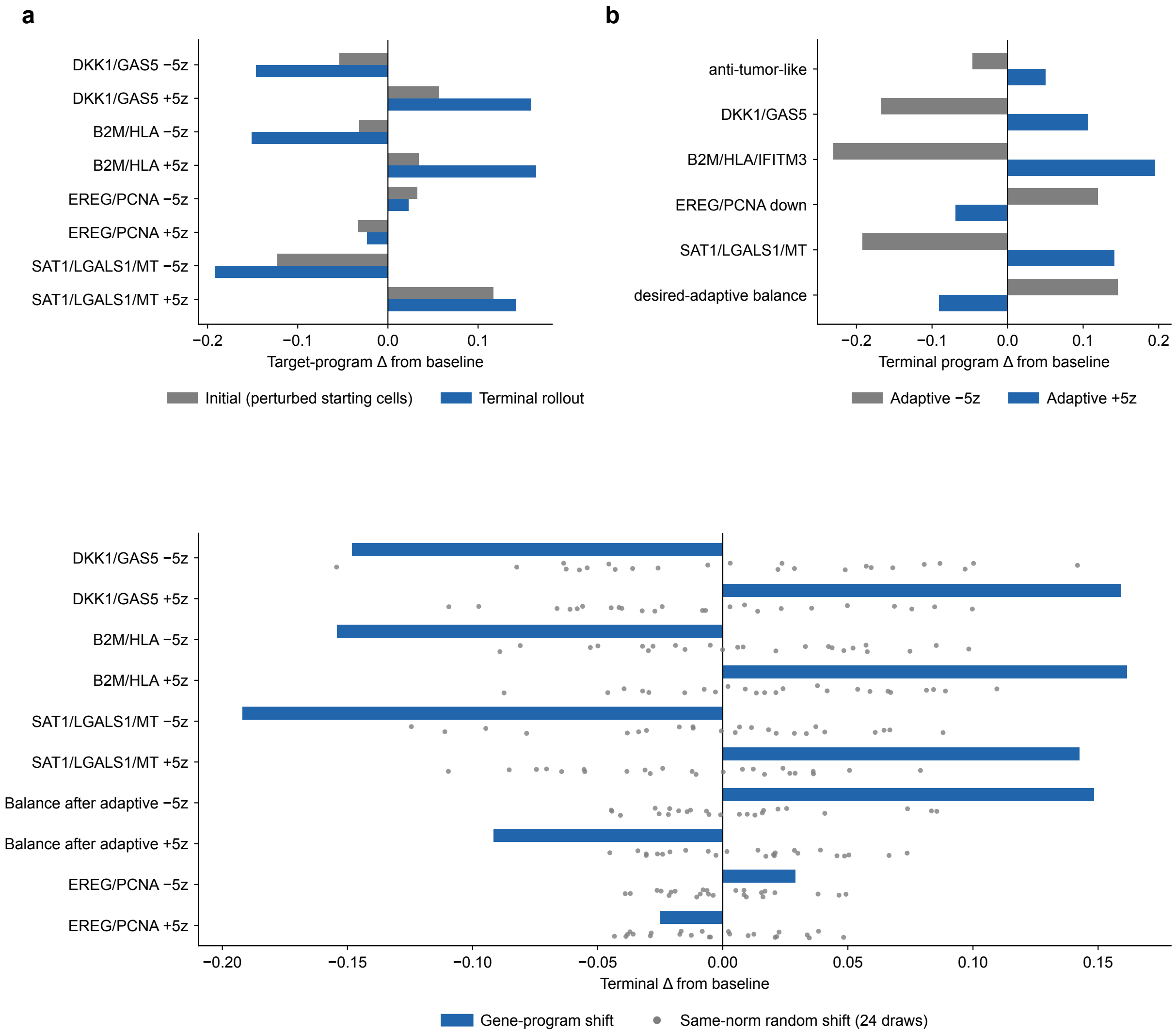

Supplementary Fig. 7: **Random-shift controls for PulseBridge program perturbations.** **a,** Initial cells are shifted along RNA directions defined by the DKK1/GAS5, B2M/HLA, EREG/PCNA and SAT1/LGALS1/mitochondrial programs, and the modified states are simulated to day 3. **b,** Decreasing or increasing the candidate adaptive program changes the predicted balance between this program and the combined anti-tumor-like response. **c,** Each program shift (bar) is compared with 24 random shifts of the same norm applied to the same starting cells (gray points).

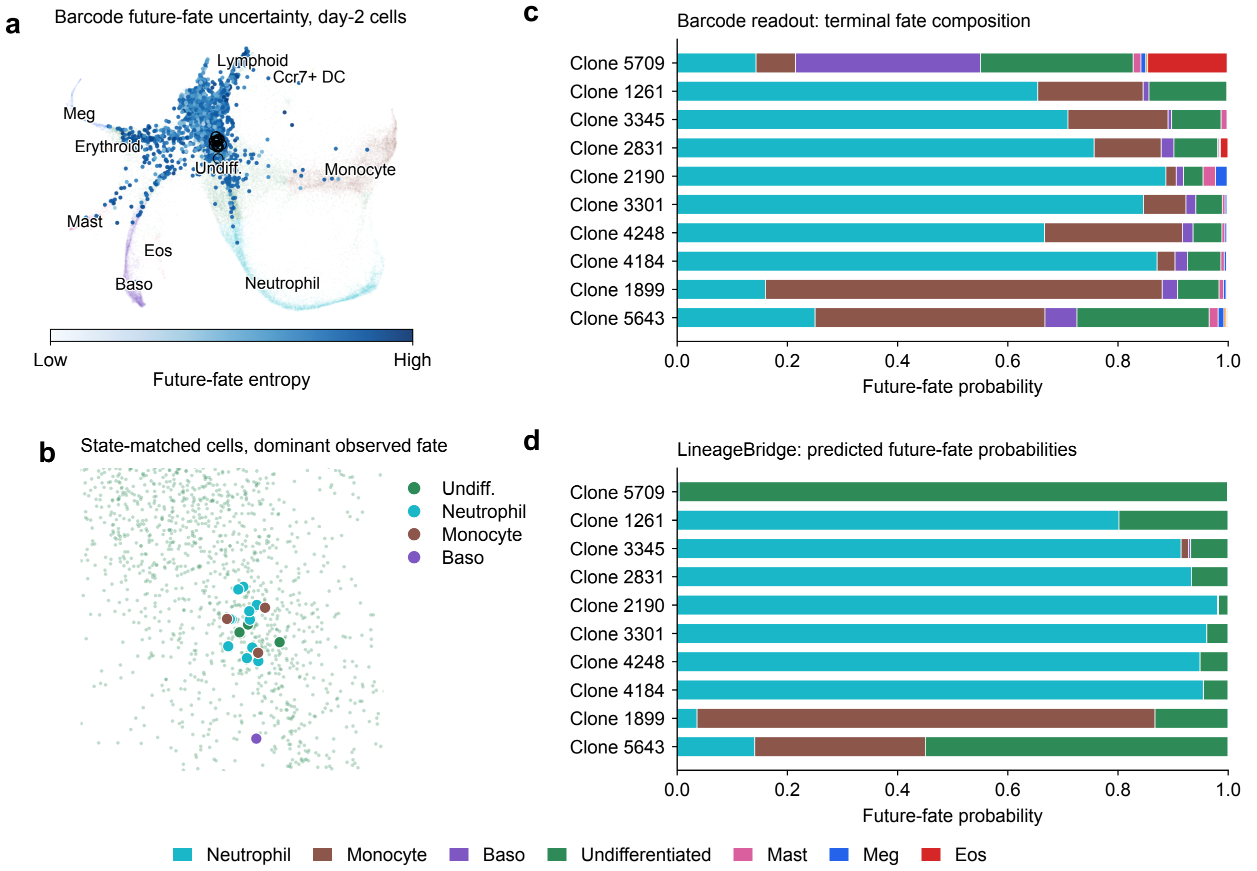

Supplementary Fig. 8: **State-matched early progenitors can have different clonal futures. a,** Barcode-derived future-fate uncertainty on the day-2 LARRY RNA map. **b,** Eighteen barcode-labeled cells selected from a compact region of the early RNA state space. **c,** Observed terminal fate composition of selected clones from this region. **d,** LineageBridge-predicted fate probabilities for the corresponding early cells.

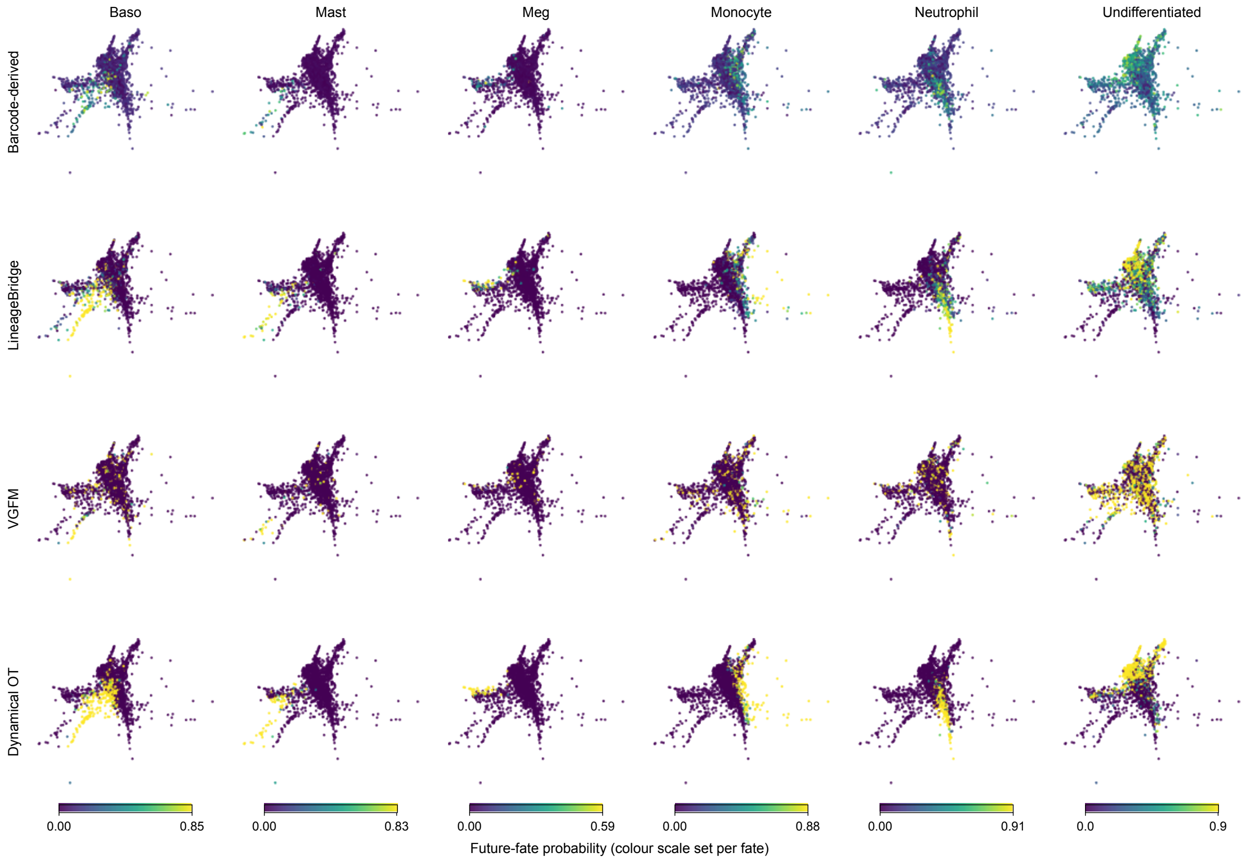

Supplementary Fig. 9: **Clone-supported future-fate predictions across selected hematopoietic lineages.** Columns correspond to basophil, mast-cell, megakaryocytic, monocytic, neutrophilic and undifferentiated outcomes. The top row shows barcode-derived future-fate probabilities for 1,728 eligible day-2 cells on the SPRING map [22]. The following rows show LineageBridge, VGFM and dynamical OT predictions for the same cells. Color scales are set separately for each fate, as indicated by the bars below each column.

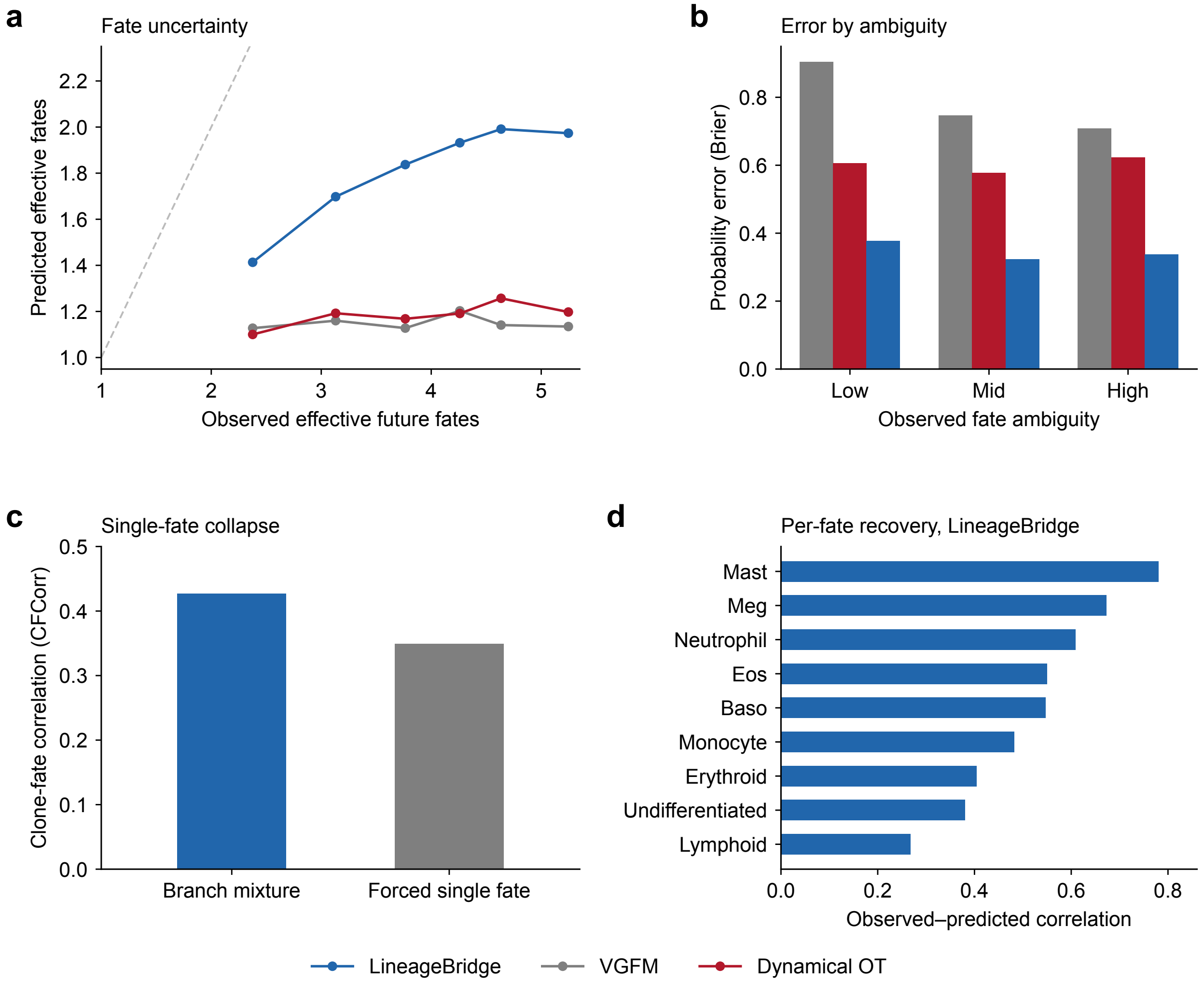

Supplementary Fig. 10: **Evaluation of fate uncertainty in LineageBridge. a,** Predicted and observed effective numbers of future fates, calculated as the exponential of fate entropy. Cells are grouped into six bins by barcode-derived fate entropy, and points show clone-reliability-weighted means. The dashed line indicates equal predicted and observed values. **b,** Brier error for cells with low, intermediate and high barcode-derived fate entropy, divided at the tertiles. **c,** Clone-fate correlation for the full LineageBridge fate distribution and after assigning each cell entirely to its most probable fate. **d,** Clone-reliability-weighted correlations between predicted and barcode-derived probabilities for the displayed fates.

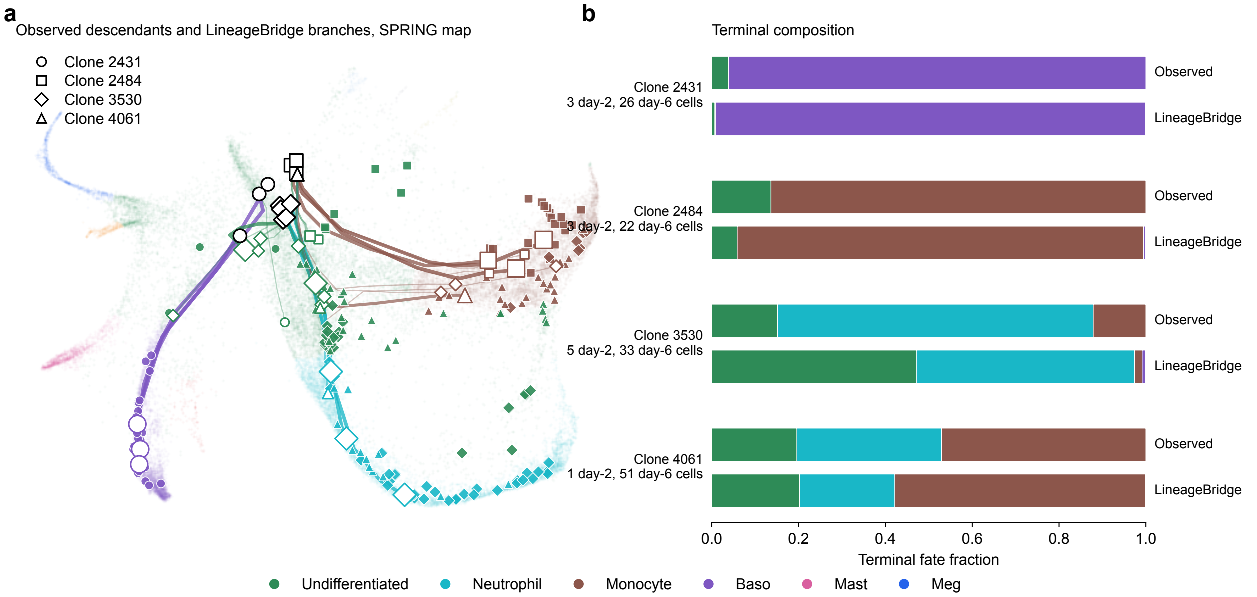

Supplementary Fig. 11: **Representative LARRY clone trajectories and terminal fates. a,** Observed cells from four clones and their predicted branches are shown on the SPRING map. Shapes identify clones and colors identify cell types. Black-outlined open symbols mark day-2 cells, filled symbols mark observed descendants, and colored open symbols mark predicted endpoints. Lines show predicted branches, with greater width indicating greater branch weight. **b,** Observed and predicted terminal fate fractions for each clone. Labels give the numbers of observed cells at days 2 and 6.

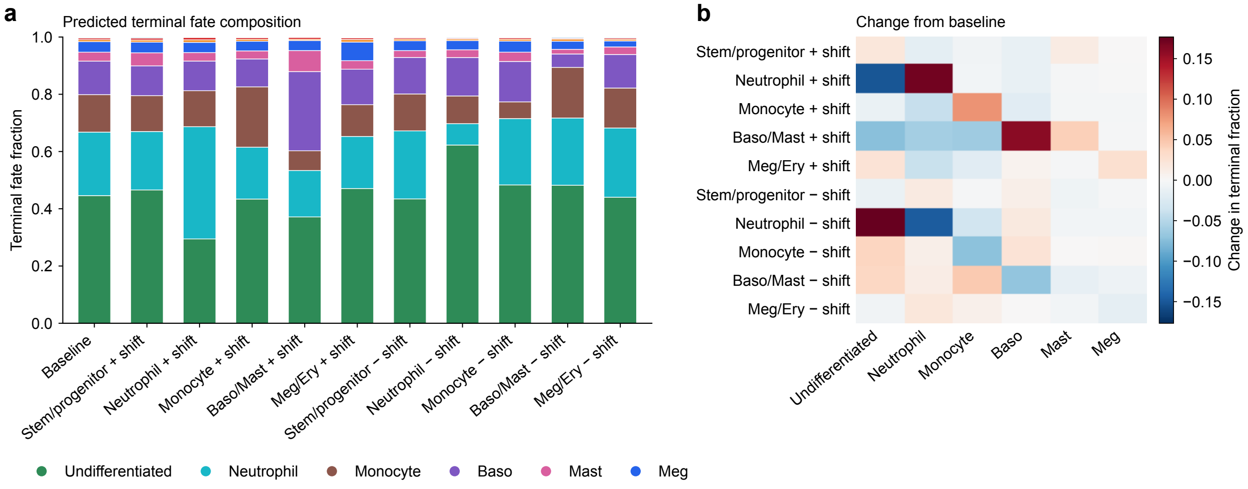

Supplementary Fig. 12: **LineageBridge program perturbations redirect predicted terminal fates.** The initial RNA states of 360 day-2 cells were shifted by +5 or −5 times the mean principal-component loading vector of the genes in each program: stem/progenitor, neutrophil, monocyte, basophil/mast-cell and megakaryocyte/erythroid. The modified and unmodified cells were then propagated through the same fitted LineageBridge dynamics. **a,** Stacked bars show the mean terminal fractions assigned to undifferentiated, neutrophil, monocyte, basophil, mast-cell and megakaryocytic fates. **b,** Change in each terminal fate fraction relative to the unmodified prediction, for every perturbation.

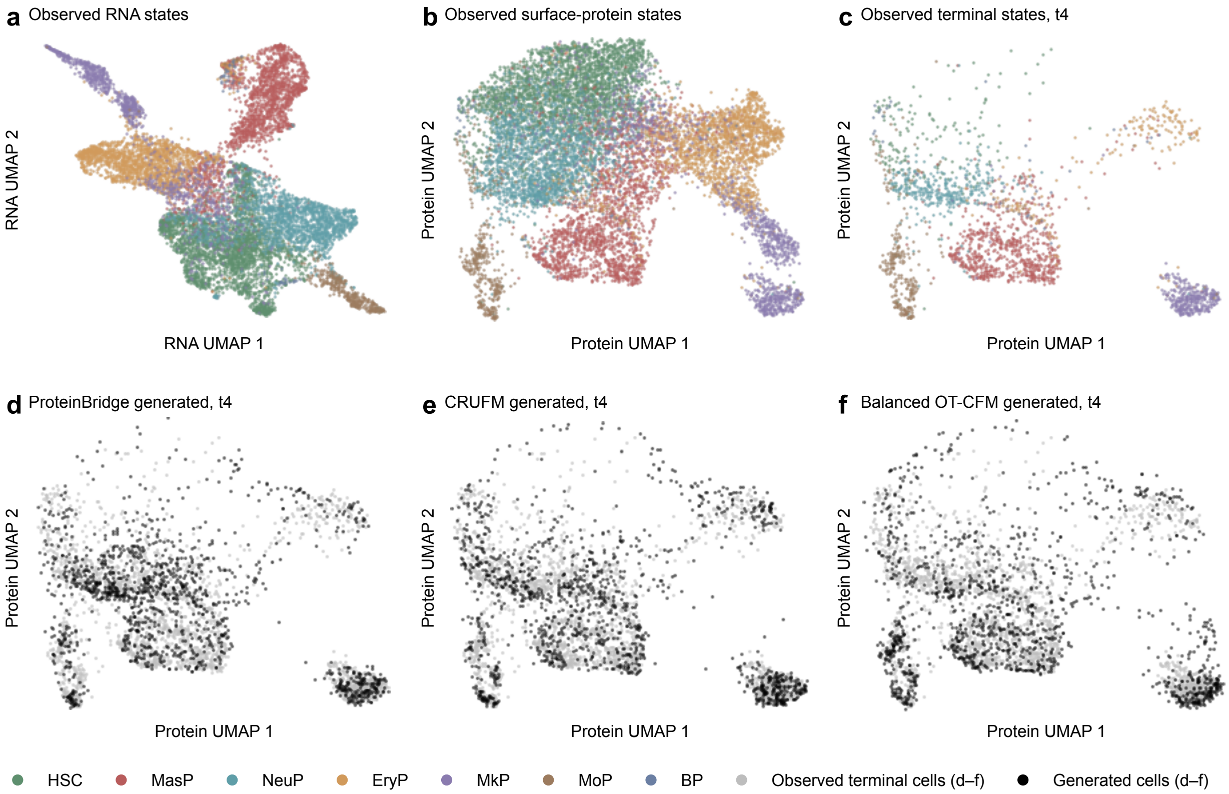

Supplementary Fig. 13: **Generated surface-protein distributions compared on the observed CITE-seq manifold. a,** Observed RNA states and **b,** observed surface-protein states from the paired CITE-seq measurements, colored by annotated phenotype. **c,** Observed surface-protein states at the terminal time. **d–f,** Terminal surface-protein states generated by ProteinBridge, CRUFM and balanced OT-CFM, respectively, projected onto the same observed surface-protein map. Gray points show observed terminal cells and black points show generated cells. Phenotype colors are used only in **a**–**c**.

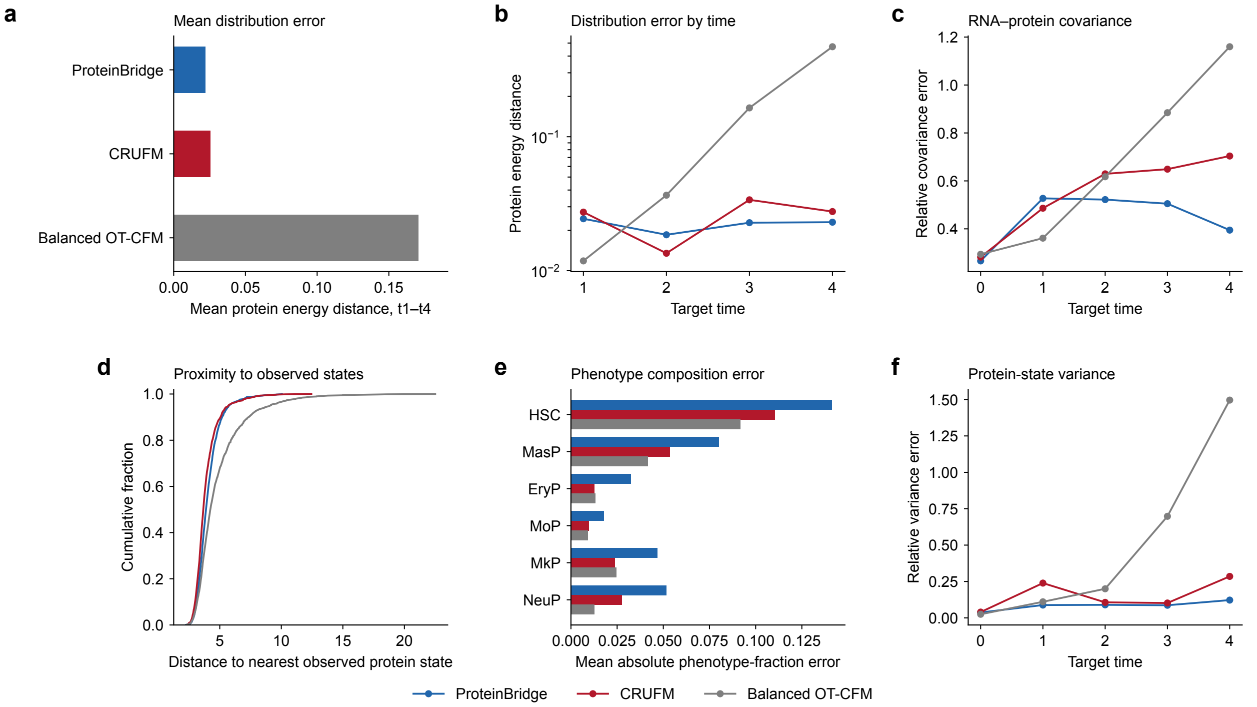

Supplementary Fig. 14: **Quantitative checks of generated surface-protein distributions. a,** Mean protein energy distance across the four predicted times for ProteinBridge, CRUFM and balanced OT-CFM. **b,** Protein energy distance at each predicted time. **c,** Relative RNA–protein cross-covariance error across future times. **d,** Cumulative distribution of the distance from each generated surface-protein state to its nearest observed surface-protein state. **e,** Mean absolute error of the predicted phenotype fractions for the five descendant surface-protein phenotypes and the retained HSC-like category. **f,** Relative error in surface-protein-state variance.

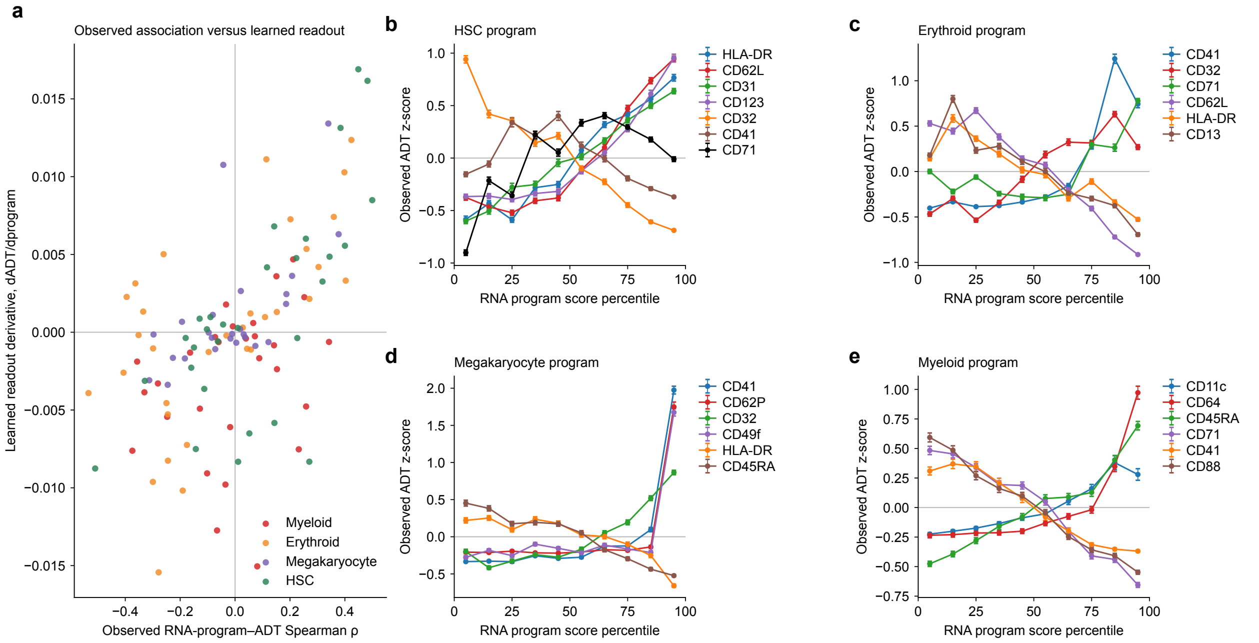

Supplementary Fig. 15: **Validation of RNA-to-surface-protein readouts used for ProteinBridge interpretation. a,** Spearman correlations between observed RNA-program scores and ADT levels are compared with derivatives of the fitted surface-protein readout for selected marker pairs. Points are colored by RNA program. **b–e,** Observed ADT z-scores (mean ± s.e.m.) across percentile bins of the HSC, erythroid, megakaryocyte and myeloid RNA program scores for selected markers.

### References

[1] Marco Cuturi. Sinkhorn distances: Lightspeed computation of optimal transport. In *Advances in Neural Information Processing Systems*, volume 26, 2013.

[8] Zhenyi Zhang, Tiejun Li, and Peijie Zhou. Learning stochastic dynamics from snapshots through regularized unbalanced optimal transport. In *The Thirteenth International Conference on Learning Representations*, 2025.

[9] Zhenyi Zhang, Zihan Wang, Yuhao Sun, Jiantao Shen, Qiangwei Peng, Tiejun Li, and Peijie Zhou. Deciphering cell-fate trajectories using spatiotemporal single-cell transcriptomic data. *npj Systems Biology and Applications*, 12:2, 2026.

[10] Yaron Lipman, Ricky T. Q. Chen, Heli Ben-Hamu, Maximilian Nickel, and Matt Le. Flow matching for generative modeling. In *The Eleventh International Conference on Learning Representations*, 2023.

[11] Alexander Tong, Kilian Fatras, Nikolay Malkin, Guillaume Huguet, Yanlei Zhang, Jarrid Rector-Brooks, Guy Wolf, and Yoshua Bengio. Improving and generalizing flow-based generative models with minibatch optimal transport. *Transactions on Machine Learning Research*, 2024.

[12] Dongyi Wang, Yuanwei Jiang, Zhenyi Zhang, Xiang Gu, Peijie Zhou, and Jian Sun. Joint velocity-growth flow matching for single-cell dynamics modeling. In *Advances in Neural Information Processing Systems*, volume 38, 2025.

[13] Qiangwei Peng, Zihan Wang, Junda Ying, Yuhao Sun, Qing Nie, Lei Zhang, Tiejun Li, and Peijie Zhou. WFR-FM: Simulation-free dynamic unbalanced optimal transport. In *The Fourteenth International Conference on Learning Representations*, 2026.
